# How Error Correction Affects PCR Deduplication: A Survey Based on UMI Datasets of Short Reads

**DOI:** 10.1101/2024.05.30.596723

**Authors:** Pengyao Ping, Tian Lan, Shuquan Su, Wei Liu, Jinyan Li

## Abstract

Next-Generation Sequencing (NGS) data is widely utilised for various downstream applications in bioinformatics, and numerous techniques have been developed for *PCR-deduplication* and *error-correction* to eliminate bias and errors introduced during the sequencing. This study first-time provides a joint overview of recent advances in PCR-deduplication and error-correction on short reads. In particular, we utilise UMI-based PCR-deduplication strategies and sequencing data to assess the performance of the solely-computational PCR-deduplication approaches and investigate how error correction affects the performance of PCR-deduplication. Our survey and comparative analysis reveal that the deduplicated reads generated by the solely-computational PCR-deduplication and error-correction methods exhibit substantial differences and divergence from the sets of reads obtained by the UMI-based deduplication methods. The existing solely-computational PCR-deduplication and error-correction tools can eliminate some errors but still leave hundreds of thousands of erroneous reads uncorrected. All the error-correction approaches raise thousands or more new sequences after correction which do not have any benefit to the PCR-deduplication process. Upon these discoveries, we offer practical suggestions to enhance the existing computational approaches for improving the quality of short-read sequencing data.

## Introduction

Next-Generation Sequencing (NGS) has revolutionized genomics research, enabling extensive acquisition of genome-wide data with unprecedented speed, precision, and cost-effectiveness and making significant progress in the field of DNA sequencing (DNA-Seq) [1] and RNA sequencing (RNA-seq) [2]. NGS data plays a vital role in various downstream analyses, including estimation of microbial diversity [3], variant calling [4], immune cell responses [5, 6], RNA quantification [7], cancer mutation detection [8], de novo genome assembly [9], de novo transcriptome assembly [10] and nonclinical genotoxicity and carcinogenicity testing [11], detecting allele-specific expression [12], isomiR identification [13] and genome base editing [14]. One of the key steps in NGS is the polymerase chain reaction (PCR) process which is purposely used in the library construction and cluster amplification to increase the number of DNA/RNA molecule fragments. This amplified library or flowcell containing multiple copies of each original molecule fragment is helpful for a reliable sequencing of these molecule fragments into digital reads. Such identical reads from the replicates of one molecule fragment are referred to as the PCR duplicates of the read of the original molecule fragment [15, 16].

However, PCR has biases or preferences in the amplification of certain types of molecule fragments, meaning that the duplicate number of a molecule fragment can be quite different from others with a varying amplification rate. Sometimes, PCR also generates new hybrid molecules; and sometimes PCR generates an unprecise replicate of a molecule. These mistaken nucleotide bases in an unprecise PCR replicate of a molecule are called PCR errors. PCR errors that happen at early PCR cycles are usually inherited or propagated into subsequent cycles, making the error distribution complicated [17]. These uneven amplification rates (biases) and the complicated PCR error distributions confuse many NGS downstream applications such as estimation of microbial diversity and composition [3], variant calling [4], immune cell heterogeneity understanding [5, 6], RNA quantification [7], cancer mutation detection [8].

To this end, numerous biological and computational methods for PCR-deduplication have been proposed over the past years. PCR-deduplication identifies and removes duplicated reads in a fasta/fastq file, leaving only unique reads as true biological sequences for downstream analysis. Considering unprecise duplicates are also generated in the PCR process, most of these PCR-deduplication methods identify the base errors and remove these erroneous reads to keep only those unique reads close to the ground truth as much as possible. The PCR-deduplication process involves sequence comparisons of expensive complexity to find exact and unprecise duplicates for retaining only the unique copies of the reads [18, 19, 20, 21].

In addition to these PCR biases and errors, NGS also makes errors during other steps such as at the sample handling step or the base calling step (due to fluorophore crosstalk) [22, 23]. The sequencing errors stemmed from these sources add more confusion to the accuracy of downstream data analysis, especially in de novo genome assembly [9], de novo transcriptome assembly [10] and nonclinical genotoxicity and carcinogenicity testing [11]. Therefore, it is crucial to overcome all of these PCR and other steps’ sequencing biases and errors whenever NGS data is involved to ensure the reliability and accuracy of downstream analyses [5, 24].

Error-correction next-generation sequencing technology is defined as the method that identifies erroneous bases no matter caused by PCR or by fluorophore crosstalk as many as possible in a fasta/fastq dataset and then turns these erroneous bases into their correct formats.

PCR-deduplication makes the size of the original sequencing data (fasta or fastq files) much smaller as it aims to maintain only the unique genuine biological reads, while error-correction never changes the number of reads in the original sequencing data—it just identifies erroneous reads in the dataset and makes corrections on them. Although PCR-deduplication and error-correction methods employ different strategies to achieve different goals, both of them have a common technical component to eliminate sequencing uncertainty such as biases and errors. Recent PCR-deduplication algorithms [25, 26] also consider fluorophore crosstalk errors during the deduplication process. Some deduplicated results for example by Calib [27] and DAUMI [26] can be restored to their original data using the original sequencing IDs stored in the Calib description section or using the read abundance levels stored in the DAUMI description section of the deduplicated read set files. Figure 1 is a schematic diagram depicting an overview of molecular tracking, PCR-deduplication and error-correction processes. It is important to recognise that some PCR-deduplication tools incorporate error correction processes. Therefore, in this study, the term “error correction” encompasses the correction procedures employed in both deduplication and error correction methods. Meanwhile, “error-correction” specifically denotes methods solely focused on error-correction techniques for sequencing data, which identify erroneous reads in the dataset and make corrections to them.

**Figure 1:**
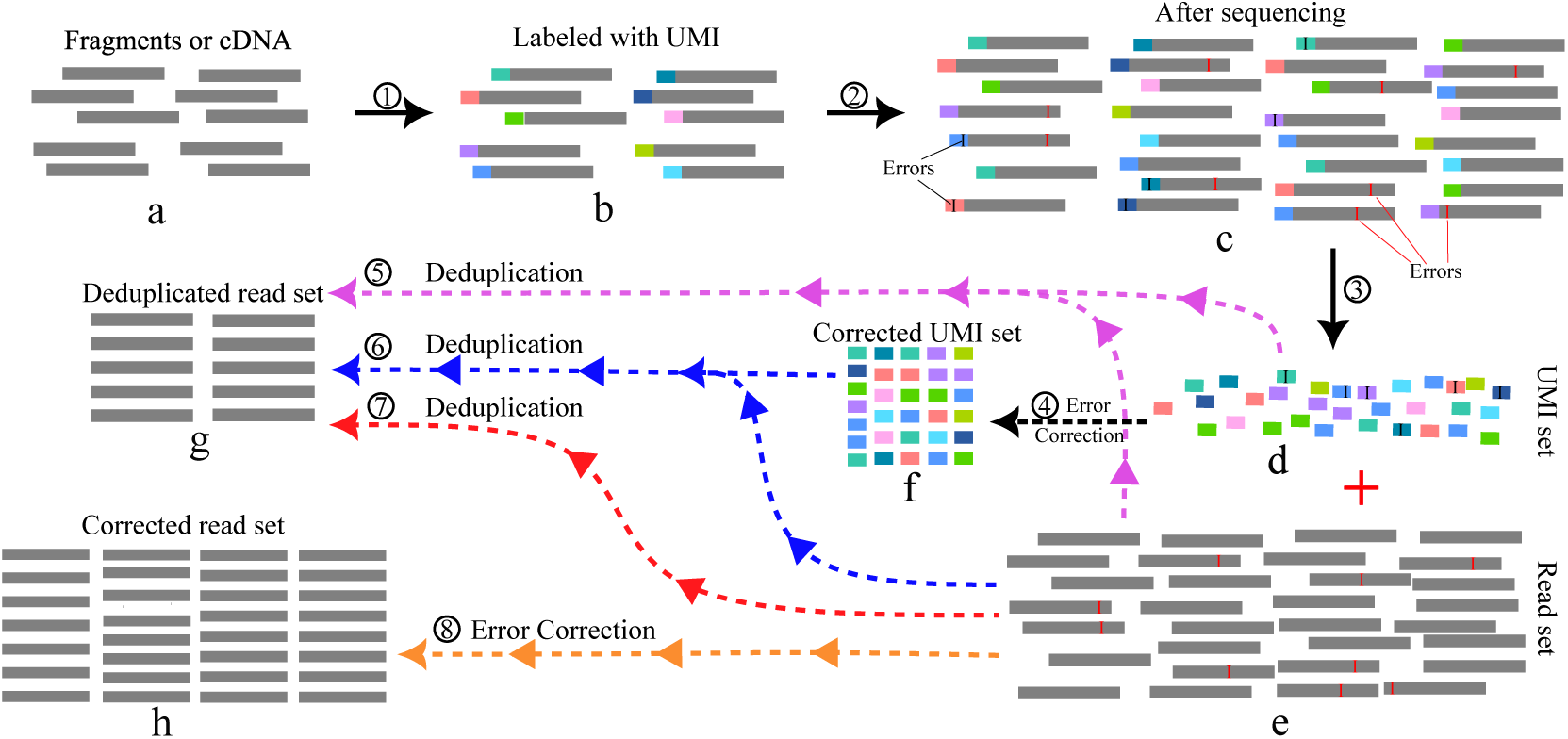
Schematic overview of molecular tracking, PCR-deduplication and error-correction processes. (a-c) illustrate the utilization of unique molecular identifiers (UMIs) for precise molecule tracking, accounting for errors introduced during sequencing, where errors may manifest in both UMI sequences and read sequences. (d-g) describe the PCR-deduplication process: ➄ signifies the removal of duplicated reads using UMI tags, while ➃-f-➅ depicts the elimination of duplicates with corrected UMI tags. ➆ represents duplicate read removal using only computational methods. (e-➇-h) elucidate the computational error-correction process aimed at eliminating all the errors introduced during sequencing.

To our knowledge, there is no joint investigation of PCR-deduplication and error-correction, nor research work assessing the impact of error correction on PCR-deduplication so far. Instead, most of the methods are only for read duplicate removal or separately for error correction assessment on simulated sequencing datasets. In fact, evaluating the performance of these algorithms is challenging due to the absence of ground truth in the sequencing fragments. Utilising unique molecular identifiers (UMIs) provides one of the most effective strategies for recording and tracking sequencing errors during the PCR sequencing process [5, 24]. In particular, error-correction methods have been extensively evaluated in the literature using constructed UMI-based ground truth datasets [28].

This study briefly reviews current methodologies for PCR-deduplication and error-correction and performs a comparative analysis from novel perspectives using UMI short-read sequencing datasets. An ideal algorithm for alleviating uncertainties from the PCR and fluorophore crosstalk biases and errors should achieve two key objectives: (1) identifying and preserving the authentic biological sequences and (2) maintaining precise read abundance levels without introducing new errors or sequences; therefore, this study

- compares the differences of deduplicated sets of reads generated by different UMI-based PCR-deduplication methods;
- utilises UMI-based PCR-deduplication methods to assess the performance of solely-computational PCR-deduplication tools;
- examines the percentage of corrected reads and the number of wrongly introduced new sequences after the error-correction process;
- investigates the impact of error correction procedures of PCR-deduplication and error-correction methods on deduplication for short reads.

### PCR-deduplication for Short Reads: a Brief Review

Existing PCR-deduplication methods can be categorised into two categories. The main idea of the first category is to utilise biochemical techniques such as unique molecular identifiers (UMIs) to accurately count and track precise and unprecise molecule fragments. While the second category takes only computational steps for the PCR-deduplication. Both the UMI and the solely-computational methods can be further classified according to whether they use reference genome sequences. These methods are summarised in Table 1 with more details.

**Table 1:**
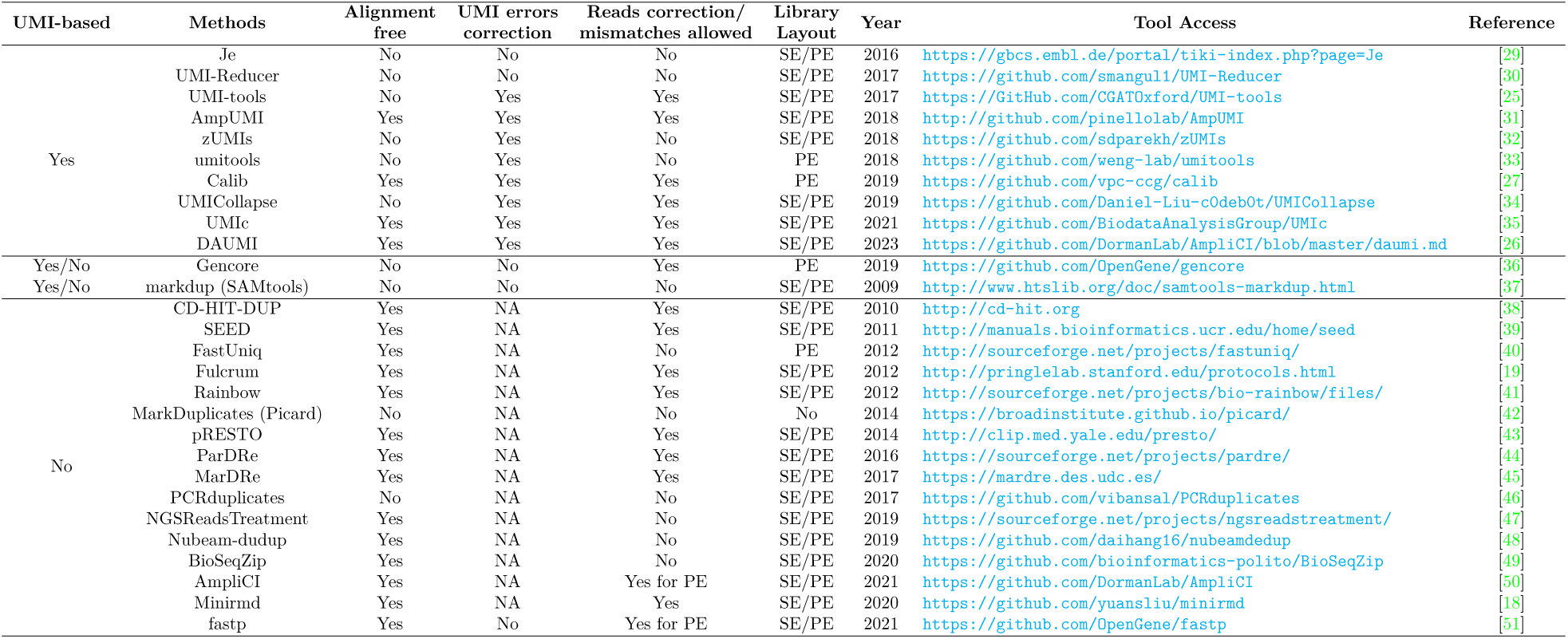
Summary of the PCR-deduplication approaches with and without use of Unique Molecular Identifiers (UMI) data.

### PCR-deduplication methods working on UMI datasets

During the library preparation step in NGS, UMIs are added into the library to attach to a specific location for every DNA fragment before PCR amplification. In ideal cases, each unique molecular fragment corresponds to a UMI. Therefore, unprecise duplicates of a molecular fragment after PCR amplification can be recognised through their UMI sequences when there exist no PCR errors in the UMI sequences.

Alignment-based PCR-deduplication methods distinguish duplicated reads by grouping the reads using mapping positions and identical UMIs under the assumption that there exist no PCR errors in the UMI sequences. The typical process of these strategies clusters the reads with mapping positions first and then regroups reads in the same cluster based on UMIs. For instance, Girardot et al. developed a suite of tools named Je – an extension of the popular PCR-deduplication tool Picard’s MarkDuplicates[42] by supporting more accurate duplicate removal. Je identifies duplicates based on mapping positions to the reference and keeps the reads with distinct UMIs for downstream processing [29]. Similarly, UMI-Reducer identifies and collapses PCR duplicates using identical UMI sequences and the mapping location of the read. It also treats these reads that have the same mapping positions but different UMIs as unique reads [30]. Gencore groups the mapped sequences and generates consensus sequences by merging these reads within each cluster. This process can clear PCR biases and errors during the generation of consensus reads. When working with sequencing data that includes UMI tags, Gencore could cluster reads based on the UMIs for the identification of reads originating from the same fragments [36].

Unlike those tools which did not consider the tiny amount of UMI errors made by the PCR amplification for PCR-deduplication, Smith et al. reported that mistaken bases in UMIs are common after PCR [25], and then they developed a network-based method, UMI-tools, to correct UMI errors before PCR-deduplication. In the correction process, UMI-tools constructs a UMI network where two UMIs are linked if they have a one-base difference; then, the method uses three techniques to determine the count of distinct molecules at a particular genomic position. The main goal of these techniques is to simplify the network by identifying a representative UMI that represents the entire network rather than identifying the exact sequence of the original UMIs [25]. Additionally, zUMIs, a method designed to process multiplexed RNA-seq data that contain known and random BCs and UMIs, employs the strategy proposed in UMI-tools to handle UMI errors when necessary. The pipeline removes reads with low-quality BCs and UMIs and then it maps the remaining reads to a reference genome [32].

Not relying on a reference genome, alignment-free methods do not map reads to a reference during PCR-deduplication. This approach avoids issues such as clustering reads that have multiple mapping locations and it still works when no references are available for mapping. Alignment-free methods typically retain the reads with the highest UMI counts and quality scores but sometimes they vary these parameters [34, 35]. For example, Calib conducts duplicate removal by constructing a graph based on UMI and sequence similarity through hashing and MinHashing techniques. In the constructed graph, each connected subgraph is identified as a cluster. One limitation is that Calib can only handle clustering paired-end reads with UMIs produced from Illumina platforms [27]. UMIc rectifies base errors in UMI sequences by leveraging base frequency and quality information. The correction process is initially implemented for reads sharing identical UMI tags. Subsequently, UMIc merges UMIs based on both UMI-to-UMI distances and sequence-to-sequence distances. Finally, it addresses errors in sequences belonging to the same group of merged UMIs [35]. AmpUMI performs sequencing error correction during the PCR-deduplication process. In essence, this method involves clustering sequences based on UMIs and retaining the most frequently occurring sequences as biological reads. The remaining sequences are then discarded as they are considered to be attributable to sequencing errors [31].

Moreover, Fu et al. introduced experimental protocols and computing approaches by integrating UMIs into standard RNA-seq procedures to determine PCR copies in RNA-seq and small-RNA sequencing data. Their study demonstrated a superior accuracy of duplicate removal when employing UMI sequence data for comparison with those without UMIs. Notably, without the use of UMIs, numerous biologically meaningful reads are mistakenly flagged as duplicates during the removal process [33]. Recently, a probabilistic algorithm, DAUMI [26], was designed for amplicon sequencing to identify actual biological sequences and estimate the read abundance by eliminating PCR and fluorophore crosstalk biases and errors in the UMIs and reads. One advantage of DAUMI is that it can deal with UMI collision, even on highly similar sequences.

### PCR-deduplication approaches without use of UMI data

Without using UMI tags, most of the solely-computational PCR-deduplication methods aim to reduce time complexity and computational memory usage for removing precise duplicate or unprecise, nearly-duplicate reads. Their key step is to develop parallel computing strategies and advanced data structures for enhancing PCR-deduplication efficiency.

These computational methods, including markdup in SAMtools [37, 52], MarkDuplicates in picard [42], PCRduplicates [46] and Gencore [36], also depend on the alignment data of the reads to reference genomes helping the removal of the duplicate reads. These methods have experienced a loss of accuracy when reads have multiple mapping locations and are rendered ineffective in scenarios where no references are available for mapping.

Alignment-free methods employ advanced data structures to recognise duplicated reads by comparing the differences in a time- and memory-efficient manner. For instance, NGSReadsTreatment [47] was designed to remove redundancies from the raw reads with the Cuckoo Filter, an enhancement of the Bloom Filter, using cuckoo hashing to prevent collisions. During the redundancy removal, NGSReadsTreatment keeps a read if its fingerprint is not found in the cuckoo hash table; otherwise, discard. Nubeam-dudup [48] employs unique numbers to represent reads via matrices product calculation by representing nucleotides as matrices and then using the unique numbers instead of reads themselves for pairwise comparison to identify duplicated reads. These methods only keep existing identical reads without considering PCR bias and error elimination for PCR-deduplication. In addition, most existing methods split the whole data set into chunks and perform pairwise comparisons in each cluster to reduce the time complexity of the pairwise comparison of all the reads. For instance, Fulcrum [19], ParDRe [44] and MarDRe [45] have employed a prefix-clustering technique, while still exploiting a greedy incremental clustering algorithm. Minirmd [18] clusters reads using minimiser for multiple rounds to remove duplicates and near-duplicates.

Moreover, some tools allow mismatches and perform read correction in the awareness of PCR bias and error removal. For instance, CD-HIT-DUP [38], Rainbow [41], pRESTO [43], ParDRe [44], MarDRe [45] and Minirmd [18] allow a user-defined number of mismatches while SEED [39] and Fulcrum [19] allow up to three mismatches during removing duplicated reads. Eliminating PCR biases and errors while identifying duplicated reads is more complex than the pure PCR-deduplication. These approaches with tolerance to small amounts of mismatches can correct some PCR errors and remove near-duplicate reads. However, these methods may only discover some expected base errors since they conduct an approximate pairwise comparison across the whole reads.

### Error-correction Methods: A Brief Review

Error-correction algorithms use various statistical/mathematical ideas to detect and correct errors to improve the quality of sequencing data. To our knowledge, existing error-correction methods are all computational approaches without using UMI information. These error-correction methods in Table 2 can be classified into *k*-mer-based, multiple sequence alignment (MSA) based algorithms and de Bruijn graph (DBG) based methods. In addition to these general-purpose methods, several other error-correction methods were designed for specific sequencing tasks, which we classified as Scenario-based error-correction.

**Table 2:**
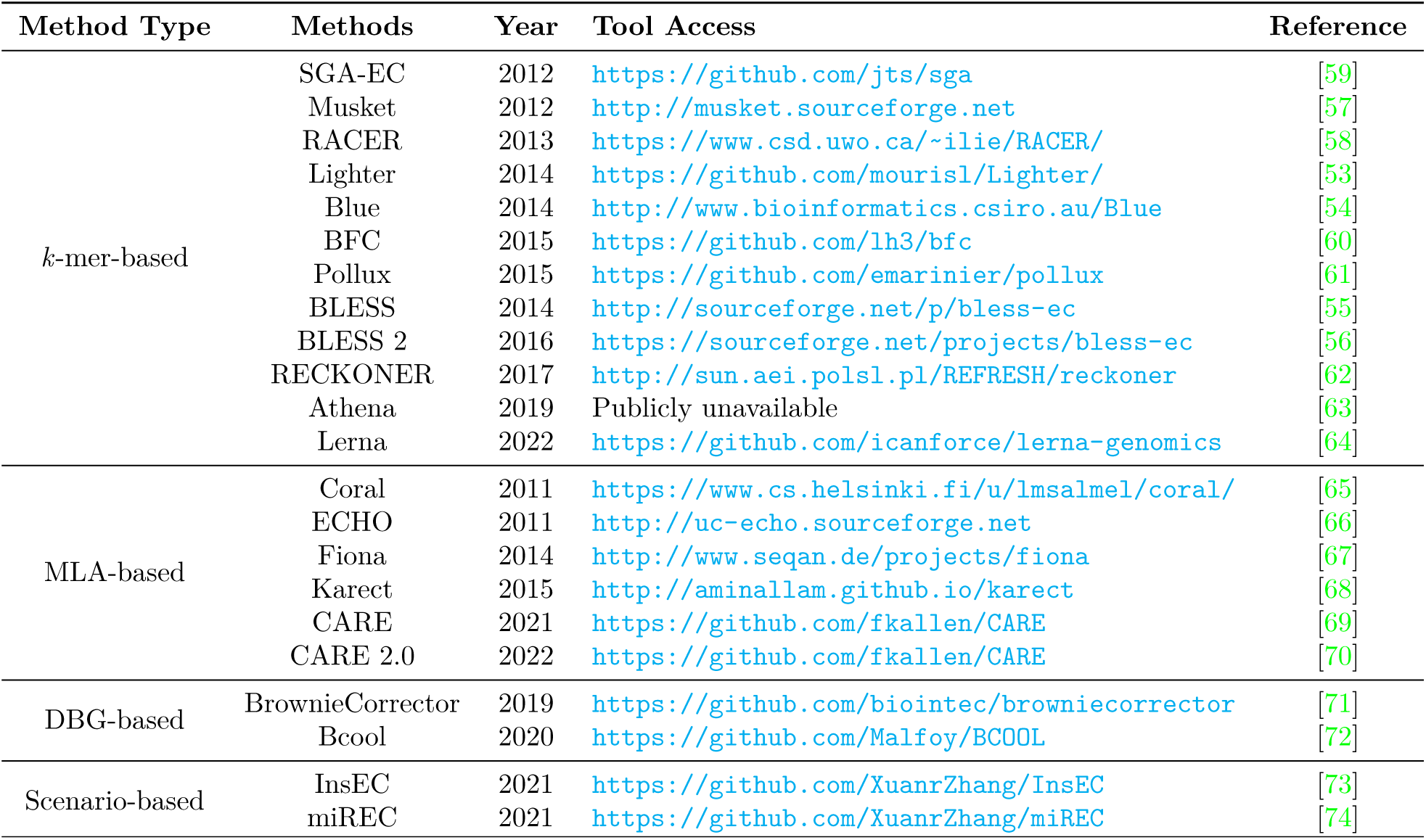
Summary of the computational error-correction methods.

### *K*-mer-based methods

*K*-mer is a sub-string of a sequence; given a frequency threshold, *k*-mer-based algorithms aim to determinate high-frequency *k*-mers, namely solid *k*-mers, otherwise marked as weak *k*-mers. Then, these algorithms replace weak *k*-mers with matching solid *k*-mers. Some of them were specifically designed to correct substitution errors. For instance, Lighter [53] and Blue [54] employ a *k*-mer-based strategy and quality scores to target substitution error correction. BLESS [55] and its successor, BLESS 2 [56], concentrate on rectifying substitution errors in sequencing reads through *k*-mer analysis-based Bloom filters. BLESS 2, in particular, demonstrates enhanced performance in both speed and accuracy, especially when handling larger datasets, compared to the original BLESS. Musket [57] and RACER [58] primarily address substitution errors. In contrast, SGA-EC [59], BFC [60], Pollux [61], and RECKONER [62] were designed to tackle various types of errors, including substitutions, insertions, and deletions.

In addition, Athena[63] and Lerna[64] were proposed using deep learning frameworks such as Recurrent neural network (RNN) and attention-based transformers to automatically find the optimal *k*-mer size to improve the performance of the *k*-mer based methods. However, these two tools’ source codes or software are not publicly available.

### MSA-based methods

MSA-based methods utilise the multiple sequence alignment information, including base coverage and erroneous positions, to group the reads with shared knowledge and generate a contig as an error-free reference to correct false bases. Earlier methods were represented by Coral [65], ECHO [66], Fiona [67], and Karect [68]. MSA-based approaches also incorporate other strategies rather than not solely utilising the MLA technique. For instance, Karect treats each read as a reference to construct an initial partial-order graph and then conducts multiple alignments for an optimized group of sequences similar to the reference read to update the partial-order graph. Next, it utilises the accumulated partial alignment results to rectify the reference read. In addition, CARE [69] implemented in 2021 aims to produce fewer false-positive corrections while achieving more true positives by searching similarity within extensive read collections based on mini hashing. Then, CARE 2.0 [70], an extension of CARE, was developed using the machine learning method of random forests to increase the correction precision and sensitivity further. One common drawback of these MSA-based algorithms is that constructing optimal MSAs is of high computational complexity and time-consumption.

Moreover, the performance of some methods, such as those by Fiona [67] and RACER [58], may be influenced by the preset parameters (e.g., the estimated genome size) because it is tricky for users or it is not suitable to estimate the genome size used for non-whole-genome sequencing data (e.g., synthetic sequencing).

### DBG-based methods

BrownieCorrector [71] and Bcool [72] rely on constructing de Bruijn graph (DBG) to detect and rectify errors in reads. Both of them utilised *k*-mer-based techniques for constructing reference DBG. Then erroneous nodes are detected and cleaned from the reference DBG based on coverage and graph topology. Finally, the error reads are corrected to their normal state by aligning them to the corrected DBG. These two methods adopt the same data structures but differ in implementation details, and BrownieCorrector is applicable to only paired-end sequencing datasets.

### Scenario-based error-correction

The non-uniform sequencing depths restrict the above methods’ effectiveness for error removal in specific sequencing scenarios. For example, when analysing single nucleotide polymorphism (SNP), particular genes or pathways involved in the disease mechanism or one specific genome segment would be more important than the other loci in the genome.

General-purpose correction methods may under-correct or over-correct the sequencing errors in short reads of disease-associated genes when targeted gene sequencing is applied. An instance-based algorithm, InsEC[73], was proposed to rectify the errors in the reads of a disease-related gene using local sequence characteristics and statistics directly correlated to these genes. Focusing on correcting miRNA sequencing errors, an error rectification method, miREC[74], was proposed using a lattice structure combining *k*-mers, (k-1)-mers and (k+1)-mers to maintain the frequency differences of the *k*-mers. Notably, InsEC and miREC have the same advantage of not introducing new errors when performing error corrections. In addition, there are other scenarios for correcting errors in short reads. For example, to eliminate the UMI errors introduced during PCR and nucleotide miscalling and indels during sequencing, the process of eliminating UMI errors was introduced by UMI-tools[25] and DAUMI [26], respectively, during PCR-deduplication.

### How Error Correction Affects PCR Deduplication: Performance Comparison on UMI Datasets

We took 11 UMI-based single-end sequencing datasets of the accession IDs [SRR1543964](https://www.ncbi.nlm.nih.gov/Traces/study/?acc=SRP045430&o=acc_s%3Aa) - [SRR1543971](https://www.ncbi.nlm.nih.gov/Traces/study/?acc=SRP045430&o=acc_s%3Aa), SRR28313972, SRR28313990, and SRR28314008 together with a pair-end sequencing dataset of the accession ID SRR11207257 for the comparative analysis. These datasets [SRR1543964](https://www.ncbi.nlm.nih.gov/Traces/study/?acc=SRP045430&o=acc_s%3Aa) - [SRR1543971](https://www.ncbi.nlm.nih.gov/Traces/study/?acc=SRP045430&o=acc_s%3Aa) and SRR11207257 were previously employed in the benchmark study conducted by Mitchell et al [28]. For those datasets, the first twelve bases of each sequence represent the sequence of the UMI, while the subsequent twelve bases serve as an index for multiplexing. We extracted the sequences of these UMIs and stored them in the description section of each record of the sequencing data. The multiplexing tags were removed using the fastp tool [51]. The group of datasets SRR28313972, SRR28313990, and SRR28314008 were recently sequenced datasets that published in 2024, and the UMI sequence with length of 11 is stored in the description section.

### Performance by PCR-deduplication methods and their comparison

We focused on four UMI-based PCR-deduplication methods, UMI-tools[25], AmpUMI[31], Calib[27] and UMIc[35], and 9 solely-computational methods NGSReadsTreatment[47], Nubeam-dedup[48], BioSeqZip[49], fastp[51], FastUniq[40], pRESTO[43], CD-HIT-DUP[38], ParDRe[44] and Minirmd[18] for our comparative analysis. The comparative analysis regarding the number changes of the unique reads after PCR-deduplication by these algorithms is summarised in Table 3, and Table 4 lists the number of new sequences introduced by each algorithm after PCR-deduplication.

**Table 3:** Unique reads number comparison after PCR-deduplication by the methods of UMI-tools[25], AmpUMI[31], Calib[27], UMIc[35], NGSReadsTreatment[47], Nubeam-dedup[48], BioSeqZip[49], fastp[51], FastUniq[40], pRESTO[43], CD-HIT-DUP[38], ParDRe[44] and Minirmd[18].

| Dataset | Original | UMI-tools | AmpUMI | Calib | UMIc | NGSReadsTreatment | Nubeam-dedup | BioSeqZip | fastp | FastUniq |
| --- | --- | --- | --- | --- | --- | --- | --- | --- | --- | --- |
| SRR1543964 | 312070 | 38589 | 50458 | / | 36998 | 312070 | 312070 | 312070 | 312065 | / |
| SRR1543965 | 273707 | 35439 | 48156 | / | 31629 | 273707 | 273707 | 273707 | 273698 | / |
| SRR1543966 | 305104 | 30498 | 52157 | / | 31961 | 305104 | 305104 | 305104 | 305097 | / |
| SRR1543967 | 336566 | 36718 | 57444 | / | 37253 | 336566 | 336566 | 336566 | 336559 | / |
| SRR1543968 | 277589 | 30517 | 47395 | / | 30688 | 277589 | 277589 | 277589 | 277586 | / |
| SRR1543969 | 375335 | 40227 | 63017 | / | 43897 | 375335 | 375335 | 375335 | 375324 | / |
| SRR1543970 | 290058 | 28620 | 46942 | / | 29538 | 290058 | 290058 | 290058 | 290053 | / |
| SRR1543971 | 182485 | 24245 | 33352 | / | 22995 | 182485 | 182485 | 182485 | 182481 | / |
| SRR28313972 | 5504307 | 5034462 | 2412798 | / | / | 5504307 | 5498421 | 5504307 | 5468005 | / |
| SRR28313990 | 2669507 | 1232133 | 1316953 | / | / | 2669507 | 2669402 | 2669507 | 2640309 | / |
| SRR28314008 | 1696037 | 703947 | 598588 | / | / | 1696037 | 1693794 | 1696037 | 1669773 | / |
| SRR11207257 | 7938261 | / | 421653 | 6401766 | / | 7938261 | 7819073 | 7938261 | 10953857 | 7938261 |
|  | 7660776 | / | 331566 | 6410184 | / | 7660776 | 7482502 | 7660776 | 11962077 | 7660776 |

| Dataset | pRESTO<sup>a</sup> | CD-HIT-DUP<sup>a</sup> | ParDRe<sup>a</sup> | Minirmd<sup>a</sup> |
| --- | --- | --- | --- | --- |
| SRR1543964 | {312069;312070;312070;312070} | {312070;295103;272744;293079} | {312070;229011;197629;180695} | {312070;221440;189201;171298} |
| SRR1543965 | {273705;273706;273706;273706} | {273707;258877;239262;256510} | {273707;201155;174111;160366} | {273707;193622;166483;152038} |
| SRR1543966 | {305104;305104;305104;305104} | {305104;285269;257973;282279} | {305104;225727;196468;180785} | {305104;219728;190308;174061} |
| SRR1543967 | {336566;336566;336566;336566} | {336566;313498;282806;309886} | {336566;244839;212383;195150} | {336566;238483;205400;187461} |
| SRR1543968 | {277587;277589;277589;277589} | {277589;257322;230970;253360} | {277589;198677;169718;153986} | {277589;193068;163597;147266} |
| SRR1543969 | {375335;375335;375335;375335} | {375335;349495;314533;345917} | {375335;272897;232328;209753} | {375335;266857;224950;201107} |
| SRR1543970 | {290057;290057;290058;290058} | {290058;270046;242548;267170} | {290058;209248;179333;163441} | {290058;202770;172885;156358} |
| SRR1543971 | {182484;182485;182485;182485} | {182485;170008;153520;167025} | {182485;133523;116273;107141} | {182485;129762;112676;103228} |
| SRR28313972 | {5484192;5496068;5496072;5496074} | {5504307;5364504;5391771;5313098} | {5504307;5074083;4689876;4451895} | {5504307;5002268;4604781;4349086} |
| SRR28313990 | {2655692;2661654;2661661;2661662} | {2669507;2564944;2575765;2502712} | {2669507;2012208;1371720;1001412} | {2669507;2042730;1401790;1014054} |
| SRR28314008 | {1694962;1695980;1695993;1695994} | {1696037;1559453;1343804;1283796} | {1696037;1166751;972934;880270} | {1696037;1107100;887314;777288} |
| SRR11207257 | {7938261;7938261;7938261;7938261} | {7938261;7938261;7938261;7938261} | {7938261;6390910;5581983;5249419} | {7938261;6548964;5750100;5291977} |
|  | {7656046;7656184;7656243;7656256} | {7660776;6589492;7138851;6549694} | {7660776;5825000;4525245;3957210} | {7660776;5991473;4847956;4242540} |

<sup>a</sup> These methods allow mismatches when PCR-deduplication and their values in {} are obtained by setting mismatch numbers as {0, 1, 2, 3}.
The symbol '/' signifies that this method does not support a single/pair-end data set for PCR-deduplication, or that it cannot produce results when executed on the corresponding data set.

**Table 4:** The number of introduced new sequences after PCR-deduplication by the methods of UMI-tools[25], AmpUMI[31], UMIc[35], NGSReadsTreatment[47], Nubeam-dedup[48], BioSeqZip[49], fastp[51], FastUniq[40], pRESTO[43], CD-HIT-DUP[38], ParDRe[44] and Minirmd[18] on the data sets SRR1543964-SRR1543971 and SRR11207257, respectively.

| Dataset | UMI-tools | AmpUMI | Calib | UMIc | NGSReadsTreatment | Nubeam-dedup | BioSeqZip | fastp | FastUniq | pRESTO<sup>a</sup> | CD-HIT-DUP<sup>a</sup> | ParDRe<sup>a</sup> | Minirmd<sup>a</sup> |
| --- | --- | --- | --- | --- | --- | --- | --- | --- | --- | --- | --- | --- | --- |
| SRR1543964 | 38147 | 0 | / | 6562 | 0 | 0 | 0 | 0 | / | {0} | {0} | {0} | {0} |
| SRR1543965 | 33993 | 2 | / | 3491 | 0 | 0 | 0 | 0 | / | {0} | {0} | {0} | {0} |
| SRR1543966 | 30189 | 1 | / | 2634 | 0 | 0 | 0 | 0 | / | {0} | {0} | {0} | {0} |
| SRR1543967 | 35759 | 0 | / | 3220 | 0 | 0 | 0 | 0 | / | {0} | {0} | {0} | {0} |
| SRR1543968 | 30209 | 0 | / | 2647 | 0 | 0 | 0 | 0 | / | {0} | {0} | {0} | {0} |
| SRR1543969 | 38918 | 1 | / | 8898 | 0 | 0 | 0 | 0 | / | {0} | {0} | {0} | {0} |
| SRR1543970 | 28151 | 0 | / | 2535 | 0 | 0 | 0 | 0 | / | {0} | {0} | {0} | {0} |
| SRR1543971 | 22909 | 1 | / | 2639 | 0 | 0 | 0 | 0 | / | {0} | {0} | {0} | {0} |
| SRR28313972 | 2311229 | 3802 | / | / | 0 | 0 | 0 | 0 | / | {0} | {0} | {0} | {0} |
| SRR28313990 | 143702 | 3993 | / | / | 0 | 0 | 0 | 0 | / | {0} | {0} | {0} | {0} |
| SRR28314008 | 369470 | 0 | / | / | 0 | 0 | 0 | 0 | / | {0} | {0} | {0} | {0} |
| SRR11207257 | / | 421653 | 6401766 | / | 0 | 0 | 0 | 10952692 | 0 | {0} | {0} | {0} | {0} |
|  | / | 331566 | 6410184 | / | 0 | 0 | 0 | 11960912 | 0 | {0} | {0} | {0} | {0} |

<sup>a</sup> These methods permit mismatches during PCR-deduplication, and instances where their values are {0} indicate that these methods never introduce new sequences, achieved by setting mismatch numbers to {0, 1, 2, 3}.
The symbol “/” signifies that this method does not support a single/pair-end data set for PCR-deduplication, or that it cannot produce results when executed on the corresponding data set.

On the 11 single-end datasets, four solely-computational methods NGSReadsTreatment, Nubeam-dedup, BioSeqZip and fastp have made the same or a quite similar unique-read set as those obtained directly from the original data without applying any noise removal ideas (see Table 3). Moreover, none of these methods introduced new sequences after PCR-deduplication as shown in Table 4. These results indicate that these methods identified all the unique reads in the sequencing data as genuine biological sequences.

On the datasets SRR1543964-SRR1543971, three UMI-based PCR-deduplication algorithms (UMI-tools, AmpUMI, and UMIc) resulted in an average decrease (by 88.61%, 82.97%, and 88.69% respectively) in the number of unique reads (see Table 3). Note that UMI-tools and UMIc introduced numerous new sequences that did not exist in the original datasets. Introducing new sequences by UMI-tools is attributed to the reason that it performed the PCR-deduplication and correction of sequencing data based on a reference genome. In detail, approximately 8.24% to 20.27% of the unique sequences after the deduplication were found as new sequences, indicating that UMIc overcorrected PCR biases and errors across the eight datasets.

On the datasets SRR28313972, SRR28313990, and SRR28314008, the number of unique reads decreased by an average of 40.3% and 57.18% after deduplication with UMI-tools and AmpUMI, respectively. Following deduplication, 11.7% to 52.5% of the unique reads obtained with UMI-tools were identified as new sequences. In contrast, only 0.15% and 0.3% of the unique reads were identified as new sequences by AmpUMI in datasets SRR28313972 and SRR28313990, respectively. UMIc was terminated or failed to yield valid deduplication results on these three datasets after running for an extended period of one month.

Furthermore, the solely-computational tools ParDRe and Minirmd allowing mismatches exhibited a gradual decrease in the number of unique reads after PCR-deduplication when the mismatch threshold increased from 0 to 3 (see Table 3), and these methods did not generate any new sequences after PCR-deduplication (see Table 4). The number of unique reads produced by CD-HIT-DUP fluctuates in response to changes in mismatches, but it does not exhibit consistent patterns across all datasets. These performances demonstrate that the mismatch-allowed strategy by these methods can effectively eliminate some PCR biases and errors. Nevertheless, the number of unique reads is still relatively higher than the UMI-based algorithms. Regarding pRESTO, the results as shown in Table 3 and Table 4 suggest that it has failed to identify base errors through the mismatch strategy after PCR-deduplication.

On the pair-end dataset SRR11207257, UMI-tools and UMIc encountered computing memory issues and could not complete the PCR-deduplication process. AmpUMI and Calib, on the other hand, yielded unique reads that consisted entirely of new sequences not present in the original dataset (refer to Table 3 and Table 4). The reason is that these methods utilized both read 1 (R1) and read 2 (R2) from opposite ends of a DNA fragment to reconstruct the complete fragment sequence.

All the solely computational PCR-deduplication methods except fastp did not result in any new sequences on the pair-end dataset SRR11207257 during the PCR-deduplication process. Instead, these methods identified and removed duplicate sequences, resulted in unique reads already present in the original dataset. In contrast, fastp produced a substantial number of new sequences as shown in Table 4, which exceeded the count of unique reads in the original data set as indicated in Table 3. Notably, the new sequence is derived from altering the original sequence during deduplication. The number in Table 3 represents the count of non-repeating sequences rather than the total number. Put differently, following deduplication by fastp, two or more sequences with distinct IDs but identical sequences have been transformed into different sequences. The result suggests that when reconstructing the entire fragment using both read 1 (R1) and read 2 (R2) of a DNA fragment, fastp may tend to overcorrect the sequencing reads.

### Comparison on the numbers of unique reads yielded by UMI-based methods

To compare the deduplicated sets of unique reads yielded by the three UMI-based PCR-deduplication methods AmpUMI, UMI-tools and UMIc, we utilised Venn diagrams to visualise the overlap and differences among the unique read sets after PCR-deduplication. These Venn diagrams are presented in Figure 2 and Supplementary Figure 1 for other datasets.

**Figure 2:**
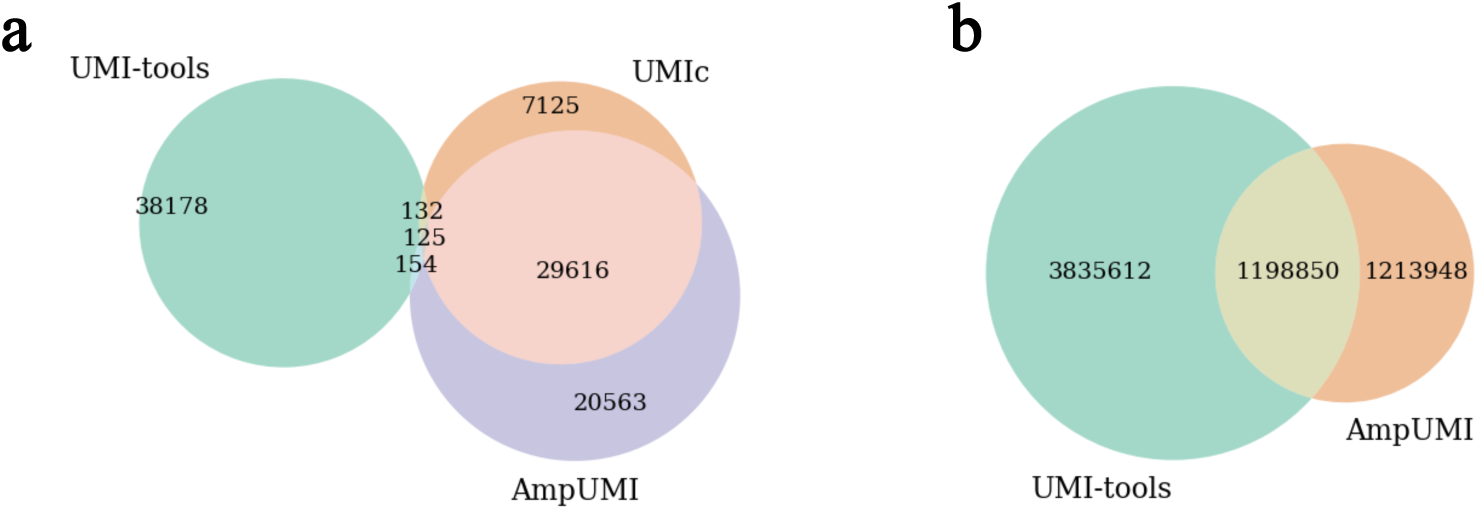
The performance comparison regarding the overlap and difference between unique read sets after PCR-deduplication by UMI-based methods. **a**. Venn diagram illustrates the result between UMI-tools[25], AmpUMI[31] and UMIc[35] on the dataset of SRR1543964. **b**. Venn diagram illustrates the result between UMI-tools[25] and AmpUMI[31] on the dataset of SRR28313972.

The Venn diagrams illustrate a significant overlap between UMIc- and AmpUMI-deduplicated reads on the datasets SRR1543964-SRR1543971. For instance, Figure 2a presents the Venn diagram when these three methods were applied to SRR1543964, where the areas of overlap between the circles represent the number of common sequences of the read sets. On the dataset SRR1543964, 80.05% of the unique reads identified by UMIc were also confirmed by AmpUMI. However, only 58.69% of the unique reads identified by AmpUMI were found by UMIc (Figure 2a). Across the eight datasets, UMIc and AmpUMI exhibit a mutual confirmation of 86.4% and 57.25% unique reads on average, as depicted in Figure 2a and in Supplementary Figure 1. It is worth noting that UMIc introduced many new sequences after PCR-deduplication, indicating a potential overcorrection of PCR biases and errors.

The unique reads identified by UMI-tools and AmpUMI show significant overlap across datasets such as SRR28313972, SRR28313990, and SRR28314008 (see Figure 2b and Supplementary Figure 1). For example, in the dataset SRR28313972, 23.8% of the unique reads identified by UMI-tools were also confirmed by AmpUMI, while AmpUMI found 49.7% of the unique reads identified by UMI-tools (Figure 2b).

The intersections between UMI-tools and UMIc and those between UMI-tools and AmpUMI were extremely small on the datasets SRR1543964-SRR1543971, and the intersections between UMI-tools and AmpUMI varied considerably across different groups of datasets. These findings stem from UMI-tools introducing numerous new sequences via alignment to a reference during its PCR-deduplication process. Its performance is likely influenced by factors such as the sequencing samples and the reference genome.

We did not perform overlap and difference analysis on the paired-end dataset SRR11207257. The reason is that the deduplicated reads consist entirely of new sequences and exhibit significant differences in counts, making comparative analysis meaningless from the perspective proposed in this article.

### UMI-based approaches in comparison to solely-computational PCR-deduplication methods

We benchmark the 9 solely-computational methods against the UMI-based ones on the 11 single-end datasets. We calculated the numbers of unique reads in the intersections between the sets of unique sequences produced by UMI-tools[25], AmpUMI[31] and UMIc[35] and those by the nine solely-computational methods NGSReadsTreatment[47], Nubeam-dedup[48], BioSeqZip[49], fastp[51], FastUniq[40], pRESTO[43], CD-HIT-DUP[38], ParDRe[44] and Minirmd[18].

The line charts presented in Figure 3a and Figure 4a compares the number of overlapped reads between the computational methods and UMI-based methods on the datasets SRR1543964 and SRR28313990, respectively. Supplementary Figures 2-10 present the performance of the other datasets. The results illustrate that the unique reads obtained using AmpUMI are a subset of those obtained by the computational methods NGSReadsTreatment, Nubeam-dedup, BioSeqZip, fastp, FastUniq, and pRESTO, as well as CD-HIT-DUP, ParDRe and Minirmd (without allowing any mismatches). All the sequences, except for those newly generated sequences after deduplication, are also present in the deduplicated set by the above computational methods. These observations are as expected because these solely computational approaches do not introduce new sequences while not correcting the PCR biases and errors. However, when mismatches (0-3) were allowed, the common sequences comparing UMI-based tools with the methods CD-HIT-DUP, ParDRe, or Minirmd decreased, which suggests that CD-HIT-DUP, ParDRe and Minirmd turned some actual biological reads into wrong states due to over error correction in the PCR-deduplication process. It is worth noting that the computational methods had a smaller amount of overlapping reads with UMI-tools on the datasets SRR1543964-SRR1543971 since UMI-tools introduced many new sequences in the step of alignment to the reference genome.

**Figure 3:**
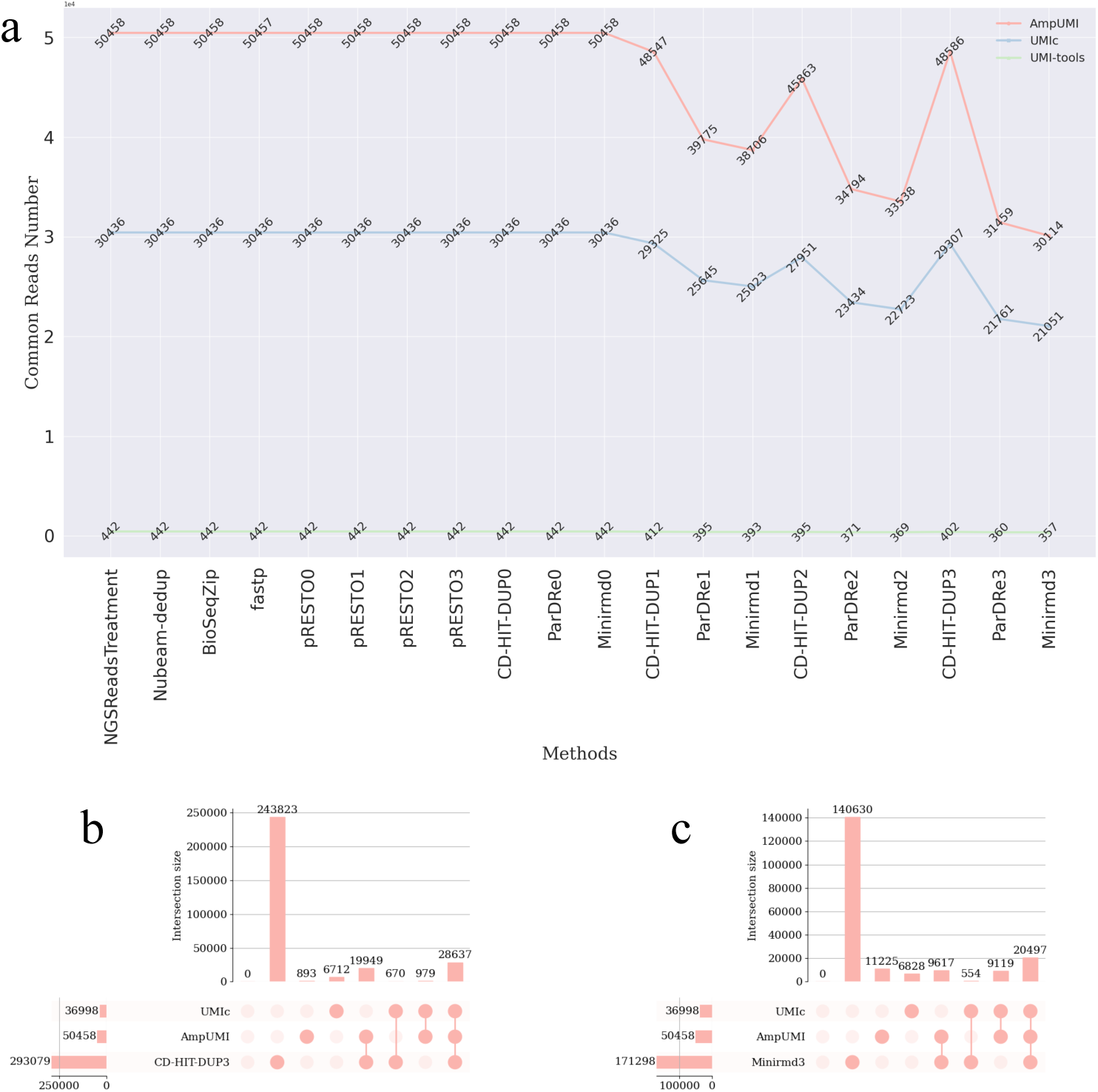
**(a)** Line chart comparing overlapped read numbers between each of the computational methods of NGSReadsTreatment[47], Nubeam-dedup[48], BioSeqZip[49], fastp[51], FastUniq[40], pRESTO[43], CD-HIT-DUP[38], ParDRe[44] and Minirmd[18] with each of the UMI-based methods of UMI-tools[25], AmpUMI[31] and UMIc[35] on the data set SRR1543964. **(b-c)** Upset plots for comparing the overlap and differences of deduplicated unique reads between the computational methods of CD-HIT-DUP[38] and Minirmd[18] and UMI-based methods of AmpUMI[31] and UMIc[35] on the data set SRR1543964. The number immediately following a method indicates the mismatch values allowed by that method.

**Figure 4:**
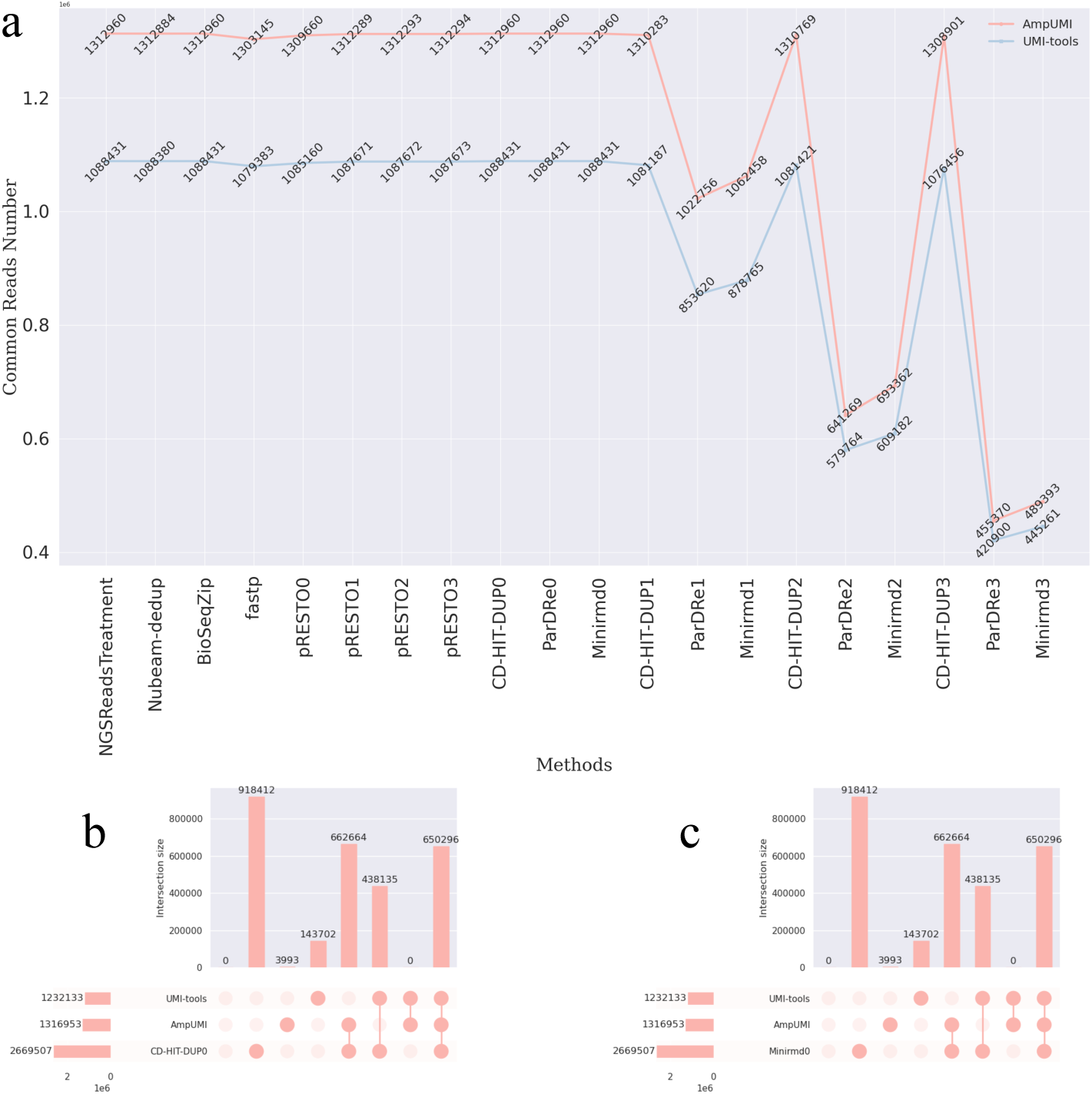
**(a)** Line chart comparing overlapped read numbers between each of the computational methods of NGSReadsTreatment[47], Nubeam-dedup[48], BioSeqZip[49], fastp[51], FastUniq[40], pRESTO[43], CD-HIT-DUP[38], ParDRe[44] and Minirmd[18] with each of the UMI-based methods of UMI-tools[25] and AmpUMI[31] on the data set SRR28313990. **(b-c)** Upset plots for comparing the overlap and differences of deduplicated unique reads between the computational methods of CD-HIT-DUP[38] and Minirmd[18] and UMI-based methods of AmpUMI[31] and UMI-tools[25] on the data set SRR28313990. The number immediately following a method indicates the mismatch values allowed by that method.

We have utilised UpSet plots to visualize the intersections between the deduplicated set of the unique reads obtained from every computational method and that from the UMI-based method AmpUMI and UMIc or UMI-tools. The rows in these UpSet plots stand for the read sets produced by PCR-deduplication methods and the columns stand for the intersections between these unique read sets. The highlighted cell or multiple cells connected with a line indicate the reading direction of an intersection. The size of the intersections is shown aligned with the rows, also as bar charts. The UpSet plots in Figure 3 b, c and Figure 4 b, c depict the deduplicated intersections between CD-HIT-DUP or minirmd and the methods AmpUMI and UMIc or UMI-tools on datasets SRR1543964 and SRR28313990, respectively. In the UpSet plots, where the column index begins at 0, superior performance in accurately identifying genuine reads is indicated by lower values in columns 1, 2, and 6 and higher values in columns 4, 5 and 7. Additionally, the proximity of the third column to the number of new sequences introduced by UMIc or UMI-tools implies better performance.

To facilitate the comparison of the performance of different methods, we use the heat map to compare the size of the seven intersections (columns 1-7 of the standard UpSet plot) of different methods, and the results are shown in Figure 5, and Supplementary Figures 11-20 depict the results on the data sets SRR1543965-SRR1543971, SRR28313972, SRR28313990, and SRR28314008. The results (Figure 5) in the first eleven rows of the heatmap indicate that these computational methods kept all the unique reads from the original datasets without correcting the PCR biases errors, except the methods fastp lost one unique read. CD-HIT-DUP, ParDRe and Minirmd can eliminate some PCR biases and errors by allowing mismatches when PCR-deduplication was processed (see the last nine rows of the heatmap in Figure 5). However, there are hundreds of thousands of distinct reads unique to these methods after PCR-deduplication (first column of the heatmap), suggesting that these methods left hundreds of thousands of erroneous reads uncorrected. The numbers in the fourth, fifth and seventh columns decreased, and the numbers in the second, third and sixth columns increased, demonstrating that these methods wrongly identified some biological reads as PCR errors.

**Figure 5:**
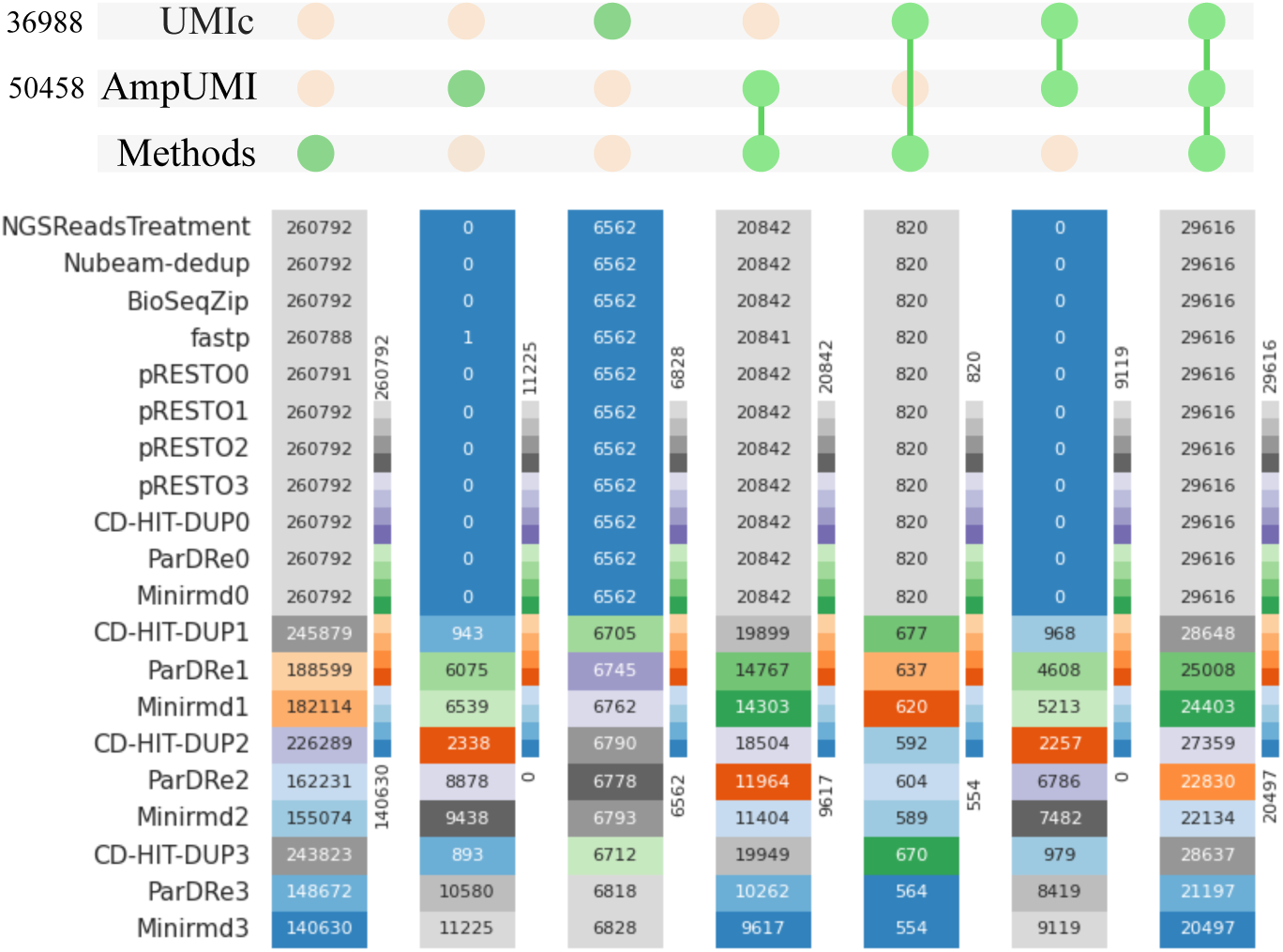
Heatmap for comparing overlapped reads number among each of the computational methods of NGSReadsTreatment[47], Nubeam-dedup[48], BioSeqZip[49], fastp[51], FastUniq[40], pRESTO[43], CD-HIT-DUP[38], ParDRe[44] and Minirmd[18] with the UMI-based methods of AmpUMI[31] and UMIc[35] on the data set SRR1543964. The number immediately following a method indicates the mismatch values allowed by that method.

### Runtime and memory consumption

We compared the CPU time and the peak memory usage consumed by UMI-tools[25], AmpUMI[31], UMIc[35], NGSReadsTreatment[47], Nubeam-dedup[48], BioSeqZip[49], fastp[51], pRESTO[43], CD-HIT-DUP[38], ParDRe[44] and Minirmd[18] on the datasets SRR1543964-SRR1543971 (see Table 5). AmpUMI achieved the fastest speed among the UMI-based methods and it was able to complete the deduplication in one minute with a memory usage of less than 550 megabytes (MB). The computational methods except for pRESTO, NGSReadsTreatment or ParDRe ran fast to complete in one minute with a peak memory ranging from 17 MB to 13329 MB. ParDRe was the slowest method that took up to more than 1 hour to complete on the dataset SRR1543969.

**Table 5:**
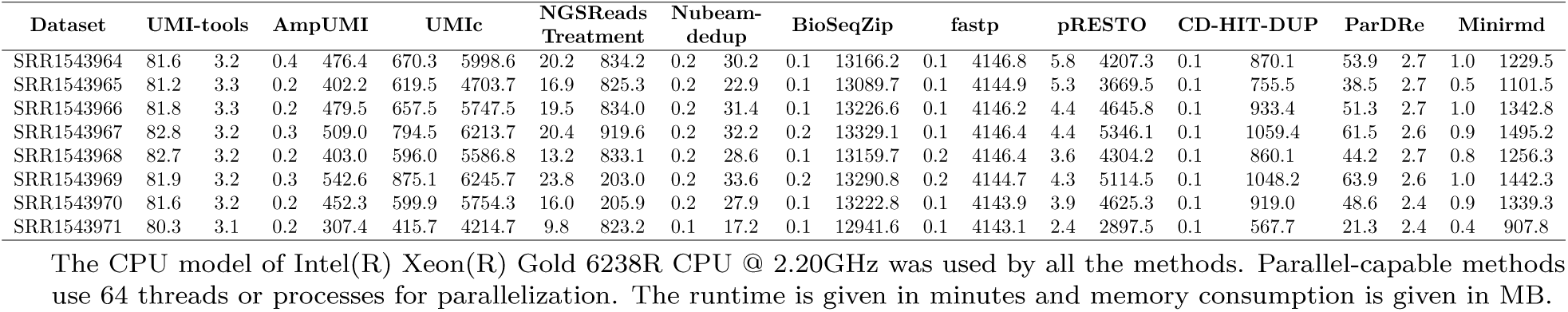
Runtime(T) and peak Memory(M) consumption by PCR-deduplication methods of UMI-tools[25], AmpUMI[31], UMIc[35], NGSReadsTreatment[47], Nubeam-dedup[48], BioSeqZip[49], fastp[51], pRESTO[43], CD-HIT-DUP[38], ParDRe[44] and Minirmd[18] on the datasets SRR1543964-SRR1543971.

In summary, the comparative analyses have revealed that: (1) AmpUMI is the best tool among different UMI-based PCR-deduplication methods as it runs fast and introduces the smallest number or almost does not introduce any new sequences after PCR-deduplication; (2) the computational PCR-deduplication methods, such as NGSReadsTreatment, Nubeam-dedup, BioSeqZip, fastp, and pRESTO, always produce the same deduplicated results. However, these methods do not consider any PCR biases and errors, and the results are the unique reads of the original data set. (3) The mismatch-allowed computational PCR-deduplication can eliminate some PCR biases and errors but could mistakenly identify genuine reads as near-duplicates to be removed and still leave hundreds of thousands of erroneous reads uncorrected. The best mismatch-allowed computational PCR-deduplication approach is CD-HIT-DUP as it keeps more original reads after PCR-deduplication.

### Performance comparisons for different error-correction methods

To compare the performance of the error-correction methods, we investigated the number of changes of unique reads, the number of corrected reads with the corresponding percentage out of the total reads, and the number of wrongly introduced new sequences after the correction. Table 6 presents the performance comparisons for the methods BFC[60], Bcool[72], Care[69, 70], Coral[65], Fiona[67], Lighter[53], Pollux[61] and RACER[58] on SRR1543964 and SRR28313972. The results on the other datasets are shown in Supplementary Tables 1-2. We observed a decrease in the number of unique reads for all of the methods after the error correction except for the methods RACER and BFC on the datasets SRR1543971 and SRR28313990. There is no relevant pattern in the percentage changes of unique reads in different data sets between different methods except for method BFC as it has the smallest change (e.g., only 0.41% on SRR1543964 and 0.05% on SRR28313972) in unique reads in each data set by comparing with other methods. For the other methods, Lighter made the largest reduction (a percentage decrease of 37.19%) on dataset SRR1543964, in contrast, Fiona made the largest reduction on dataset SRR28313972 (Table 6). On the other hand, some methods corrected a high percentage of reads. For example, RACER corrected a high percentage of reads (88.22%) on SRR1543964 while Fiona corrected 60.55% of reads on SRR28313972, suggesting that a significant proportion of 88.22% and 60.55% of the reads may have been wrongly identified as erroneous. Moreover, all of these methods generated tens to hundreds of thousands of non-existing reads after error correction. This result implies that these methods introduced numerous new errors during the correction process. Error correction results on multiple data sets show that no method is the best, and from the perspective of generating new sequences, Coral demonstrated the best performance on the datasets SRR1543964-SRR1543971, while Bcool demonstrated the best performance on the group of datasets SRR28313972, SRR28313990 and SRR28314008, as they introduced the smallest number of new reads.

**Table 6:**
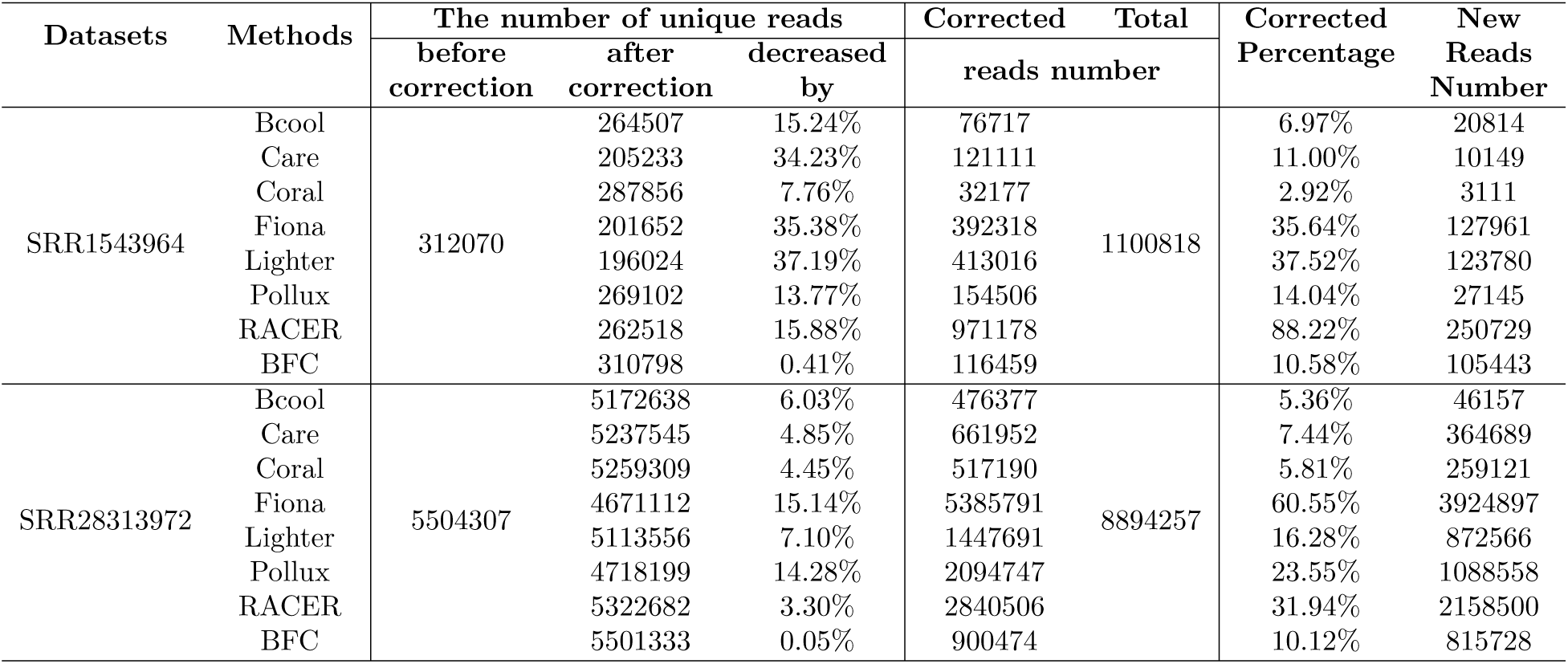
Summary of changes in unique reads, corrected reads, and erroneously introduced new reads after error correction using the error-correction methods of BFC[60], Bcool[72], Care[69, 70], Coral[65], Fiona[67], Lighter[53], Pollux[61] and RACER[58] on the datasets SRR1543964 and SRR28313972, respectively.

Furthermore, we analysed the overlaps and differences between the sets of unique reads obtained by different error-correction algorithms using UpSet plots. Comparative results can be found in Supplementary Figures 21-31 for the datasets SRR1543964-SRR1543971, SRR28313972, SRR28313990 and SRR28314008.

We observed that these algorithms produced many sequences unique to each algorithm, ranging from 1636 to 1681595 sequences. In contrast, all the algorithms’ intersection sizes on the eight data sets ranged from 1527 to 2811 on datasets SRR1543964-SRR1543971, and from 53451 to 602074 on the group of datasets SRR28313972, SRR28313990 and SRR28314008. The algorithm with the fewest unique sequences on SRR1543964-SRR1543971 was Coral and on the group of datasets SRR28313972, SRR28313990 and SRR28314008 was Bcool. This observation aligns with their respective percentages of corrected reads and the numbers of newly generated sequences, as indicated in Table 6 and Supplementary Tables 1-2. The smaller size of the common intersections between the algorithms suggests that the results obtained from these error-correction algorithms exhibit significant variation. In other words, employing various error-correction methods will result in divergent conclusions during subsequent downstream analyses.

### Runtime and memory consumption by different error-correction methods

We also compared the CPU time and the peak memory usage required by the methods BFC[60], Bcool[72], Care[69, 70], Coral[65], Fiona[67], Lighter[53], Pollux[61] and RACER[58] on the datasets SRR1543964-SRR1543971, as illustrated in Table 7. Lighter, BFC, Care, and RACER were able to correct errors in a minute with less memory, while Coral could complete in less than 10 minutes, but consumed much more memory up to 128.6 gigabytes (GB). Pollux ran slowest with time up to 26.4 hours. One of the reasons is that Pollux cannot run in parallel.

**Table 7:** Runtime(T) and peak Memory(M) consumption by error-correction methods of BFC[60], Bcool[72], Care[69, 70], Coral[65], Fiona[67], Lighter[53], Pollux[61] and RACER[58] on the datasets SRR1543964-SRR1543971.

| Dataset | BFC |  | Bcool |  | Care |  | Coral |  | Fiona |  | Lighter |  | Pollux |  | RACER |  |
| --- | --- | --- | --- | --- | --- | --- | --- | --- | --- | --- | --- | --- | --- | --- | --- | --- |
|  | T | M | T | M | T | M | T | M | T | M | T | M | T | M | T | M |
| SRR1543964 | 0.3 | 3.2 | 30.3 | 14.8 | 0.4 | 815.7 | 7.0 | 124136.4 | 84.1 | 2117.7 | 0.1 | 647.7 | 1263.2 | 1053.1 | 3.9 | 144.8 |
| SRR1543965 | 0.3 | 3.1 | 20.8 | 14.4 | 0.4 | 803.2 | 5.9 | 128940.7 | 67.4 | 1889.3 | 0.1 | 647.7 | 1106.8 | 1053.2 | 2.8 | 122.5 |
| SRR1543966 | 0.3 | 3.0 | 25.3 | 14.5 | 0.4 | 924.7 | 7.1 | 126143.5 | 96.1 | 2200.6 | 0.1 | 645.2 | 1329.0 | 1053.1 | 3.0 | 156.0 |
| SRR1543967 | 0.3 | 3.2 | 24.8 | 14.9 | 0.4 | 1030.8 | 9.1 | 108028.4 | 101.2 | 2155.0 | 0.1 | 649.4 | 1570.3 | 1053.2 | 4.1 | 178.2 |
| SRR1543968 | 0.3 | 3.2 | 17.8 | 14.4 | 0.4 | 840.3 | 6.4 | 123999.8 | 86.5 | 2119.7 | 0.1 | 649.5 | 1276.9 | 1039.6 | 2.9 | 149.3 |
| SRR1543969 | 0.3 | 3.1 | 35.4 | 13.1 | 0.5 | 991.9 | 8.9 | 124624.7 | 106.1 | 2264.0 | 0.1 | 647.7 | 1583.1 | 1119.8 | 4.0 | 172.3 |
| SRR1543970 | 0.3 | 3.1 | 19.6 | 13.6 | 0.4 | 895.7 | 7.7 | 122601.4 | 91.3 | 2191.1 | 0.1 | 649.6 | 1306.5 | 1053.1 | 3.1 | 156.7 |
| SRR1543971 | 0.2 | 3.2 | 14.3 | 12.9 | 0.3 | 606.3 | 5.1 | 128633.8 | 60.3 | 1710.5 | 0.1 | 649.5 | 782.8 | 585.1 | 2.2 | 100.1 |

The CPU model of Intel(R) Xeon(R) Gold 6238R CPU @ 2.20GHz was used by all the methods. Parallel-capable methods use 64 threads or processes for parallelization. The runtime is given in minutes and memory consumption is given in MB.

### How error-correction methods affect PCR-deduplication

We conducted duplicate removal using solely-computational PCR-deduplication methods on the error-corrected datasets from SRR1543964-SRR1543971, SRR28313972, SRR28313990 and SRR28314008 to investigate how error-correction methods affect PCR-deduplication. In detail, each of these datasets was firstly corrected by BFC[60], Bcool[72], Care[69, 70], Coral[65], Fiona[67], Lighter[53], Pollux[61] and RACER[58]. Then we removed duplicates using the three computational methods CD-HIT-DUP[38], ParDRe[44] and Minirmd[18] when the mismatch threshold was set differently.

The line charts in Figure 6, Figure 7 and Supplementary Figures 32-62 illustrate the intersection size of the sets of the deduplicated reads after error correction in comparison with the corresponding sets directly obtained through UMI-based PCR-deduplication methods. Three solid lines, representing CD-HIT-DUP, ParDRe, and Minirmd (with a permissible mismatch of not bigger than 3 bases), are contrasted with a dashed line depicting CD-HIT-DUP with no mismatch allowance.

**Figure 6:**
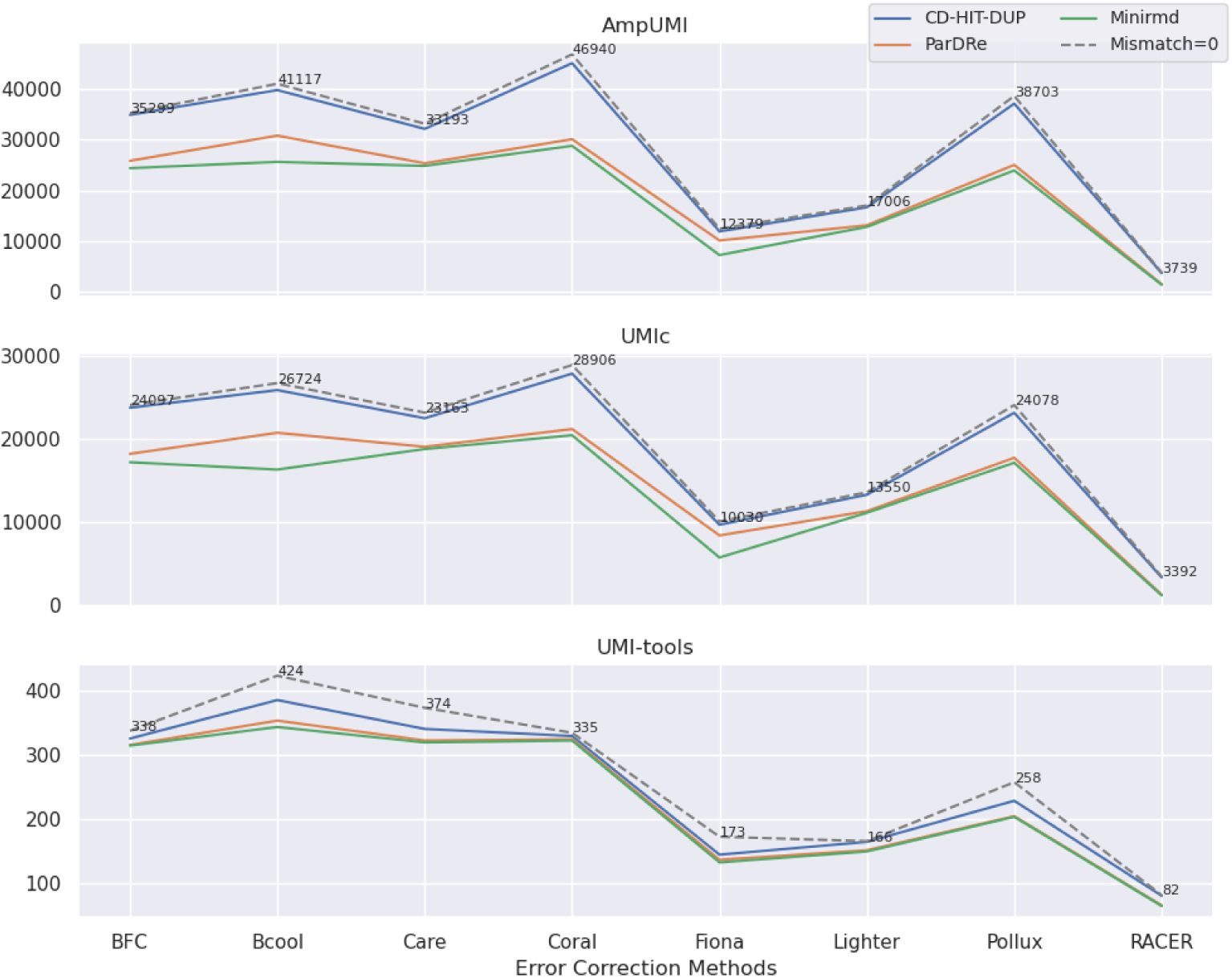
Line charts comparing the overlapped read numbers in the deduplicated read set by the PCR-deduplication methods of CD-HIT-DUP[38], ParDRe[44] and Minirmd[18] on error-corrected dataset SRR1543964, with each UMI-based PCR-deduplication methods of UMI-tools[25], AmpUMI[31] and UMIc[35] on dataset SRR1543964. Error correction was performed using error-correction methods of BFC[60], Bcool[72], Care[69, 70], Coral[65], Fiona[67], Lighter[53], Pollux[61] and RACER[58], respectively. CD-HIT-DUP, ParDRe, and Minirmd employed a mismatched number set to 3. The dashed line labeled ’Mismatch=0’ represents results obtained by CD-HIT-DUP with a mismatch setting of 0.

**Figure 7:**
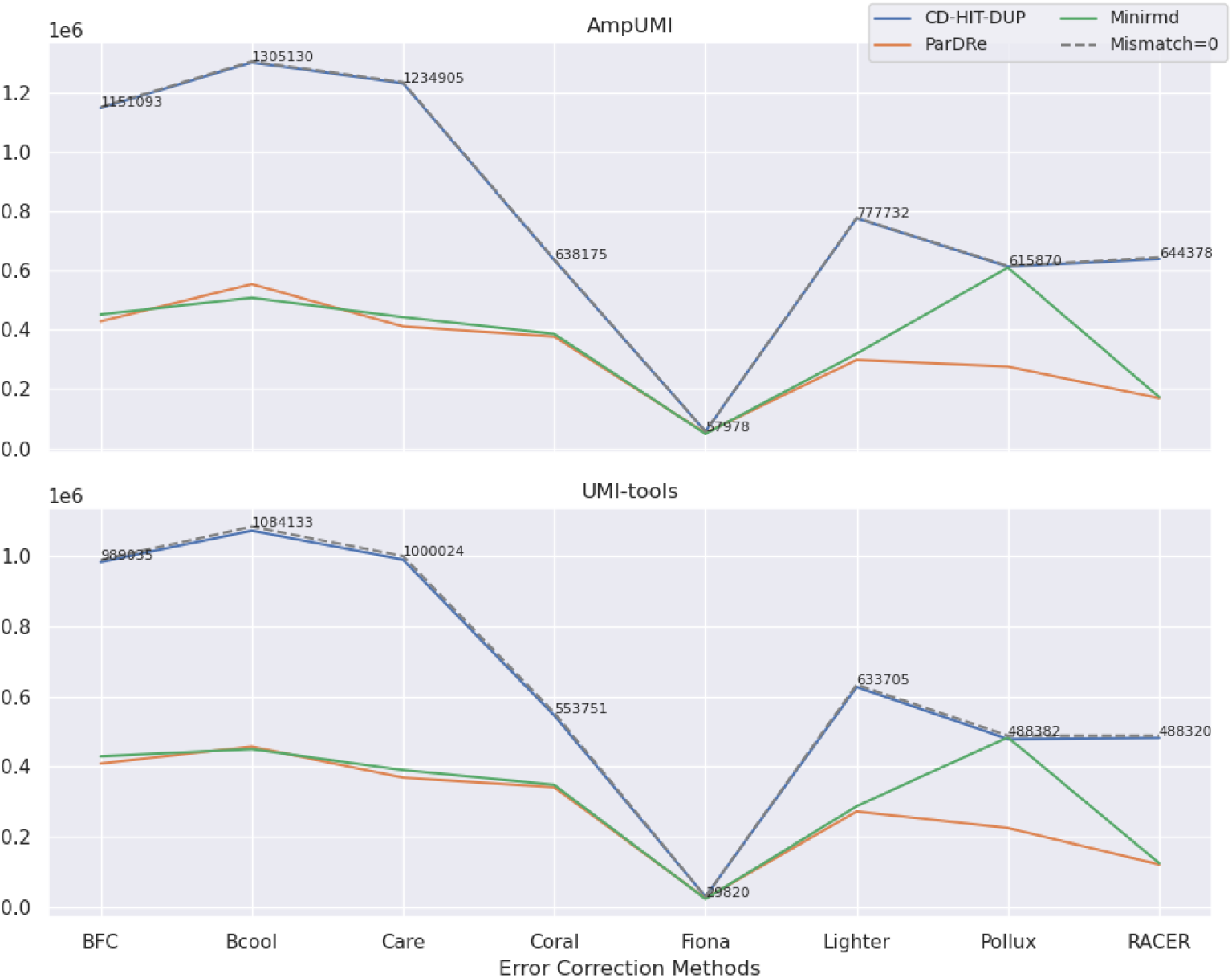
Line charts comparing the overlapped read numbers in the deduplicated read set by the PCR-deduplication methods of CD-HIT-DUP[38], ParDRe[44] and Minirmd[18] on error-corrected dataset SRR28313990, with each UMI-based PCR-deduplication methods of UMI-tools[25] and AmpUMI[31] on dataset SRR28313990. Error correction was performed using error-correction methods of BFC[60], Bcool[72], Care[69, 70], Coral[65], Fiona[67], Lighter[53], Pollux[61] and RACER[58], respectively. CD-HIT-DUP, ParDRe, and Minirmd employed a mismatched number set to 3. The dashed line labeled ’Mismatch=0’ represents results obtained by CD-HIT-DUP with a mismatch setting of 0.

Comparing the intersection size of unique reads after the PCR-deduplication process applied to error- corrected reads (Figure 6 and Figure 7 ) with those of the directly deduplicated reads (Figure 3a and Figure 4a), decrease in the number is observed on different group of datasets. This decrease suggests that the error corrections erroneously identified some authentic biological sequences as PCR errors. Similar outcomes observed across other datasets, as depicted in Supplementary Figures 32-62, indicate that the current error- correction methods are not advantageous for PCR-deduplication. In other words, the employment of the existing error-correction methods appears to be an unnecessary step before the PCR-deduplication process.

Moreover, most importantly, the size of both the directly deduplicated read set by computational methods and the deduplicated read set after error correction is significantly larger than that of the set of the deduplicated reads generated by UMI-based PCR-deduplication methods. For instance, in the dataset SRR1543964, the directly deduplicated read set by CD-HIT-DUP (with 2 mismatches) consists of 272,744 unique reads (Table 3), while the Coral error-corrected read set comprises 287,856 unique reads (Table 6). However, the AmpUMI- deduplicated read set only contains 50,458 unique reads (Table 3). Similar trends are observed in the other datasets for these PCR-deduplication and error-correction methods. This analysis demonstrates that both existing PCR-deduplication and error-correction methods leave a considerable number of erroneous reads untouched while generating a considerable number of new sequences by error-correction approaches.

Furthermore, while error-correction techniques offer only marginal benefits to PCR-deduplication, the effectiveness varies among different methods. In the datasets SRR1543964-SRR1543971, the Coral method yielded the highest number of overlapped reads with both the AmpUMI and UMIc methods (refer to Figure 6 and Supplementary Figures 32-54); in contrast, RACER resulted in the smallest size of the corresponding intersections. On the datasets SRR28313972, SRR28313990 and SRR28314008, Bcool yielded the highest number of overlapped reads with both of the AmpUMI and UMIc methods (refer to Figure 7 and Supplementary Figures 55-62) while Fiona resulted in the smallest size of the corresponding intersections.

In summary, these comparative studies have revealed that: (1) All error-correction methods introduce many non-existent sequences after error correction and leave many erroneous reads untouched. (2) No error-correction method works best across all the evaluation datasets. Coral and Bcool introduce the fewest new sequences and exhibit the largest overlap of unique reads with the UMI-based deduplicated read set on the two different group of datasets.

## Discussion

This study conducted a comprehensive investigation on both PCR-deduplication and error-correction, exploring the impact of error correction on PCR-deduplication as well as comparing the performance of these state-of-the-art methods from novel perspectives based on UMI datasets of short reads.

While incorporating UMIs proves to be an effective strategy for mitigating data uncertainty brought up by PCR biases and errors, it is crucial to acknowledge significant practical challenges associated with their use. Firstly, PCR biases and errors may still occur within UMIs, posing a challenge for existing methods to differentiate between PCR duplicates and genuine biological sequences, particularly when PCR errors arise during the early-cycle PCR process. Similarly, the identification and resolution of errors in the sequence during the initial PCR cycles present a complex task for existing error-correction methods. One previous experimental analysis presented in the literature [33] has demonstrated that the prevalence of PCR duplicates is primarily determined by the quantity of initial materials and sequencing depth and there is no additional impact attributed to the cycles of PCR amplification. Secondly, UMI collisions may occur during library preparation, especially in large-scale sequencing when short-length UMIs are used. Most existing PCR-deduplication methods are not equipped to handle UMI collisions. Thirdly, UMI-based strategies are characterized by their time-consuming and expensive nature, rendering them less practical for large-scale applications.

In the PCR-deduplication and error-correction processes, it remains unclear whether there exists a means to discern whether errors in a sequence stem from PCR or fluorophore crosstalk. Future studies could delve into these issues more comprehensively, as their resolution holds significant implications for sequencing library preparation and sequencing technology. Alignment-based PCR-deduplication methods may have the possibility of deleting actual biological sequences because they are based on a reference genome or transcriptome to remove PCR errors. However, these references were built on implicit assumptions. For example, substantial evidence supports that error, noise, and bias exist in de novo transcriptome assemblies [10].

After conducting a deep analysis, we have uncovered notable limitations within existing algorithms working without UMI sequence data. The reads deduplicated solely through computational PCR-deduplication and error-correction methods display significant disparities when compared to those generated by UMI-based techniques. This observation aligns with the established conclusion in the literature [33] that PCR duplicate removal without UMI sequence data tends to be overly aggressive.

Moreover, even though NGS platforms (e.g., Illumina instruments) share some standard techniques, their specific sequencing methods differ from each other, leading to reads with different error profiles [23, 75]. In addition, these error profiles may also vary depending on the sequencing task. For instance, the sequencing error profiles of miRNA sequencing data may differ from whole-genome sequencing data or synthetic sequencing data. Therefore, further and more investigation and analysis are necessary to uncover and understand the underlying causes of PCR bias and sequencing errors.

Addressing these challenges can enhance the accuracy and reliability of downstream analyses and enable a more robust interpretation of NGS data. An ideal sequencing error removal algorithm should accurately restore erroneous reads to their correct states without introducing new errors or sequences. Here, we propose potential solutions for developing novel methodologies and for improving existing computational methods to mitigate PCR biases and sequencing errors in short reads.

Firstly, current PCR-deduplication and error-correction methods can be improved by incorporating iterative and screening mechanisms to prevent the generation of non-existent sequences. Existing mismatch- allowed PCR-deduplication methods remove near-duplicates to eliminate PCR biases and errors. However, experimental evidence shows that these methods mistakenly identify some genuine identical reads as near-duplicates to be removed. Enhancing their performance may be achieved by introducing additional information and parameters, such as sequence frequency, to filter out true near-duplicates accurately.

Secondly, PCR-deduplication tools and error-correction methods can be mutually applied and extended. The PCR-deduplication method can record sequencing IDs or store the accumulated counts for each unique read while removing duplicated reads, resulting in corrected reads with preserved read abundance. On the other hand, error-correction methods can be upgraded by generating unique reads as part of the output for PCR-deduplication purposes after correction.

Thirdly, many current computational methods for PCR-deduplication and error-correction strategies rely on *k*-mer-based, Multiple Sequence Alignment (MSA)-based, de Bruijn graph (DBG), hashing or minimiser techniques to establish relationships among generated sequences. However, these approaches often fail to address the fundamental causes of PCR and sequencing biases and errors. To overcome this limitation, the introduction of new mechanisms is crucial. Understanding the intricacies of PCR and fluorophore crosstalk error mechanisms, coupled with the incorporation of machine learning and deep learning techniques, may provide more effective solutions. An illustration of this need is evident in a study where the predominant factor influencing PCR bias was identified as the relationship between library complexity and sequencing depth. Amplification noise, along with its correlation with the number of PCR cycles, was found to play a secondary role in contributing to this artifact. Recognizing and incorporating these insights can be instrumental in identifying inherent error patterns specific to each sequencing platform used for sequencing particular biomolecular samples.

## Conclusion

Next-Generation Sequencing (NGS) data has become increasingly prevalent in various bioinformatics applications in recent years. However, addressing the PCR and base calling errors is a long-standing problem and is crucial for any downstream analysis whenever NGS short read is involved. PCR-deduplication and error-correction tools utilise computational techniques to address biases and errors introduced during short-read sequencing. We have provided a comprehensive review of the existing PCR-deduplication and error-correction methods by assessing the performance of these algorithms from novel perspectives.

This study identified the limitations and deficiencies of existing PCR-deduplication and error-correction methods and proposed potential solutions from a broader perspective. The comparative analyses have revealed that the existing computational-based PCR-deduplication and error-correction methods can eliminate some PCR and sequencing errors but still leave hundreds of thousands of erroneous reads uncorrected. Most solely-computational PCR-deduplication methods do not address bias and error removal, and mismatch-allowed approaches can remove some PCR errors but also delete some genuine identical reads as near-duplicates or convert erroneous reads into different error states instead of their correct or normal states. All error-correction methods introduce tens of thousands of new sequences after error restoration and the unique read sets obtained from different error-correction methods show limited overlap, as these methods treat and correct just varying percentages of erroneous reads. Employing the existing error-correction methods appears to be unnecessary as a preprocessing step for the PCR-deduplication process.

## Supplementary Data

Supplementary materials, including Supplementary Figures 1-62 and Supplementary Tables 1-2, are extended data supporting the analysis in this study.

## Data Availability

The 11 UMI-based single-end sequencing datasets of the accession IDs [SRR1543964](https://www.ncbi.nlm.nih.gov/Traces/study/?acc=SRP045430&o=acc_s%3Aa) - [SRR1543971](https://www.ncbi.nlm.nih.gov/Traces/study/?acc=SRP045430&o=acc_s%3Aa), SRR28313972, SRR28313990, and SRR28314008 together with a pair-end sequencing dataset of the accession ID SRR11207257 were downloaded from SRA archive (https://www.ncbi.nlm.nih.gov/sra). The Python scripts for comparative analysis in this study are open source and publicly available at the Github repository https://github.com/Jappy0/Deduplication_ErrorCorrection.

## Author contributions statement

P.P. and J.L. initiated this survey paper. P.P. conducted the comparison experiments and wrote the manuscript. S.S., T.L., W.L. and J.L. revised the manuscript.

## Acknowledgments

The authors thank the anonymous reviewers for their valuable suggestions and thank for the computational resources provided by the UTS (University of Technology Sydney) eResearch High-Performance Computer Facilities.

## References

[1] Jay Shendure, Shankar Balasubramanian, George M Church, Walter Gilbert, Jane Rogers, Jeffery A Schloss, and Robert H Waterston. Dna sequencing at 40: past, present and future. Nature, 550(7676):345– 353, 2017.

[2] Kimberly R Kukurba and Stephen B Montgomery. Rna sequencing and analysis. Cold Spring Harbor Protocols, 2015(11):951–969, 2015.

[3] Justin D. Silverman, Rachael J. Bloom, Sharon Jiang, Heather K. Durand, Eric Dallow, Sayan Mukherjee, and Lawrence A. David. Measuring and mitigating PCR bias in microbiota datasets. 17(7):e1009113.

[4] Mark T. W. Ebbert, Mark E. Wadsworth, Lyndsay A. Staley, Kaitlyn L. Hoyt, Brandon Pickett, Justin Miller, John Duce, John S. K. Kauwe, Perry G. Ridge, and for the Alzheimer’s Disease Neuroimaging Initiative. Evaluating the necessity of PCR duplicate removal from next-generation sequencing data and a comparison of approaches. 17(7):239.

[5] Vivien Marx. How to deduplicate pcr. Nature Methods, 14(5):473–476, 2017.

[6] Kevin M. Gao, Alan G. Derr, Zhiru Guo, Kerstin Nündel, Ann Marshak-Rothstein, Robert W. Finberg, and Jennifer P. Wang. Human nasal wash RNA-Seq reveals distinct cell-specific innate immune responses in influenza versus SARS-CoV-2. 6(22):e152288.

[7] Swati Parekh, Christoph Ziegenhain, Beate Vieth, Wolfgang Enard, and Ines Hellmann. The impact of amplification on differential expression analyses by RNA-seq. 6(1):25533.

[8] Wenming Xiao, Luyao Ren, Zhong Chen, Li Tai Fang, Yongmei Zhao, Justin Lack, Meijian Guan, Bin Zhu, Erich Jaeger, Liz Kerrigan, Thomas M. Blomquist, Tiffany Hung, Marc Sultan, Kenneth Idler, Charles Lu, Andreas Scherer, Rebecca Kusko, Malcolm Moos, Chunlin Xiao, Stephen T. Sherry, Ogan D. Abaan, Wanqiu Chen, Xin Chen, Jessica Nordlund, Ulrika Liljedahl, Roberta Maestro, Maurizio Polano, Jiri Drabek, Petr Vojta, Sulev Kõks, Ene Reimann, Bindu Swapna Madala, Timothy Mercer, Chris Miller, Howard Jacob, Tiffany Truong, Ali Moshrefi, Aparna Natarajan, Ana Granat, Gary P. Schroth, Rasika Kalamegham, Eric Peters, Virginie Petitjean, Ashley Walton, Tsai-Wei Shen, Keyur Talsania, Cristobal Juan Vera, Kurt Langenbach, Maryellen de Mars, Jennifer A. Hipp, James C. Willey, Jing Wang, Jyoti Shetty, Yuliya Kriga, Arati Raziuddin, Bao Tran, Yuanting Zheng, Ying Yu, Margaret Cam, Parthav Jailwala, Cu Nguyen, Daoud Meerzaman, Qingrong Chen, Chunhua Yan, Ben Ernest, Urvashi Mehra, Roderick V. Jensen, Wendell Jones, Jian-Liang Li, Brian N. Papas, Mehdi Pirooznia, Yun-Ching Chen, Fayaz Seifuddin, Zhipan Li, Xuelu Liu, Wolfgang Resch, Jingya Wang, Leihong Wu, Gokhan Yavas, Corey Miles, Baitang Ning, Weida Tong, Christopher E. Mason, Eric Donaldson, Samir Lababidi, Louis M. Staudt, Zivana Tezak, Huixiao Hong, Charles Wang, and Leming Shi. Toward best practice in cancer mutation detection with whole-genome and whole-exome sequencing. Nature Biotechnology, 39(9):1141–1150, Sep 2021.

[9] Mahdi Heydari, Giles Miclotte, Piet Demeester, Yves Van de Peer, and Jan Fostier. Evaluation of the impact of Illumina error correction tools on de novo genome assembly. 18(1):374.

[10] Adam H. Freedman, Michele Clamp, and Timothy B. Sackton. Error, noise and bias in de novo transcriptome assemblies. Molecular Ecology Resources, 21(1):18–29, 2021.

[11] Francesco Marchetti, Renato Cardoso, Connie L Chen, George R Douglas, Joanne Elloway, Patricia A Escobar, Tod Harper Jr, Robert H Heflich, Darren Kidd, Anthony M Lynch, et al. Error-corrected next-generation sequencing to advance nonclinical genotoxicity and carcinogenicity testing. Nature Reviews Drug Discovery, 22(3):165–166, 2023.

[12] Manuela Piazzi, Alberto Bavelloni, Sara Salucci, Irene Faenza, and William L. Blalock. Alternative splicing, rna editing, and the current limits of next generation sequencing. Genes, 14(7), 2023.

[13] Aristeidis G. Telonis, Rogan Magee, Phillipe Loher, Inna Chervoneva, Eric Londin, and Isidore Rigoutsos. Knowledge about the presence or absence of miRNA isoforms (isomiRs) can successfully discriminate amongst 32 TCGA cancer types. Nucleic Acids Research, 45(6):2973–2985, 02 2017.

[14] Yong Huang, Meiqi Shang, Tingting Liu, and Kejian Wang. High-throughput methods for genome editing: the more the better. Plant Physiology, 188(4):1731–1745, 2022.

[15] James A. Casbon, Robert J. Osborne, Sydney Brenner, and Conrad P. Lichtenstein. A method for counting PCR template molecules with application to next-generation sequencing. Nucleic Acids Research, 39(12):e81–e81, 04 2011.

[16] Nicolas C. Rochette, Angel G. Rivera-Colón, Jessica Walsh, Thomas J. Sanger, Shane C. Campbell-Staton, and Julian M. Catchen. On the causes, consequences, and avoidance of pcr duplicates: Towards a theory of library complexity. Molecular Ecology Resources, 23(6):1299–1318, 2023.

[17] Justus M. Kebschull and Anthony M. Zador. Sources of PCR-induced distortions in high-throughput sequencing data sets. Nucleic Acids Research, 43(21):e143–e143, 07 2015.

[18] Yuansheng Liu, Xiaocai Zhang, Quan Zou, and Xiangxiang Zeng. Minirmd: accurate and fast duplicate removal tool for short reads via multiple minimizers. Bioinformatics, 37(11):1604–1606, 2021.

[19] Matthew S Burriesci, Erik M Lehnert, and John R Pringle. Fulcrum: condensing redundant reads from high-throughput sequencing studies. Bioinformatics, 28(10):1324–1327, 2012.

[20] Mark A DePristo, Eric Banks, Ryan Poplin, Kiran V Garimella, Jared R Maguire, Christopher Hartl, Anthony A Philippakis, Guillermo Del Angel, Manuel A Rivas, Matt Hanna, et al. A framework for variation discovery and genotyping using next-generation dna sequencing data. Nature genetics, 43(5):491–498, 2011.

[21] Robert Schmieder and Robert Edwards. Quality control and preprocessing of metagenomic datasets. Bioinformatics, 27(6):863–864, 2011.

[22] Xiaotu Ma, Ying Shao, Liqing Tian, Diane A Flasch, Heather L Mulder, Michael N Edmonson, Yu Liu, Xiang Chen, Scott Newman, Joy Nakitandwe, et al. Analysis of error profiles in deep next-generation sequencing data. Genome biology, 20:1–15, 2019.

[23] Nicholas Stoler and Anton Nekrutenko. Sequencing error profiles of illumina sequencing instruments. NAR genomics and bioinformatics, 3(1):lqab019, 2021.

[24] Teemu Kivioja, Anna Vähärautio, Kasper Karlsson, Martin Bonke, Martin Enge, Sten Linnarsson, and Jussi Taipale. Counting absolute numbers of molecules using unique molecular identifiers. Nature methods, 9(1):72–74, 2012.

[25] Tom Smith, Andreas Heger, and Ian Sudbery. UMI-tools: Modeling sequencing errors in Unique Molecular Identifiers to improve quantification accuracy. Genome Research, 27(3):491–499, 2017.

[26] Xiyu Peng and Karin S Dorman. Accurate estimation of molecular counts from amplicon sequence data with unique molecular identifiers. Bioinformatics, 39(1):btad002, 01 2023.

[27] Baraa Orabi, Emre Erhan, Brian McConeghy, Stanislav V Volik, Stephane Le Bihan, Robert Bell, Colin C Collins, Cedric Chauve, and Faraz Hach. Alignment-free clustering of umi tagged dna molecules. Bioinformatics, 35(11):1829–1836, 2019.

[28] Keith Mitchell, Jaqueline J. Brito, Igor Mandric, Qiaozhen Wu, Sergey Knyazev, Sei Chang, Lana S. Martin, Aaron Karlsberg, Ekaterina Gerasimov, Russell Littman, Brian L. Hill, Nicholas C. Wu, Harry Ta-egyun Yang, Kevin Hsieh, Linus Chen, Eli Littman, Taylor Shabani, German Enik, Douglas Yao, Ren Sun, Jan Schroeder, Eleazar Eskin, Alex Zelikovsky, Pavel Skums, Mihai Pop, and Serghei Mangul. Benchmarking of computational error-correction methods for next-generation sequencing data. Genome Biology, 21(1):71, 2020.

[29] Charles Girardot, Jelle Scholtalbers, Sajoscha Sauer, Shu-Yi Su, and Eileen EM Furlong. Je, a versatile suite to handle multiplexed ngs libraries with unique molecular identifiers. BMC bioinformatics, 17:1–6, 2016.

[30] Serghei Mangul, Sarah Van Driesche, Lana S. Martin, Kelsey C. Martin, and Eleazar Eskin. UMI-Reducer: Collapsing duplicate sequencing reads via Unique Molecular Identifiers, 2017.

[31] Kendell Clement, Rick Farouni, Daniel E. Bauer, and Luca Pinello. AmpUMI: Design and analysis of unique molecular identifiers for deep amplicon sequencing. Bioinformatics (Oxford, England), 34(13):i202– i210, 2018.

[32] Swati Parekh, Christoph Ziegenhain, Beate Vieth, Wolfgang Enard, and Ines Hellmann. zumis-a fast and flexible pipeline to process rna sequencing data with umis. Gigascience, 7(6):giy059, 2018.

[33] Yu Fu, Pei-Hsuan Wu, Timothy Beane, Phillip D. Zamore, and Zhiping Weng. Elimination of PCR duplicates in RNA-seq and small RNA-seq using unique molecular identifiers. BMC Genomics, 19(1):531, 2018.

[34] Daniel Liu. Algorithms for efficiently collapsing reads with unique molecular identifiers. PeerJ, 7(e8275):e8275, December 2019.

[35] Maria Tsagiopoulou, Maria Christina Maniou, Nikolaos Pechlivanis, Anastasis Togkousidis, Michaela Kotrová, Tobias Hutzenlaub, Ilias Kappas, Anastasia Chatzidimitriou, and Fotis Psomopoulos. UMIc: A Preprocessing Method for UMI Deduplication and Reads Correction. Frontiers in Genetics, 12:660366, 2021.

[36] Shifu Chen, Yanqing Zhou, Yaru Chen, Tanxiao Huang, Wenting Liao, Yun Xu, Zhicheng Li, and Jia Gu. Gencore: An efficient tool to generate consensus reads for error suppressing and duplicate removing of NGS data. BMC Bioinformatics, 20(23):606, 2019.

[37] Heng Li, Bob Handsaker, Alec Wysoker, Tim Fennell, Jue Ruan, Nils Homer, Gabor Marth, Goncalo Abecasis, Richard Durbin, and 1000 Genome Project Data Processing Subgroup. The Sequence Alignment/Map format and SAMtools. Bioinformatics (Oxford, England), 25(16):2078–2079, 2009.

[38] Ying Huang, Beifang Niu, Ying Gao, Limin Fu, and Weizhong Li. CD-HIT Suite: A web server for clustering and comparing biological sequences. *Bioinformatics (Oxford*, England), 26(5):680–682, 2010.

[39] Ergude Bao, Tao Jiang, Isgouhi Kaloshian, and Thomas Girke. SEED: Efficient clustering of next-generation sequences. Bioinformatics, 27(18):2502–2509, 2011.

[40] Haibin Xu, Xiang Luo, Jun Qian, Xiaohui Pang, Jingyuan Song, Guangrui Qian, Jinhui Chen, and Shilin Chen. FastUniq: A fast de novo duplicates removal tool for paired short reads. PloS One, 7(12):e52249, 2012.

[41] Zechen Chong, Jue Ruan, and Chung-I. Wu. Rainbow: An integrated tool for efficient clustering and assembling RAD-seq reads. Bioinformatics, 28(21):2732–2737, 2012.

[42] Picard toolkit. https://broadinstitute.github.io/picard/, 2019.

[43] Jason A. Vander Heiden, Gur Yaari, Mohamed Uduman, Joel N.H. Stern, Kevin C. O’Connor, David A. Hafler, Francois Vigneault, and Steven H. Kleinstein. pRESTO: A toolkit for processing high-throughput sequencing raw reads of lymphocyte receptor repertoires. Bioinformatics, 30(13):1930–1932, 2014.

[44] Jorge González-Domínguez and Bertil Schmidt. ParDRe: Faster parallel duplicated reads removal tool for sequencing studies. Bioinformatics (Oxford, England), 32(10):1562–1564, 2016.

[45] Roberto R Expósito, Jorge Veiga, Jorge González-Domínguez, and Juan Touriño. MarDRe: Efficient MapReduce-based removal of duplicate DNA reads in the cloud. Bioinformatics, 33(17):2762–2764, 2017.

[46] Vikas Bansal. A computational method for estimating the PCR duplication rate in DNA and RNA-seq experiments. BMC Bioinformatics, 18(3):43, 2017.

[47] Antonio Sérgio Cruz Gaia, Pablo Henrique Caracciolo Gomes de Sá, Mônica Silva de Oliveira, and Adonney Allan de Oliveira Veras. NGSReadsTreatment – A Cuckoo Filter-based Tool for Removing Duplicate Reads in NGS Data. Scientific Reports, 9(1):11681, 2019.

[48] Hang Dai and Yongtao Guan. Nubeam-dedup: A fast and RAM-efficient tool to de-duplicate sequencing reads without mapping. Bioinformatics, 36(10):3254–3256, 2020.

[49] Gianvito Urgese, Emanuele Parisi, Orazio Scicolone, Santa Di Cataldo, and Elisa Ficarra. BioSeqZip: A collapser of NGS redundant reads for the optimization of sequence analysis. Bioinformatics, 36(9):2705– 2711, 2020.

[50] Xiyu Peng and Karin S Dorman. AmpliCI: a high-resolution model-based approach for denoising Illumina amplicon data. Bioinformatics, 36(21):5151–5158, 07 2020.

[51] Shifu Chen, Yanqing Zhou, Yaru Chen, and Jia Gu. Fastp: An ultra-fast all-in-one FASTQ preprocessor. Bioinformatics, 34(17):i884–i890, 2018.

[52] Petr Danecek, James K Bonfield, Jennifer Liddle, John Marshall, Valeriu Ohan, Martin O Pollard, Andrew Whitwham, Thomas Keane, Shane A McCarthy, Robert M Davies, and Heng Li. Twelve years of SAMtools and BCFtools. GigaScience, 10(giab008), 2021.

[53] Li Song, Liliana Florea, and Ben Langmead. Lighter: Fast and memory-efficient sequencing error correction without counting. Genome Biology, 15(11):509, 2014.

[54] Paul Greenfield, Konsta Duesing, Alexie Papanicolaou, and Denis C. Bauer. Blue: Correcting sequencing errors using consensus and context. Bioinformatics (Oxford, England), 30(19):2723–2732, 2014.

[55] Yun Heo, Xiao-Long Wu, Deming Chen, Jian Ma, and Wen-Mei Hwu. BLESS: Bloom filter-based error correction solution for high-throughput sequencing reads. Bioinformatics (Oxford, England), 30(10):1354–1362, 2014.

[56] Yun Heo, Anand Ramachandran, Wen-Mei Hwu, Jian Ma, and Deming Chen. BLESS 2: Accurate, memory-efficient and fast error correction method. Bioinformatics, 32(15):2369–2371, 2016.

[57] Yongchao Liu, Jan Schröder, and Bertil Schmidt. Musket: A multistage k-mer spectrum-based error corrector for Illumina sequence data. Bioinformatics, 29(3):308–315, 2013.

[58] Lucian Ilie and Michael Molnar. RACER: Rapid and accurate correction of errors in reads. Bioinformatics (Oxford, England), 29(19):2490–2493, 2013.

[59] Jared T. Simpson and Richard Durbin. Efficient de novo assembly of large genomes using compressed data structures. Genome Research, 22(3):549–556, 2012.

[60] Heng Li. BFC: Correcting Illumina sequencing errors. Bioinformatics (Oxford, England), 31(17):2885– 2887, 2015.

[61] Eric Marinier, Daniel G. Brown, and Brendan J. McConkey. Pollux: Platform independent error correction of single and mixed genomes. BMC Bioinformatics, 16(1):10, 2015.

[62] Maciej Dlugosz and Sebastian Deorowicz. RECKONER: Read error corrector based on KMC. Bioinformatics (Oxford, England), 33(7):1086–1089, 2017.

[63] Mustafa Abdallah, Ashraf Mahgoub, Hany Ahmed, and Somali Chaterji. Athena: Automated Tuning of k-mer based Genomic Error Correction Algorithms using Language Models. Scientific Reports, 9(1):16157, 2019.

[64] Atul Sharma, Pranjal Jain, Ashraf Mahgoub, Zihan Zhou, Kanak Mahadik, and Somali Chaterji. Lerna: Transformer architectures for configuring error correction tools for short- and long-read genome sequencing. BMC Bioinformatics, 23(1):25, 2022.

[65] Leena Salmela and Jan Schröder. Correcting errors in short reads by multiple alignments. Bioinformatics, 27(11):1455–1461, 2011.

[66] Wei-Chun Kao, Andrew H. Chan, and Yun S. Song. ECHO: A reference-free short-read error correction algorithm. Genome Research, 21(7):1181–1192, 2011.

[67] Marcel H. Schulz, David Weese, Manuel Holtgrewe, Viktoria Dimitrova, Sijia Niu, Knut Reinert, and Hugues Richard. Fiona: A parallel and automatic strategy for read error correction. Bioinformatics, 30(17):i356–i363, 2014.

[68] Amin Allam, Panos Kalnis, and Victor Solovyev. Karect: Accurate correction of substitution, insertion and deletion errors for next-generation sequencing data. Bioinformatics, 31(21):3421–3428, 2015.

[69] Felix Kallenborn, Andreas Hildebrandt, and Bertil Schmidt. CARE: Context-aware sequencing read error correction. Bioinformatics, 37(7):889–895, 2021.

[70] Felix Kallenborn, Julian Cascitti, and Bertil Schmidt. CARE 2.0: Reducing false-positive sequencing error corrections using machine learning. BMC Bioinformatics, 23(1):227, 2022.

[71] Mahdi Heydari, Giles Miclotte, Yves Van de Peer, and Jan Fostier. Illumina error correction near highly repetitive DNA regions improves de novo genome assembly. BMC Bioinformatics, 20(1):298, 2019.

[72] Antoine Limasset, Jean-François Flot, and Pierre Peterlongo. Toward perfect reads: Self-correction of short reads via mapping on de Bruijn graphs. Bioinformatics, 36(2):651, 2020.

[73] Xuan Zhang, Yuansheng Liu, Zuguo Yu, Michael Blumenstein, Gyorgy Hutvagner, and Jinyan Li. Instance-based error correction for short reads of disease-associated genes. BMC Bioinformatics, 22(6):142, 2021.

[74] Xuan Zhang, Pengyao Ping, Gyorgy Hutvagner, Michael Blumenstein, and Jinyan Li. Aberration-corrected ultrafine analysis of miRNA reads at single-base resolution: A k-mer lattice approach. Nucleic Acids Research, 49(18):e106, 2021.

[75] Eric M. Davis, Yu Sun, Yanling Liu, Pandurang Kolekar, Ying Shao, Karol Szlachta, Heather L. Mulder, Dongren Ren, Stephen V. Rice, Zhaoming Wang, Joy Nakitandwe, Alexander M. Gout, Bridget Shaner, Salina Hall, Leslie L. Robison, Stanley Pounds, Jeffery M. Klco, John Easton, and Xiaotu Ma. SequencErr: Measuring and suppressing sequencer errors in next-generation sequencing data. Genome Biology, 22(1):37, 2021.

