## Supplementary Materials for "How Error Correction Affects PCR Deduplication: A Survey Based on UMI Datasets of Short Reads"

**Supplementary materials, including Supplementary Figures 1-62 and Supplementary Tables 1-2, are extended data supporting the analysis in this review.**

#### 1 Supplementary Figures

- Supplementary Figure 1 compares the overlap and differences between unique read sets after deduplication by UMI-based methods of UMI-tools, AmpUMI and UMIc using Venn diagrams on the data sets of SRR1543965-SRR1543971, SRR28313990 and SRR28314008.
- Supplementary Figures 2-10 are line charts for comparing overlapped reads numbers between each of the computational methods of NGSReadsTreatment, Nubeam-dedup, BioSeqZip, fastp, FastUniq, pRESTO, CD-HIT-DUP, ParDRe and Minirmd with each of the UMI-based methods of UMI-tools, AmpUMI and UMIc on the datasets of SRR1543965-SRR1543971, SRR28313990 and SRR28314008.
- Supplementary Figures 11-20 are heatmaps for comparing overlapped reads numbers among each of the computational methods of NGSReadsTreatment, Nubeam-dedup, BioSeqZip, fastp, FastUniq, pRESTO, CD-HIT-DUP, ParDRe and Minirmd with the UMI-based methods of AmpUMI and UMIc on the data sets of SRR1543965-SRR1543971, SRR28313972, SRR28313990 and SRR28314008. The number immediately following a method indicates the mismatch values allowed by that method.
- Supplementary Figures 21-31 compare the overlaps and differences between the unique read sets obtained by different error correction algorithms using UpSet plots on data sets SRR1543964-SRR1543971, SRR28313972, SRR28313990 and SRR28314008. These figures are long-scale pictures and their high-resolution versions are presented in the attachment separately in the form of png.
- Supplementary Figures 32-62 are line charts for comparing the number of overlapped reads between deduplicated read set by CD-HIT-DUP, ParDRe and Minirmd with mismatches ranging from 1 to 3 on error-corrected dataset SRR1543964-SRR1543971, SRR28313972, SRR28313990 and SRR28314008 with deduplicated read set by each of the UMI-based methods of AmpUMI, UMI-tools or UMIc. BFC, Bcool, Care, Coral, Fiona, Lighter, Pollux and RACER were used for error correction, respectively. The dashed line labelled "Mismatch=0" was obtained by CD-HIT-DUP with setting mismatch as 0.

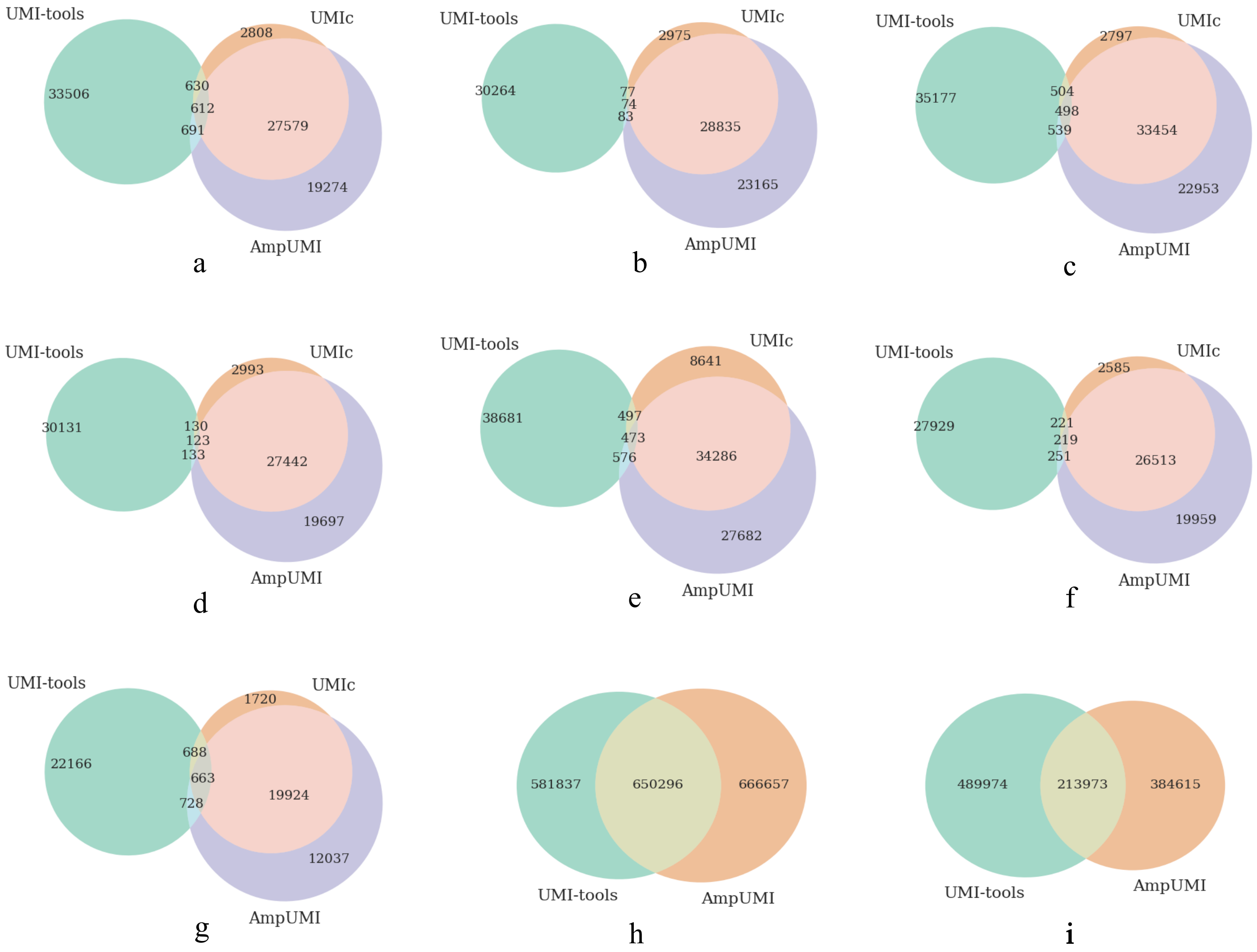


Supplementary Figure 1: The performance comparison of the overlap and differences between unique read sets after deduplication by UMI-based methods of UMI-tools, AmpUMI and UMIc using Venn diagrams on the data sets of SRR1543965-SRR1543971, SRR28313990 and SRR28314008.

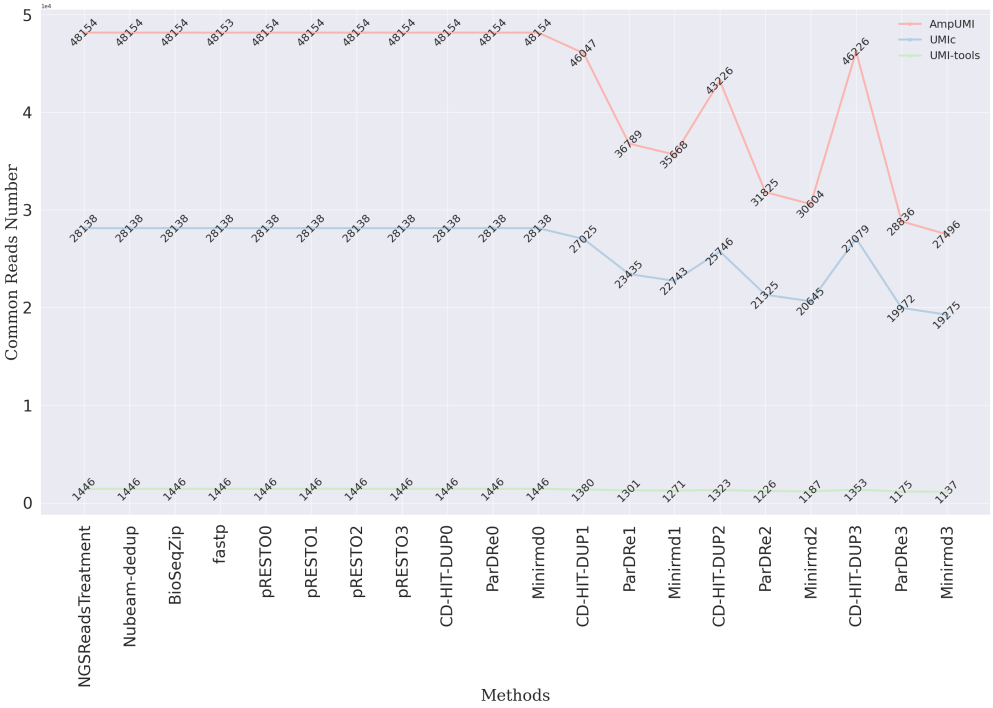

Supplementary Figure 2: Line chart for comparing overlapped reads number between each of the computational methods of NGSReadsTreatment, Nubeam-dedup, BioSeqZip, fastp, FastUniq, pRESTO, CD-HIT-DUP, ParDRe and Minirmd with each of the UMI-based methods of UMI-tools, AmpUMI and UMIc on the data set SRR1543965.

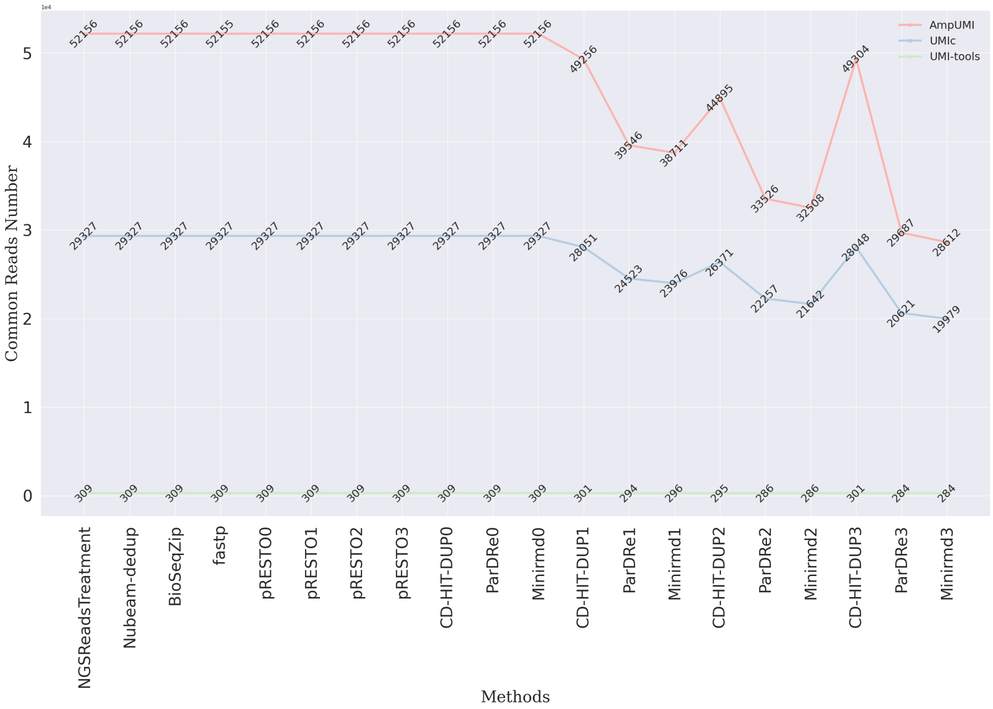

Supplementary Figure 3: Line chart for comparing overlapped reads number between each of the computational methods of NGSReadsTreatment, Nubeam-dedup, BioSeqZip, fastp, FastUniq, pRESTO, CD-HIT-DUP, ParDRe and Minirmd with each of the UMI-based methods of UMI-tools, AmpUMI and UMIc on the data set SRR1543966.

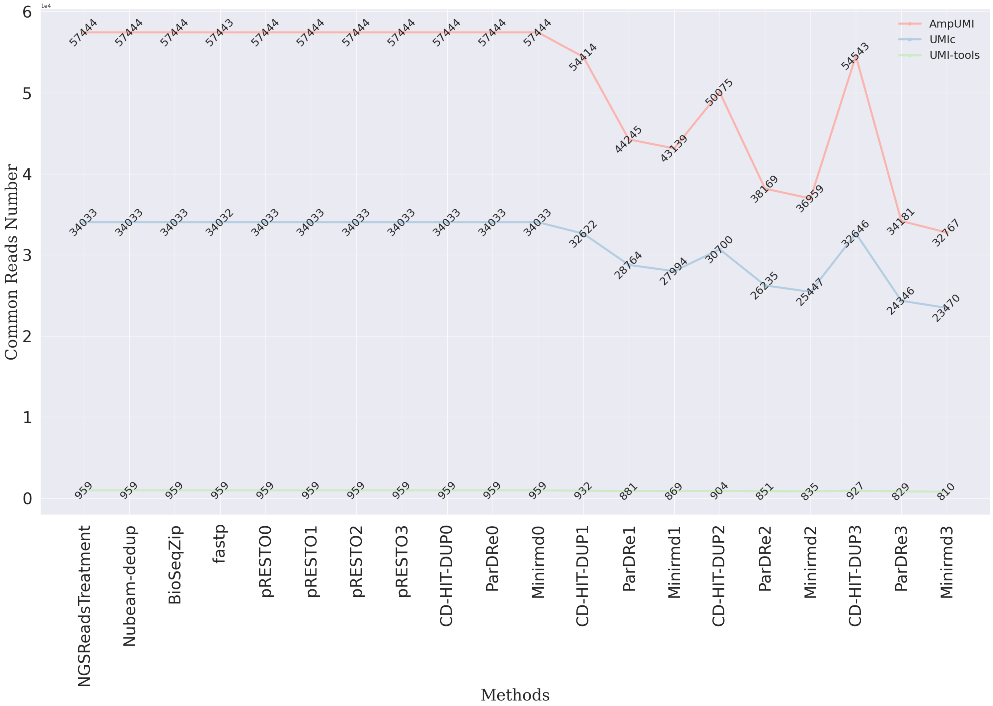


Supplementary Figure 4: Line chart for comparing overlapped reads number between each of the computational methods of NGSReadsTreatment, Nubeam-dedup, BioSeqZip, fastp, FastUniq, pRESTO, CD-HIT-DUP, ParDRe and Minirmd with each of the UMI-based methods of UMI-tools, AmpUMI and UMIc on the data set SRR1543967.

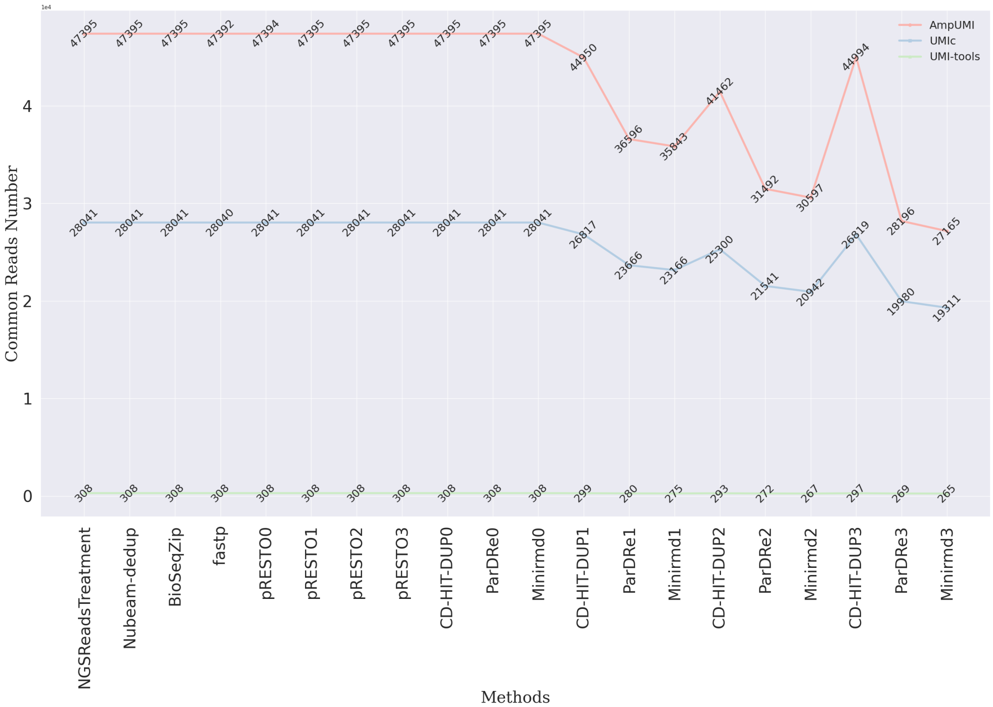


Supplementary Figure 5: Line chart for comparing overlapped reads number between each of the computational methods of NGSReadsTreatment, Nubeam-dedup, BioSeqZip, fastp, FastUniq, pRESTO, CD-HIT-DUP, ParDRe and Minirmd with each of the UMI-based methods of UMI-tools, AmpUMI and UMIc on the data set SRR1543968.

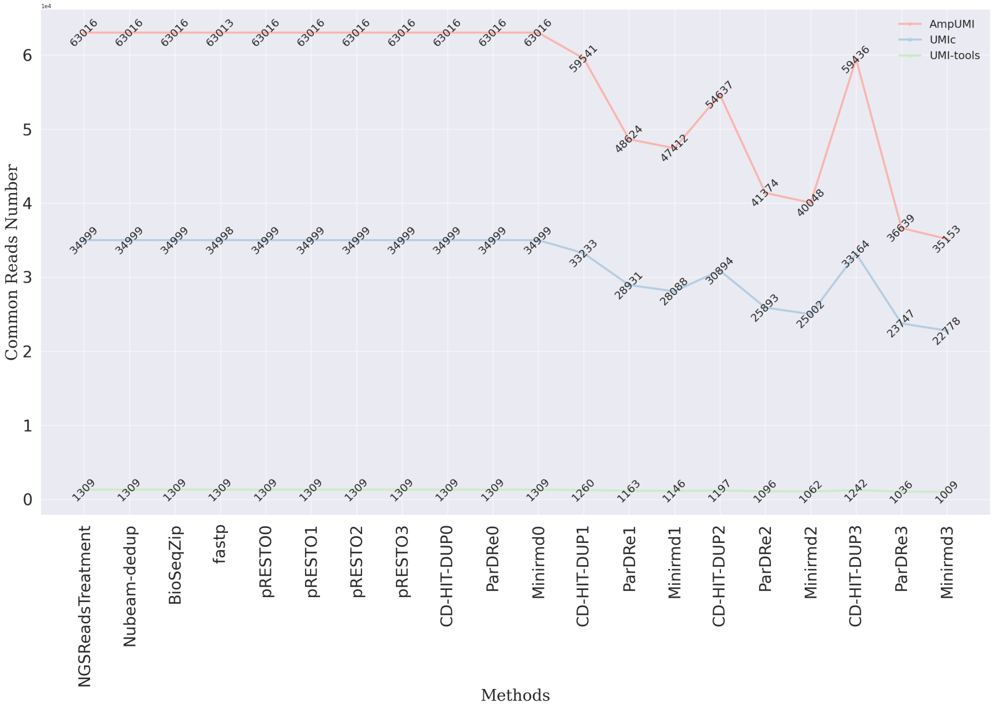


Supplementary Figure 6: Line chart for comparing overlapped reads number between each of the computational methods of NGSReadsTreatment, Nubeam-dedup, BioSeqZip, fastp, FastUniq, pRESTO, CD-HIT-DUP, ParDRe and Minirmd with each of the UMI-based methods of UMI-tools, AmpUMI and UMIc on the data set SRR1543969.

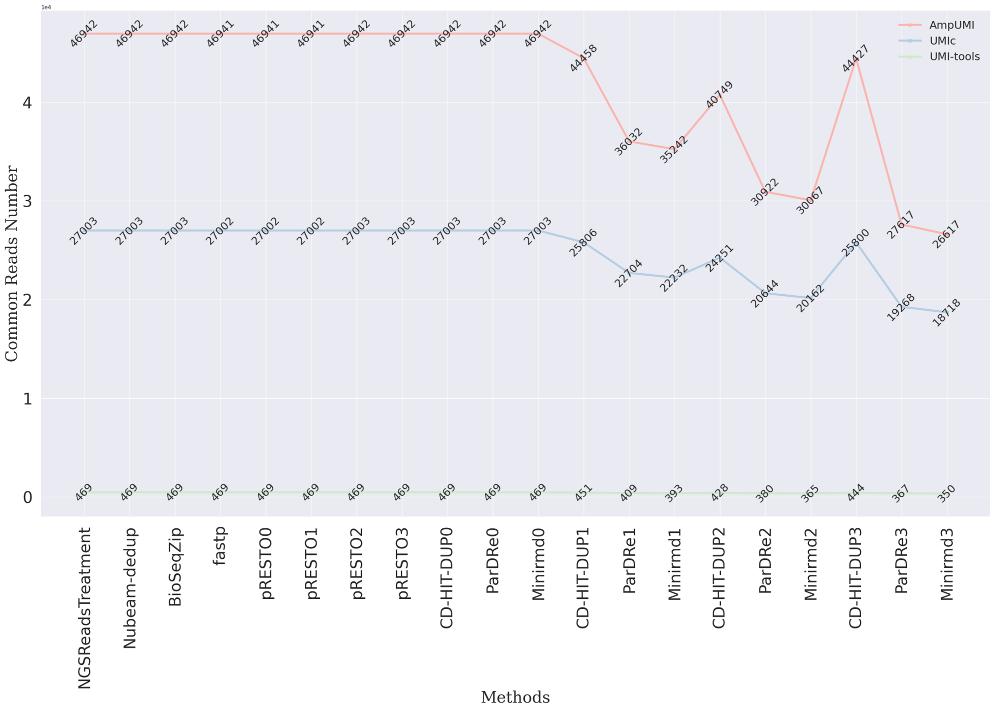


Supplementary Figure 7: Line chart for comparing overlapped reads number between each of the computational methods of NGSReadsTreatment, Nubeam-dedup, BioSeqZip, fastp, FastUniq, pRESTO, CD-HIT-DUP, ParDRe and Minirmd with each of the UMI-based methods of UMI-tools, AmpUMI and UMIc on the data set SRR1543970.

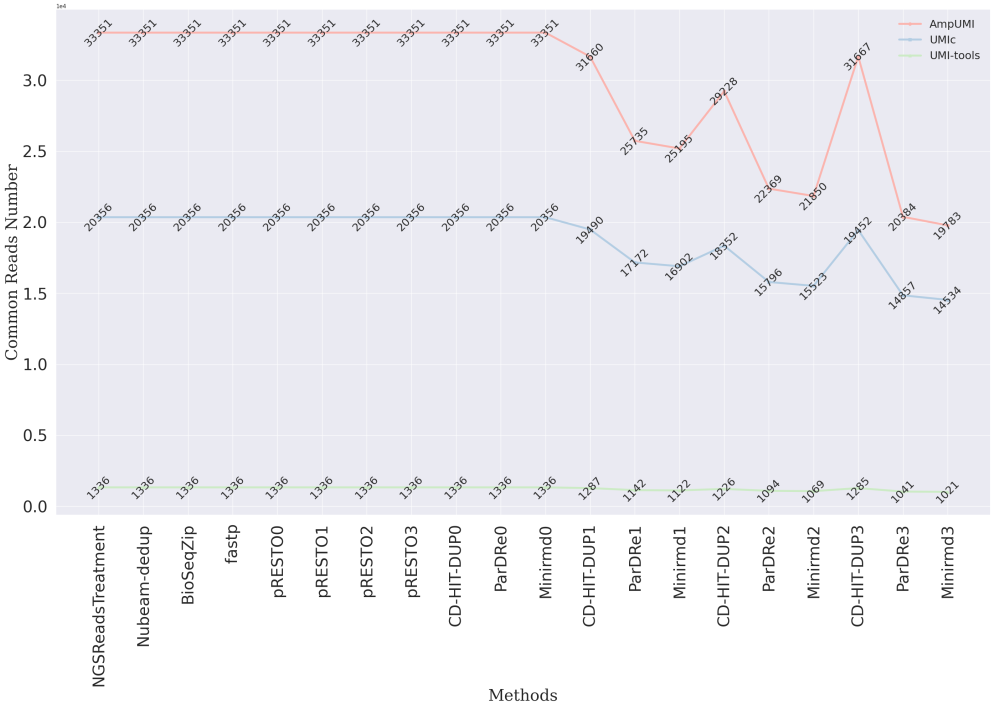


Supplementary Figure 8: Line chart for comparing overlapped reads number between each of the computational methods of NGSReadsTreatment, Nubeam-dedup, BioSeqZip, fastp, FastUniq, pRESTO, CD-HIT-DUP, ParDRe and Minirmd with each of the UMI-based methods of UMI-tools, AmpUMI and UMIc on the data set SRR1543971.

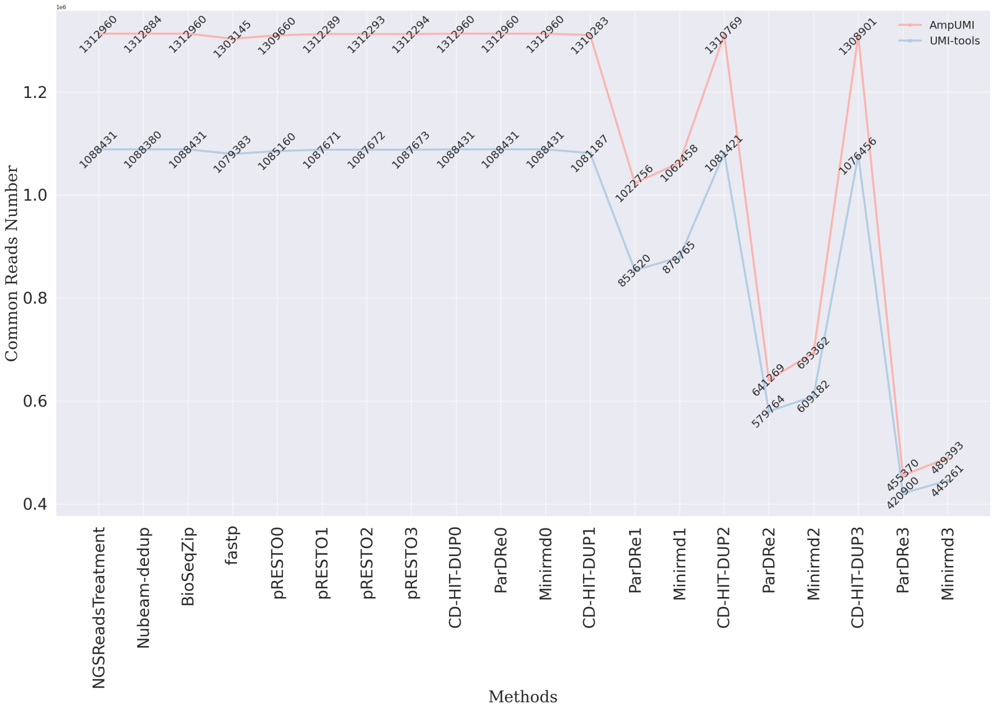


Supplementary Figure 9: Line chart for comparing overlapped reads number between each of the computational methods of NGSReadsTreatment, Nubeam-dedup, BioSeqZip, fastp, FastUniq, pRESTO, CD-HIT-DUP, ParDRe and Minirmd with each of the UMI-based methods of UMI-tools and AmpUMI on the dataset SRR28313990.

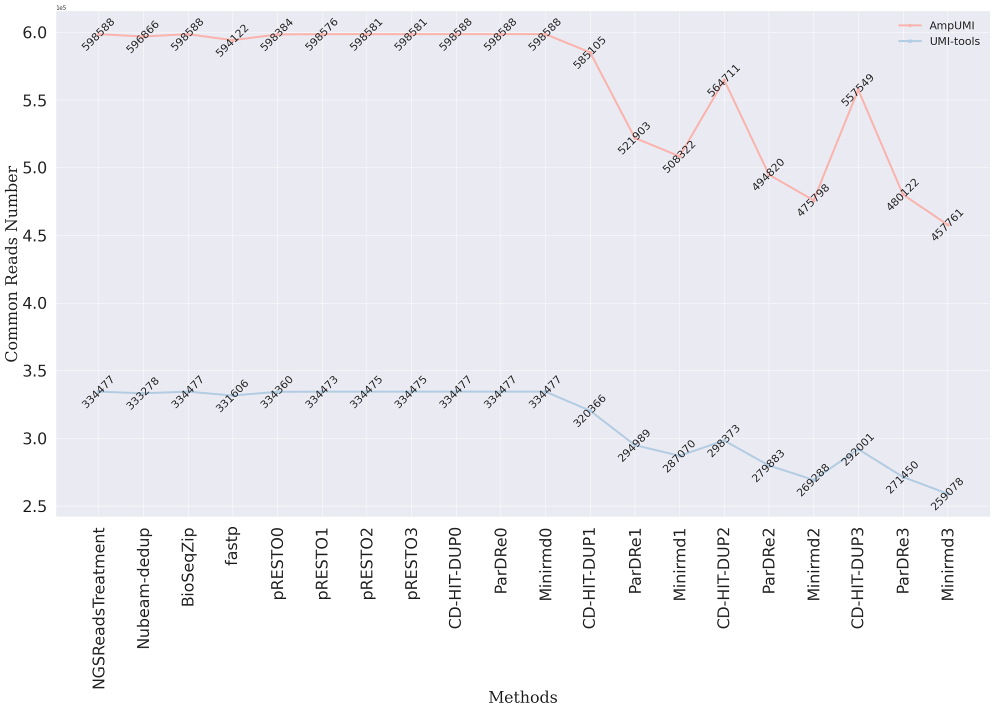


Supplementary Figure 10: Line chart for comparing overlapped reads number between each of the computational methods of NGSReadsTreatment, Nubeam-dedup, BioSeqZip, fastp, FastUniq, pRESTO, CD-HIT-DUP, ParDRe and Minirmd with each of the UMI-based methods of UMI-tools and AmpUMI on the dataset SRR28314008.

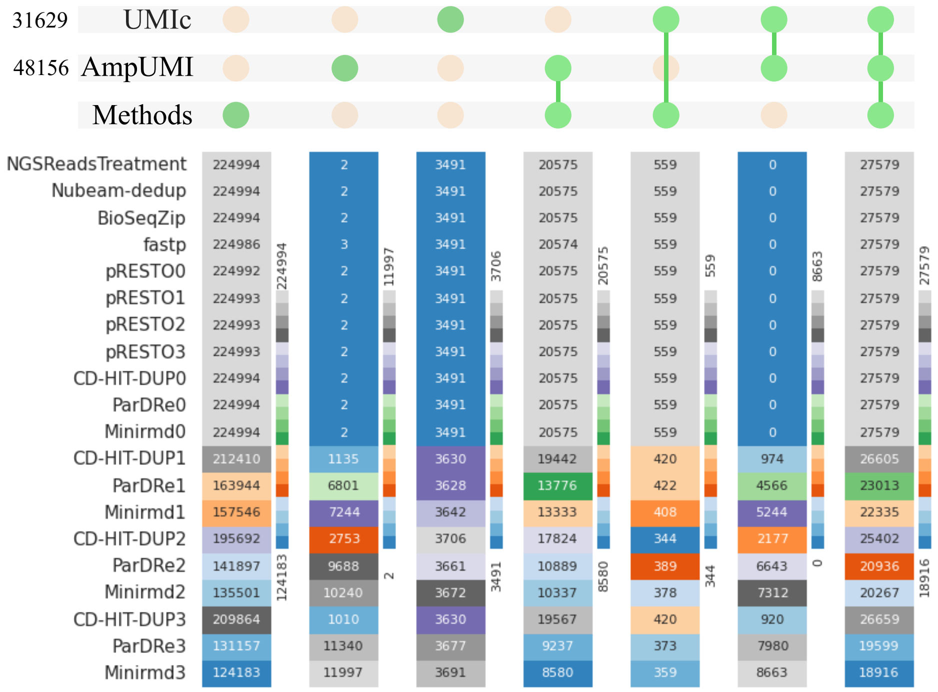


Supplementary Figure 11: Heatmap for comparing overlapped reads number by each of the computational methods of NGSReadsTreatment, Nubeam-dedup, BioSeqZip, fastp, FastUniq, pRESTO, CD-HIT-DUP, ParDRe and Minirmd with the UMI-based methods of AmpUMI and UMIc on the data set SRR1543965.

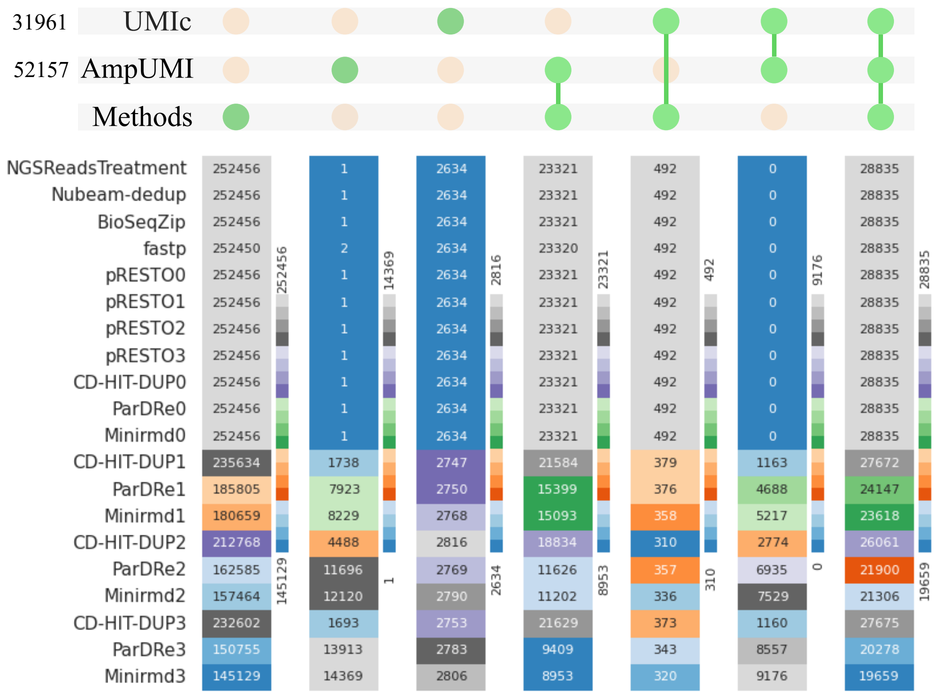


Supplementary Figure 12: Heatmap for comparing overlapped reads number by each of the computational methods of NGSReadsTreatment, Nubeam-dedup, BioSeqZip, fastp, FastUniq, pRESTO, CD-HIT-DUP, ParDRe and Minirmd with the UMI-based methods of AmpUMI and UMIc on the data set SRR1543966.

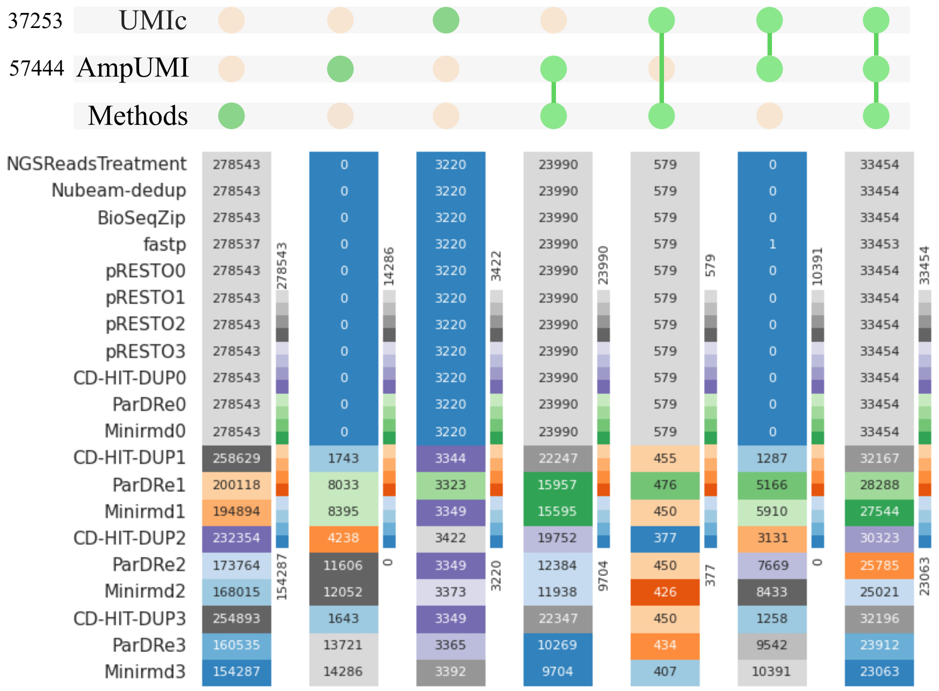


Supplementary Figure 13: Heatmap for comparing overlapped reads number by each of the computational methods of NGSReadsTreatment, Nubeam-dedup, BioSeqZip, fastp, FastUniq, pRESTO, CD-HIT-DUP, ParDRe and Minirmd with the UMI-based methods of AmpUMI and UMIc on the data set SRR1543967.

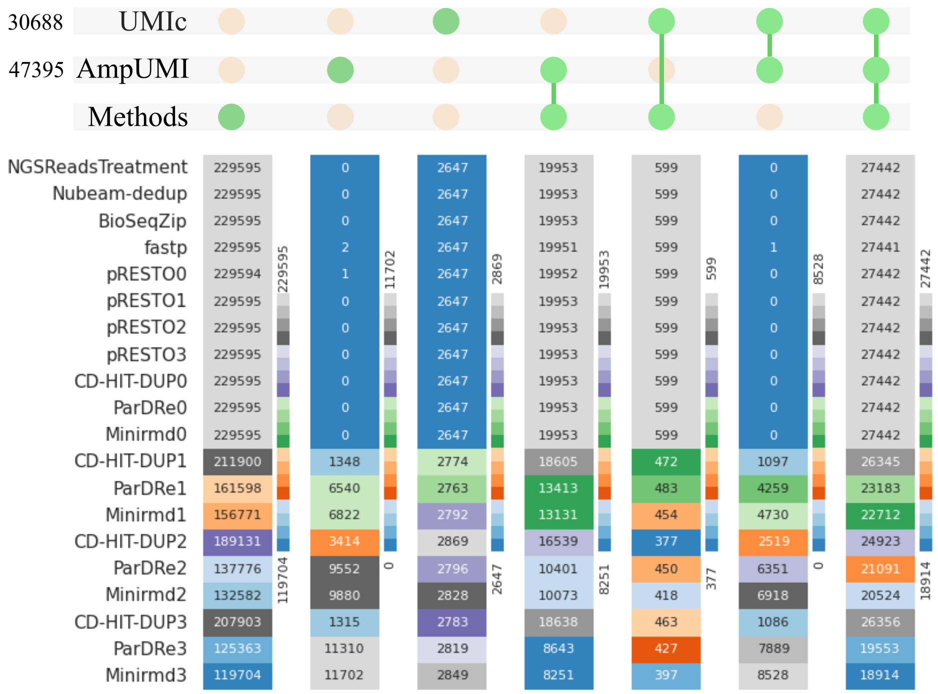


Supplementary Figure 14: Heatmap for comparing overlapped reads number by each of the computational methods of NGSReadsTreatment, Nubeam-dedup, BioSeqZip, fastp, FastUniq, pRESTO, CD-HIT-DUP, ParDRe and Minirmd with the UMI-based methods of AmpUMI and UMIc on the data set SRR1543968.

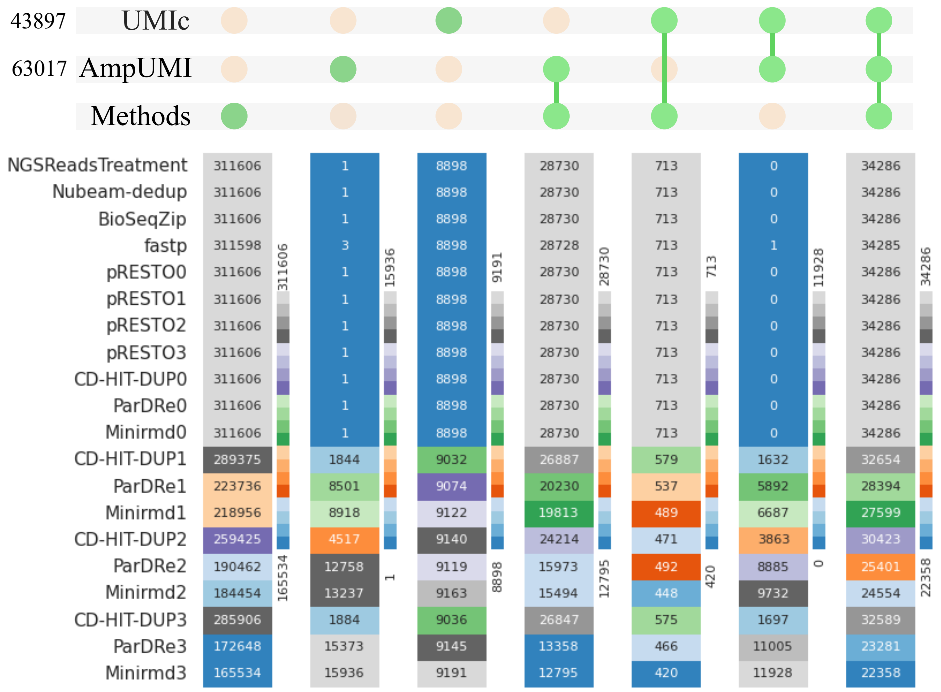


Supplementary Figure 15: Heatmap for comparing overlapped reads number by each of the computational methods of NGSReadsTreatment, Nubeam-dedup, BioSeqZip, fastp, FastUniq, pRESTO, CD-HIT-DUP, ParDRe and Minirmd with the UMI-based methods of AmpUMI and UMIc on the data set SRR1543969.

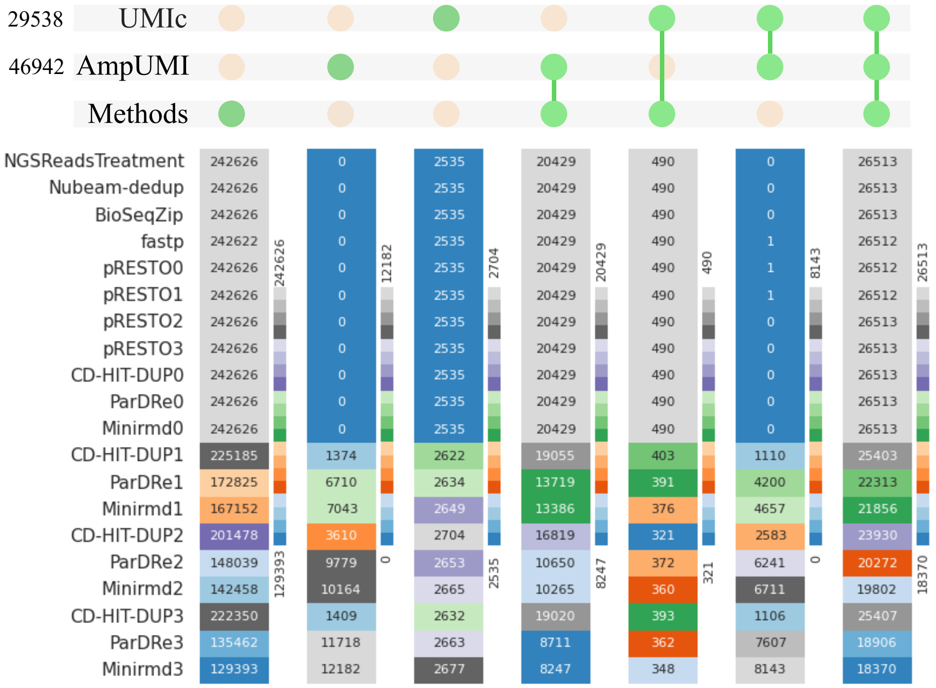


Supplementary Figure 16: Heatmap for comparing overlapped reads number by each of the computational methods of NGSReadsTreatment, Nubeam-dedup, BioSeqZip, fastp, FastUniq, pRESTO, CD-HIT-DUP, ParDRe and Minirmd with the UMI-based methods of AmpUMI and UMIc on the data set SRR1543970.

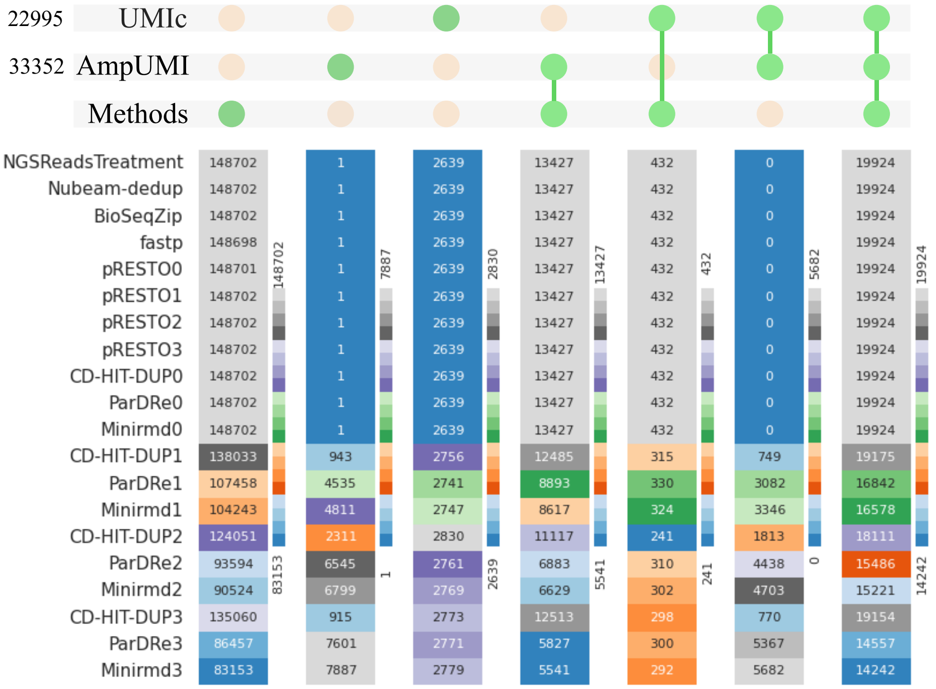


Supplementary Figure 17: Heatmap for comparing overlapped reads number by each of the computational methods of NGSReadsTreatment, Nubeam-dedup, BioSeqZip, fastp, FastUniq, pRESTO, CD-HIT-DUP, ParDRe and Minirmd with the UMI-based methods of AmpUMI and UMIc on the data set SRR1543971.

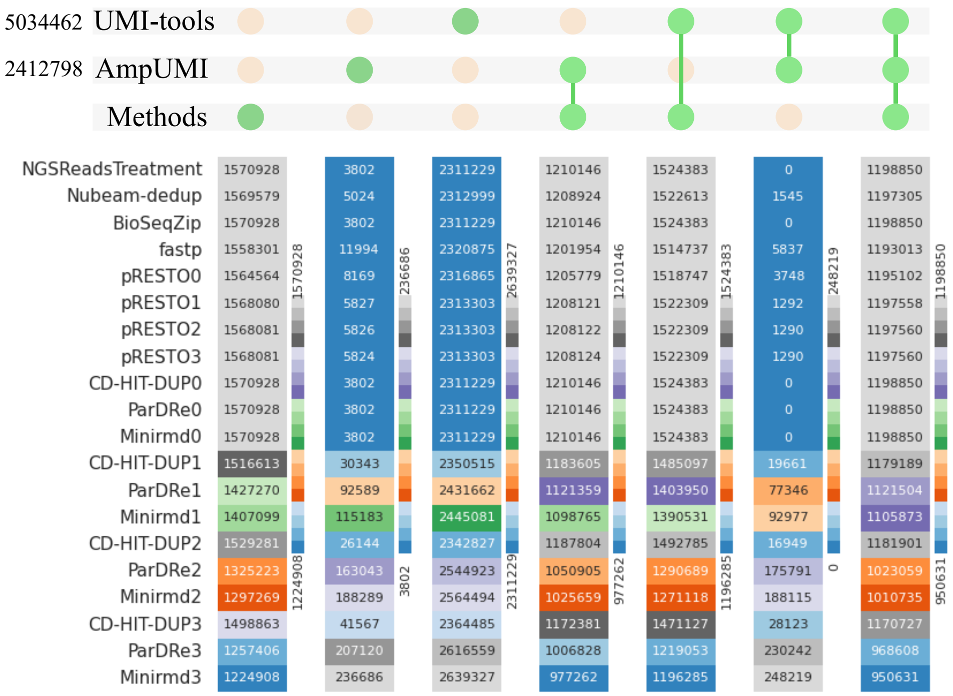


Supplementary Figure 18: Heatmap for comparing overlapped reads number by each of the computational methods of NGSReadsTreatment, Nubeam-dedup, BioSeqZip, fastp, FastUniq, pRESTO, CD-HIT-DUP, ParDRe and Minirmd with the UMI-based methods of AmpUMI and UMI-tools on the data set SRR28313972.

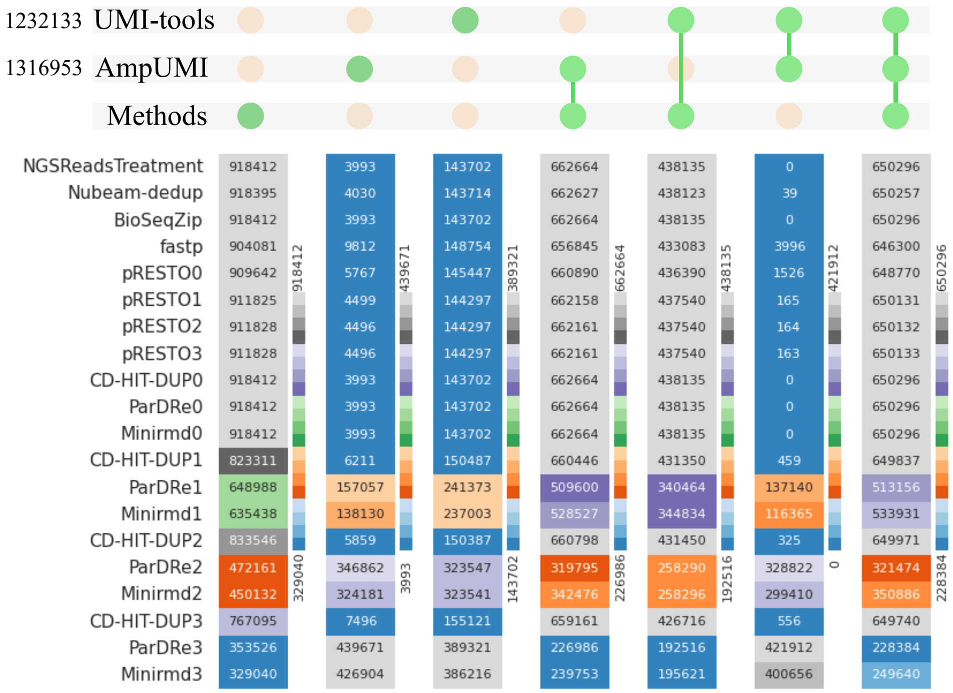


Supplementary Figure 19: Heatmap for comparing overlapped reads number by each of the computational methods of NGSReadsTreatment, Nubeam-dedup, BioSeqZip, fastp, FastUniq, pRESTO, CD-HIT-DUP, ParDRe and Minirmd with the UMI-based methods of AmpUMI and UMI-tools on the data set SRR28313990.

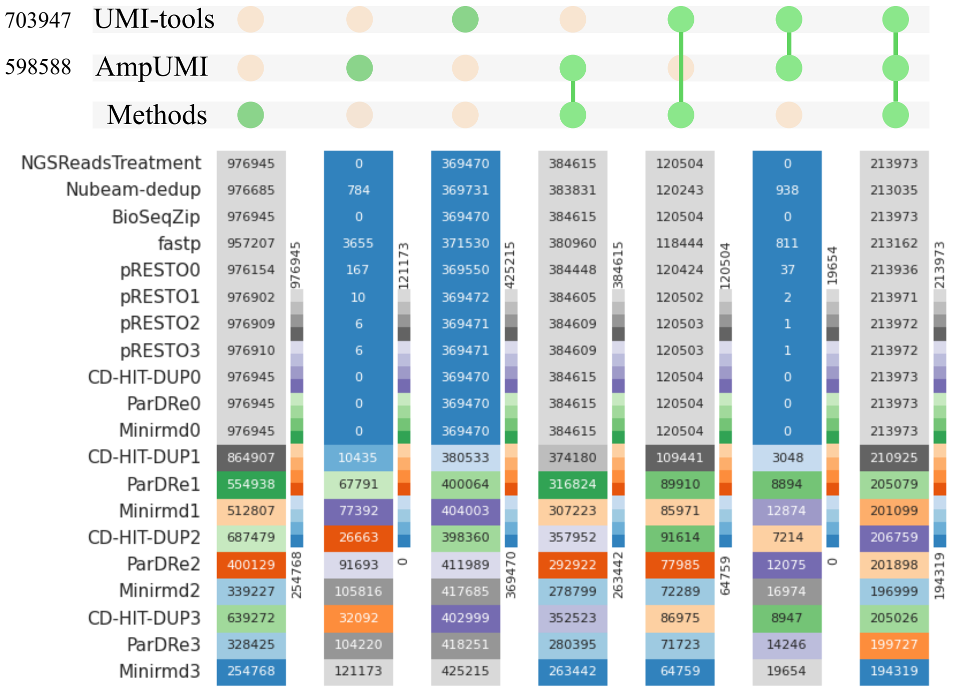


Supplementary Figure 20: Heatmap for comparing overlapped reads number by each of the computational methods of NGSReadsTreatment, Nubeam-dedup, BioSeqZip, fastp, FastUniq, pRESTO, CD-HIT-DUP, ParDRe and Minirmd with the UMI-based methods of AmpUMI and UMI-tools on the data set SRR28314008.

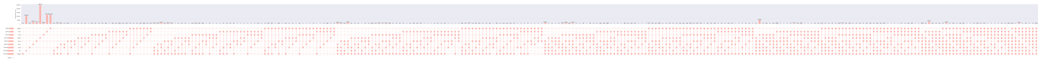

Supplementary Figure 21: The overlaps and differences comparison between the unique read sets obtained by different error correction algorithms using UpSet plots on data sets SRR1543964. The original high-resolution figure is a long-scale picture presented in the attachment separately in png format.

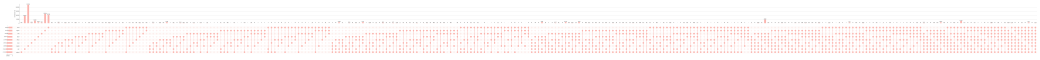

Supplementary Figure 22: The overlaps and differences comparison between the unique read sets obtained by different error correction algorithms using UpSet plots on data sets SRR1543965. The original high-resolution figure is a long-scale picture presented in the attachment separately in png format.

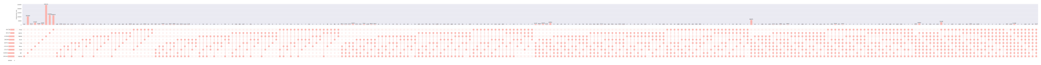

Supplementary Figure 23: The overlaps and differences comparison between the unique read sets obtained by different error correction algorithms using UpSet plots on data sets SRR1543966. The original high-resolution figure is a long-scale picture presented in the attachment separately in png format.

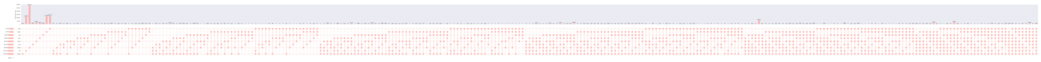

Supplementary Figure 24: The overlaps and differences comparison between the unique read sets obtained by different error correction algorithms using UpSet plots on data sets SRR1543967. The original high-resolution figure is a long-scale picture presented in the attachment separately in png format.

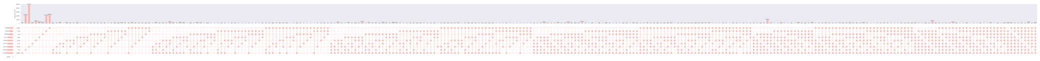

Supplementary Figure 25: The overlaps and differences comparison between the unique read sets obtained by different error correction algorithms using UpSet plots on data sets SRR1543968. The original high-resolution figure is a long-scale picture presented in the attachment separately in png format.

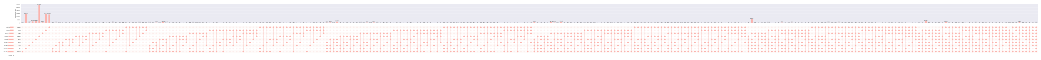

Supplementary Figure 26: The overlaps and differences comparison between the unique read sets obtained by different error correction algorithms using UpSet plots on data sets SRR1543969. The original high-resolution figure is a long-scale picture presented in the attachment separately in png format.

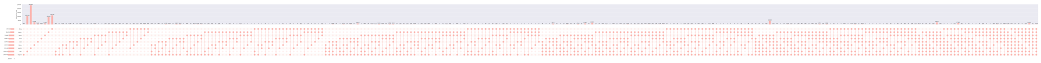

Supplementary Figure 27: The overlaps and differences comparison between the unique read sets obtained by different error correction algorithms using UpSet plots on data sets SRR1543970. The original high-resolution figure is a long-scale picture presented in the attachment separately in png format.

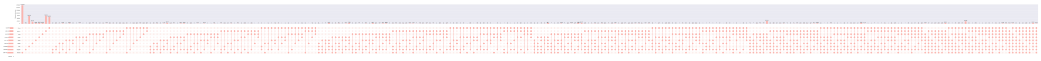

Supplementary Figure 28: The overlaps and differences comparison between the unique read sets obtained by different error correction algorithms using UpSet plots on data sets SRR1543971. The original figure is a long-scale picture presented in the attachment separately in png format.

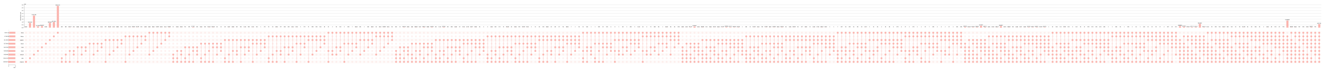

Supplementary Figure 29: The overlaps and differences comparison between the unique read sets obtained by different error correction algorithms using UpSet plots on data sets SRR28313972. The original figure is a long-scale picture presented in the attachment separately in png format.

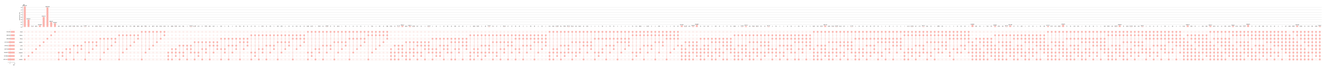

Supplementary Figure 30: The overlaps and differences comparison between the unique read sets obtained by different error correction algorithms using UpSet plots on data sets SRR28313990. The original figure is a long-scale picture presented in the attachment separately in png format.

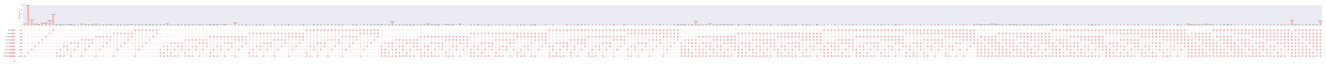

Supplementary Figure 31: The overlaps and differences comparison between the unique read sets obtained by different error correction algorithms using UpSet plots on data sets SRR28314008. The original figure is a long-scale picture presented in the attachment separately in png format.

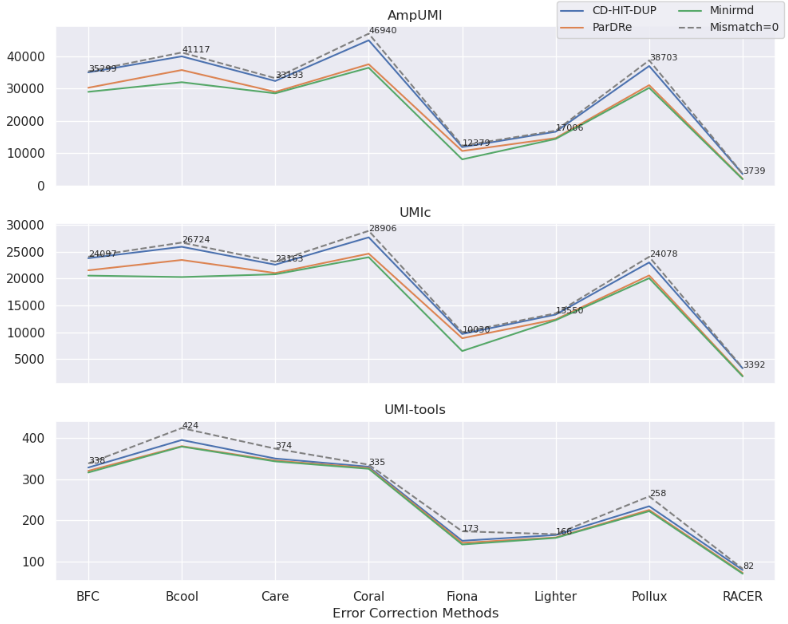


Supplementary Figure 32: Line charts comparing the overlapped read numbers in the deduplicated read set by the PCR-deduplication methods of CD-HIT-DUP, ParDRe, and Minirmd on the error-corrected dataset SRR1543964, with each UMI-based PCR-deduplication methods of UMI-tools, AmpUMI, and UMIc on dataset SRR1543964. Error correction was performed using error-correction methods of BFC, Bcool, Care, Coral, Fiona, Lighter, Pollux, and RACER, respectively. CD-HIT-DUP, ParDRe, and Minirmd employed a mismatched number set to 1. The dashed line labelled 'Mismatch=0' represents results obtained by CD-HIT-DUP with a mismatch setting of 0.

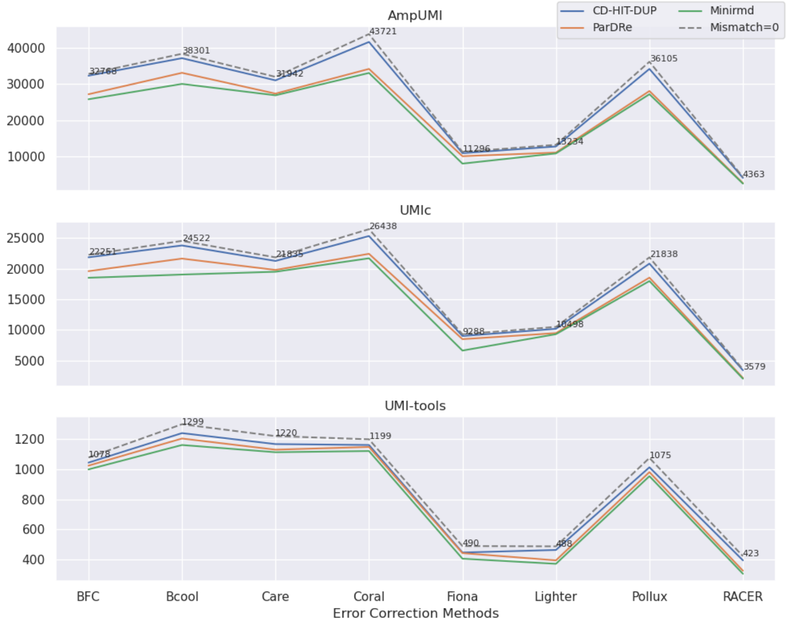


Supplementary Figure 33: Line charts comparing the overlapped read numbers in the deduplicated read set by the PCR-deduplication methods of CD-HIT-DUP, ParDRe, and Minirmd on the error-corrected dataset SRR1543965, with each UMI-based PCR-deduplication methods of UMI-tools, AmpUMI, and UMIc on dataset SRR1543965. Error correction was performed using error-correction methods of BFC, Bcool, Care, Coral, Fiona, Lighter, Pollux, and RACER, respectively. CD-HIT-DUP, ParDRe, and Minirmd employed a mismatched number set to 1. The dashed line labelled 'Mismatch=0' represents results obtained by CD-HIT-DUP with a mismatch setting of 0.

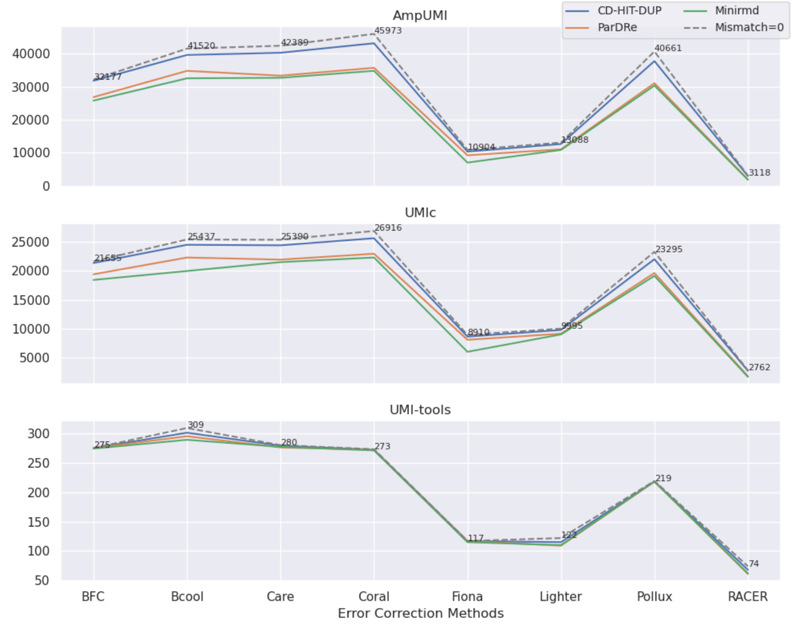


Supplementary Figure 34: Line charts comparing the overlapped read numbers in the deduplicated read set by the PCR-deduplication methods of CD-HIT-DUP, ParDRe, and Minirmd on the error-corrected dataset SRR1543966, with each UMI-based PCR-deduplication methods of UMI-tools, AmpUMI, and UMIc on dataset SRR1543966. Error correction was performed using error-correction methods of BFC, Bcool, Care, Coral, Fiona, Lighter, Pollux, and RACER, respectively. CD-HIT-DUP, ParDRe, and Minirmd employed a mismatched number set to 1. The dashed line labelled 'Mismatch=0' represents results obtained by CD-HIT-DUP with a mismatch setting of 0.

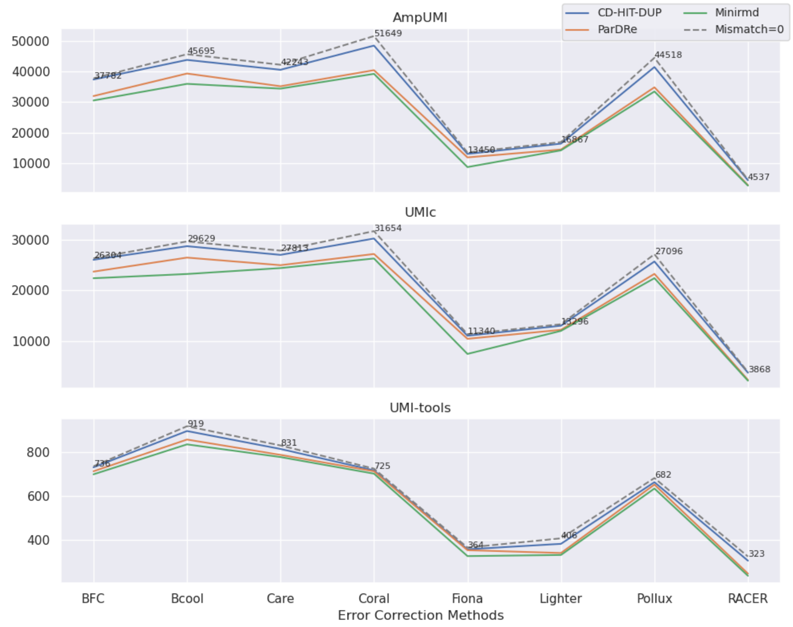


Supplementary Figure 35: Line charts comparing the overlapped read numbers in the deduplicated read set by the PCR-deduplication methods of CD-HIT-DUP, ParDRe, and Minirmd on the error-corrected dataset SRR1543967, with each UMI-based PCR-deduplication methods of UMI-tools, AmpUMI, and UMIc on dataset SRR1543967. Error correction was performed using error-correction methods of BFC, Bcool, Care, Coral, Fiona, Lighter, Pollux, and RACER, respectively. CD-HIT-DUP, ParDRe, and Minirmd employed a mismatched number set to 1. The dashed line labelled 'Mismatch=0' represents results obtained by CD-HIT-DUP with a mismatch setting of 0.

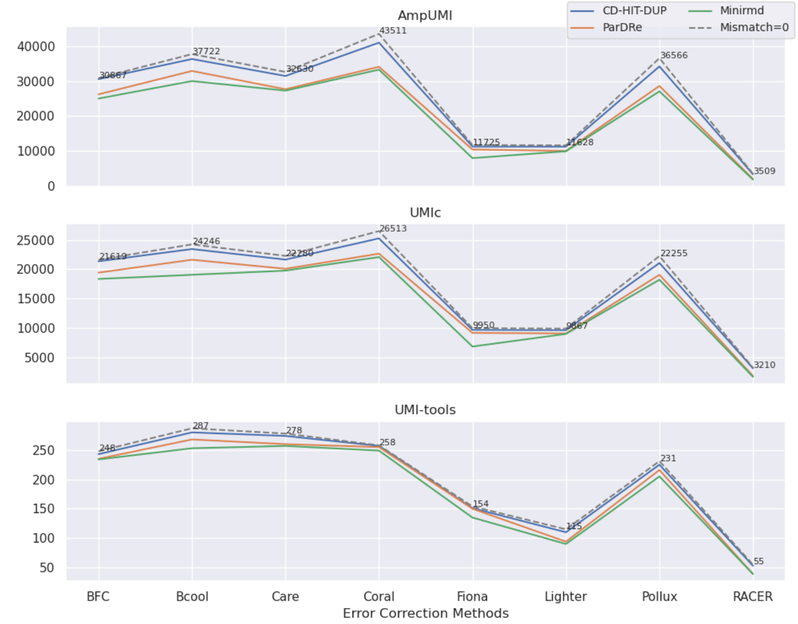


Supplementary Figure 36: Line charts comparing the overlapped read numbers in the deduplicated read set by the PCR-deduplication methods of CD-HIT-DUP, ParDRe, and Minirmd on the error-corrected dataset SRR1543968, with each UMI-based PCR-deduplication methods of UMI-tools, AmpUMI, and UMIc on dataset SRR1543968. Error correction was performed using error-correction methods of BFC, Bcool, Care, Coral, Fiona, Lighter, Pollux, and RACER, respectively. CD-HIT-DUP, ParDRe, and Minirmd employed a mismatched number set to 1. The dashed line labelled 'Mismatch=0' represents results obtained by CD-HIT-DUP with a mismatch setting of 0.

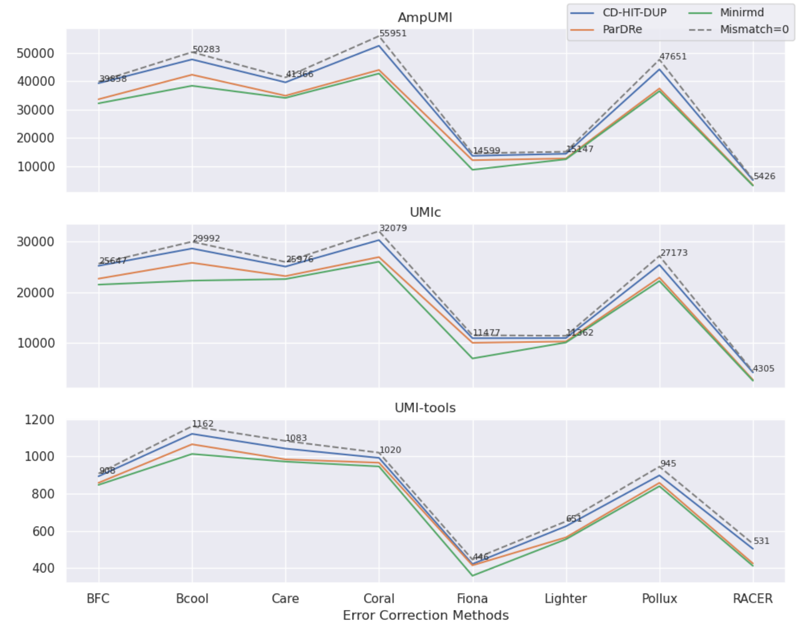


Supplementary Figure 37: Line charts comparing the overlapped read numbers in the deduplicated read set by the PCR-deduplication methods of CD-HIT-DUP, ParDRe, and Minirmd on the error-corrected dataset SRR1543969, with each UMI-based PCR-deduplication methods of UMI-tools, AmpUMI, and UMIc on dataset SRR1543969. Error correction was performed using error-correction methods of BFC, Bcool, Care, Coral, Fiona, Lighter, Pollux, and RACER, respectively. CD-HIT-DUP, ParDRe, and Minirmd employed a mismatched number set to 1. The dashed line labelled 'Mismatch=0' represents results obtained by CD-HIT-DUP with a mismatch setting of 0.

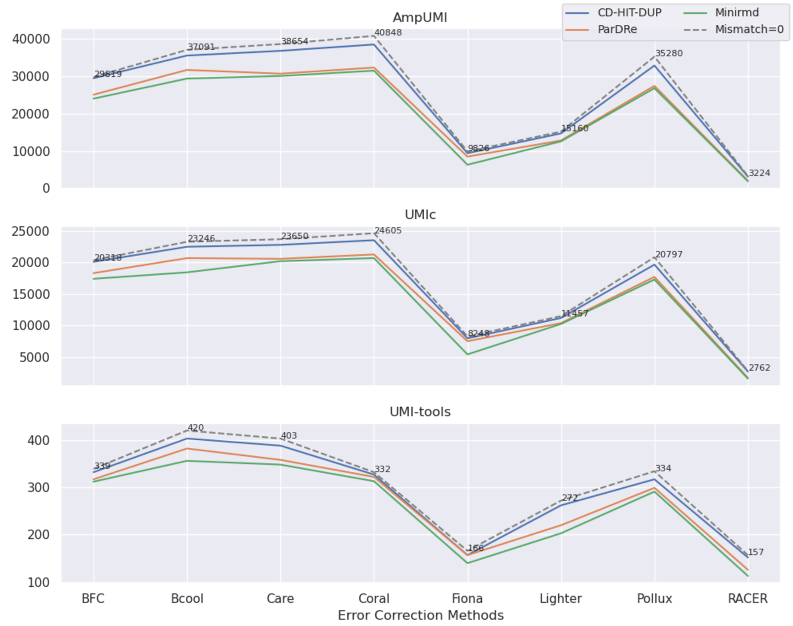


Supplementary Figure 38: Line charts comparing the overlapped read numbers in the deduplicated read set by the PCR-deduplication methods of CD-HIT-DUP, ParDRe, and Minirmd on the error-corrected dataset SRR1543970, with each UMI-based PCR-deduplication methods of UMI-tools, AmpUMI, and UMIc on dataset SRR1543970. Error correction was performed using error-correction methods of BFC, Bcool, Care, Coral, Fiona, Lighter, Pollux, and RACER, respectively. CD-HIT-DUP, ParDRe, and Minirmd employed a mismatched number set to 1. The dashed line labelled 'Mismatch=0' represents results obtained by CD-HIT-DUP with a mismatch setting of 0.

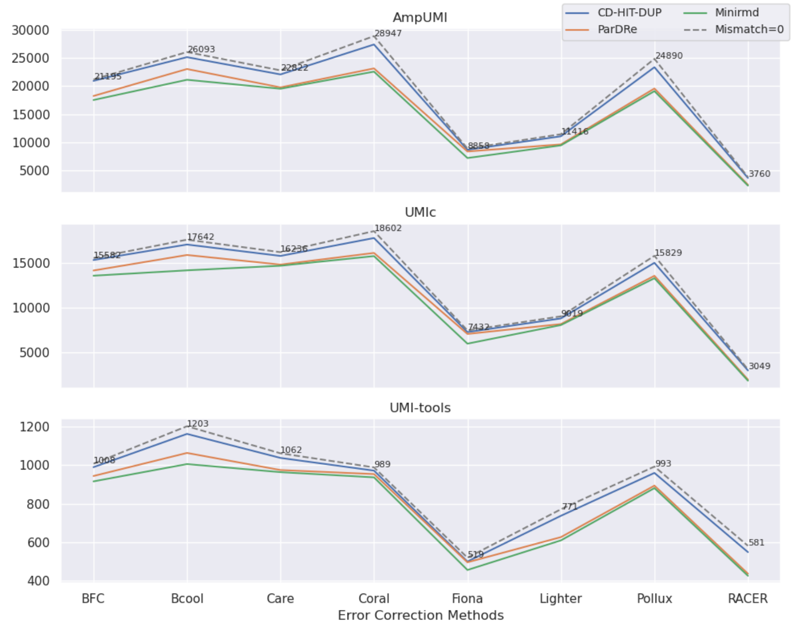


Supplementary Figure 39: Line charts comparing the overlapped read numbers in the deduplicated read set by the PCR-deduplication methods of CD-HIT-DUP, ParDRe, and Minirmd on the error-corrected dataset SRR1543971, with each UMI-based PCR-deduplication methods of UMI-tools, AmpUMI, and UMIc on dataset SRR1543971. Error correction was performed using error-correction methods of BFC, Bcool, Care, Coral, Fiona, Lighter, Pollux, and RACER, respectively. CD-HIT-DUP, ParDRe, and Minirmd employed a mismatched number set to 1. The dashed line labelled 'Mismatch=0' represents results obtained by CD-HIT-DUP with a mismatch setting of 0.

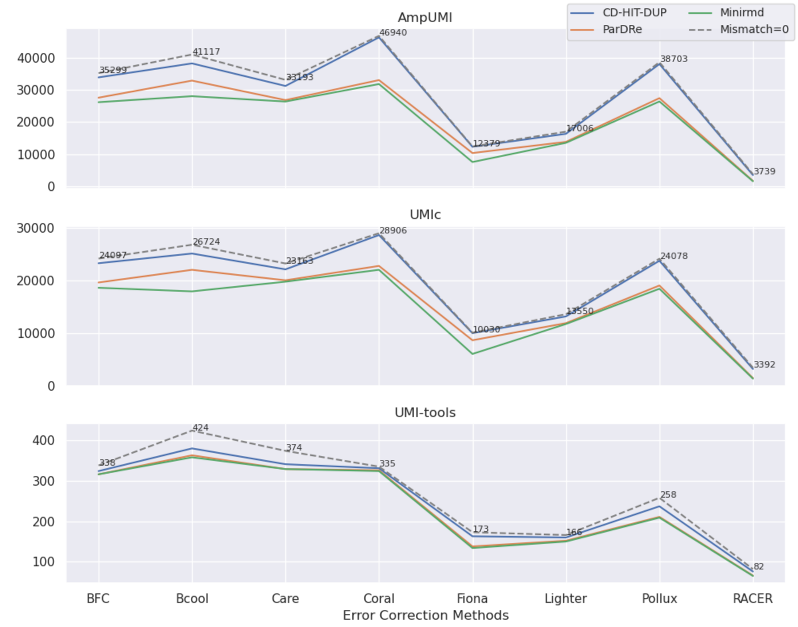


Supplementary Figure 40: Line charts comparing the overlapped read numbers in the deduplicated read set by the PCR-deduplication methods of CD-HIT-DUP, ParDRe, and Minirmd on the error-corrected dataset SRR1543964, with each UMI-based PCR-deduplication methods of UMI-tools, AmpUMI, and UMIc on dataset SRR1543964. Error correction was performed using error-correction methods of BFC, Bcool, Care, Coral, Fiona, Lighter, Pollux, and RACER, respectively. CD-HIT-DUP, ParDRe, and Minirmd employed a mismatched number set to 2. The dashed line labelled 'Mismatch=0' represents results obtained by CD-HIT-DUP with a mismatch setting of 0.

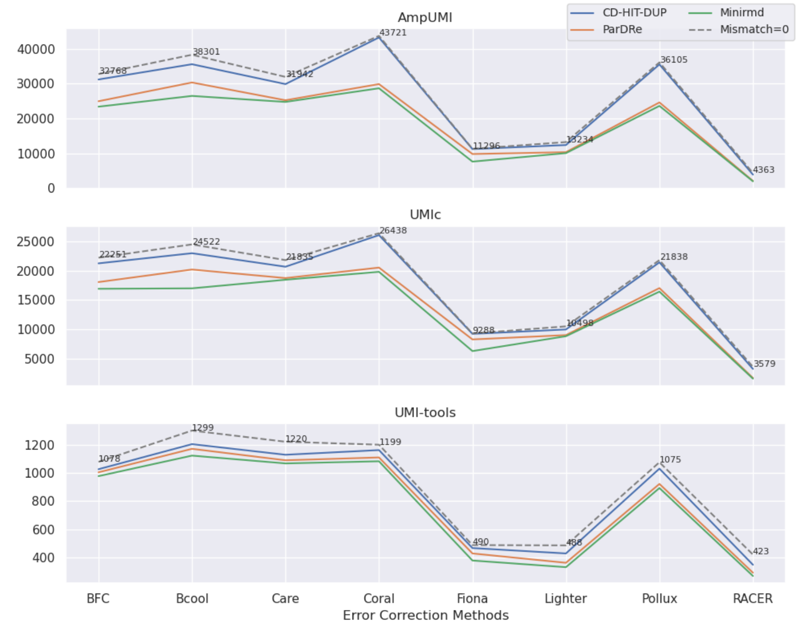


Supplementary Figure 41: Line charts comparing the overlapped read numbers in the deduplicated read set by the PCR-deduplication methods of CD-HIT-DUP, ParDRe, and Minirmd on the error-corrected dataset SRR1543965, with each UMI-based PCR-deduplication methods of UMI-tools, AmpUMI, and UMIc on dataset SRR1543965. Error correction was performed using error-correction methods of BFC, Bcool, Care, Coral, Fiona, Lighter, Pollux, and RACER, respectively. CD-HIT-DUP, ParDRe, and Minirmd employed a mismatched number set to 2. The dashed line labelled 'Mismatch=0' represents results obtained by CD-HIT-DUP with a mismatch setting of 0.

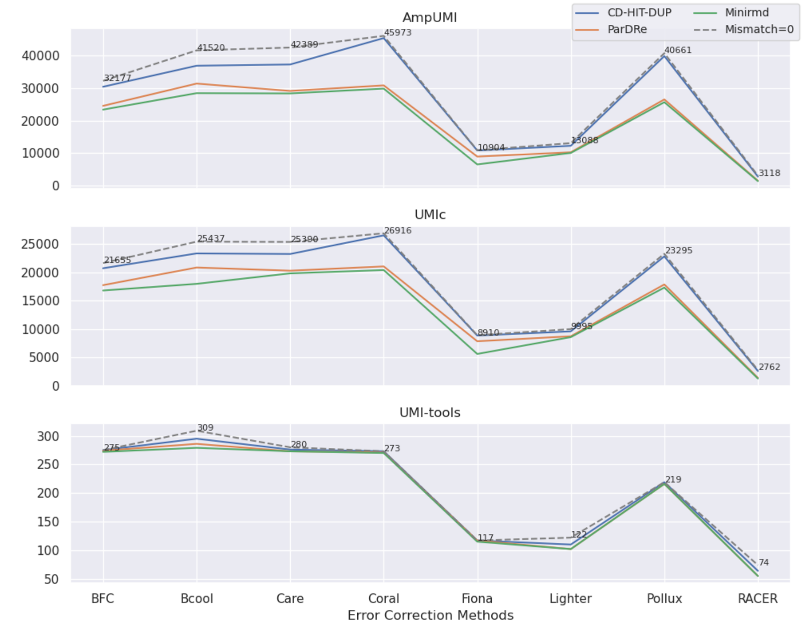


Supplementary Figure 42: Line charts comparing the overlapped read numbers in the deduplicated read set by the PCR-deduplication methods of CD-HIT-DUP, ParDRe, and Minirmd on the error-corrected dataset SRR1543966, with each UMI-based PCR-deduplication methods of UMI-tools, AmpUMI, and UMIc on dataset SRR1543966. Error correction was performed using error-correction methods of BFC, Bcool, Care, Coral, Fiona, Lighter, Pollux, and RACER, respectively. CD-HIT-DUP, ParDRe, and Minirmd employed a mismatched number set to 2. The dashed line labelled 'Mismatch=0' represents results obtained by CD-HIT-DUP with a mismatch setting of 0.

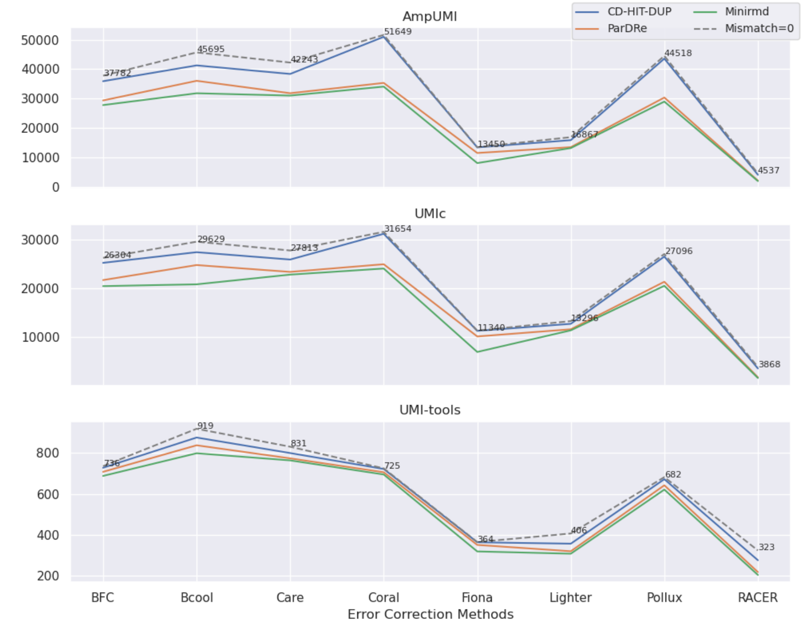


Supplementary Figure 43: Line charts comparing the overlapped read numbers in the deduplicated read set by the PCR-deduplication methods of CD-HIT-DUP, ParDRe, and Minirmd on the error-corrected dataset SRR1543967, with each UMI-based PCR-deduplication methods of UMI-tools, AmpUMI, and UMIc on dataset SRR1543967. Error correction was performed using error-correction methods of BFC, Bcool, Care, Coral, Fiona, Lighter, Pollux, and RACER, respectively. CD-HIT-DUP, ParDRe, and Minirmd employed a mismatched number set to 2. The dashed line labelled 'Mismatch=0' represents results obtained by CD-HIT-DUP with a mismatch setting of 0.

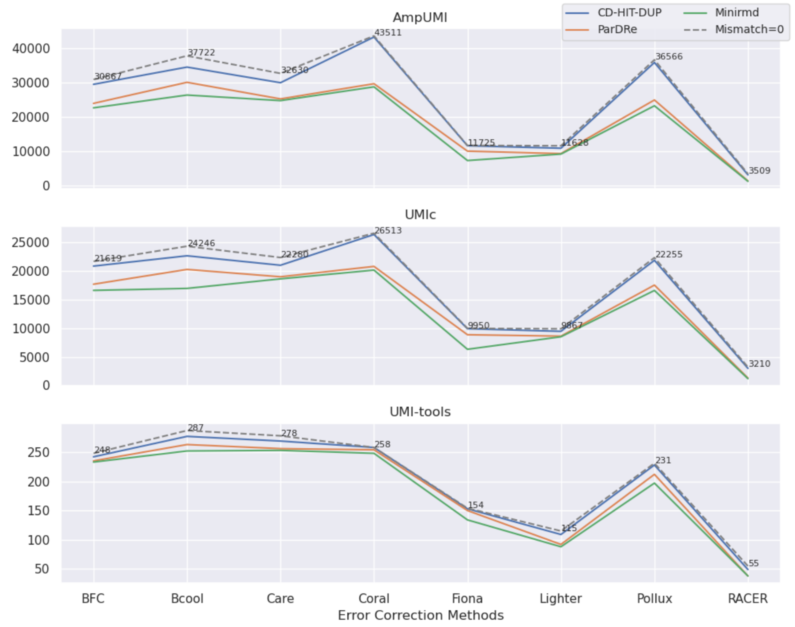


Supplementary Figure 44: Line charts comparing the overlapped read numbers in the deduplicated read set by the PCR-deduplication methods of CD-HIT-DUP, ParDRe, and Minirmd on the error-corrected dataset SRR1543968, with each UMI-based PCR-deduplication methods of UMI-tools, AmpUMI, and UMIc on dataset SRR1543968. Error correction was performed using error-correction methods of BFC, Bcool, Care, Coral, Fiona, Lighter, Pollux, and RACER, respectively. CD-HIT-DUP, ParDRe, and Minirmd employed a mismatched number set to 2. The dashed line labelled 'Mismatch=0' represents results obtained by CD-HIT-DUP with a mismatch setting of 0.

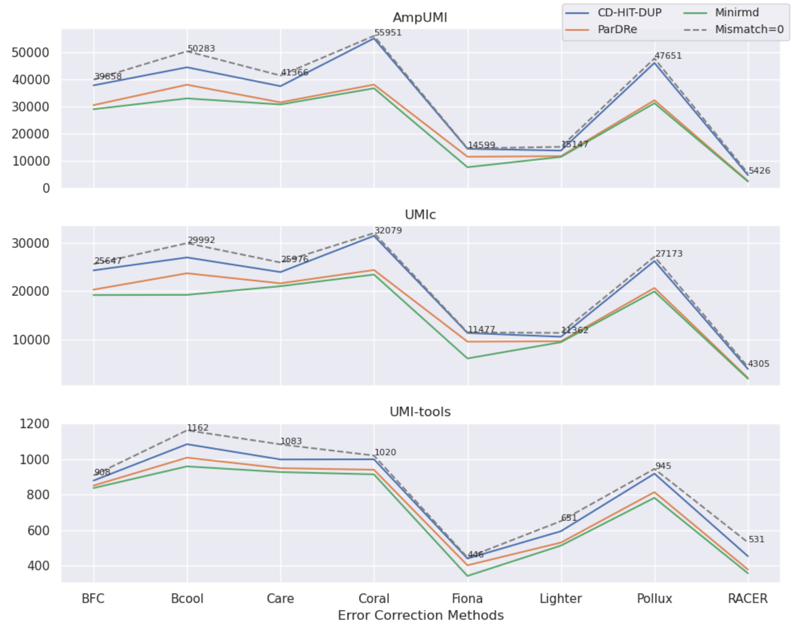


Supplementary Figure 45: Line charts comparing the overlapped read numbers in the deduplicated read set by the PCR-deduplication methods of CD-HIT-DUP, ParDRe, and Minirmd on the error-corrected dataset SRR1543969, with each UMI-based PCR-deduplication methods of UMI-tools, AmpUMI, and UMIc on dataset SRR1543969. Error correction was performed using error-correction methods of BFC, Bcool, Care, Coral, Fiona, Lighter, Pollux, and RACER, respectively. CD-HIT-DUP, ParDRe, and Minirmd employed a mismatched number set to 2. The dashed line labelled 'Mismatch=0' represents results obtained by CD-HIT-DUP with a mismatch setting of 0.

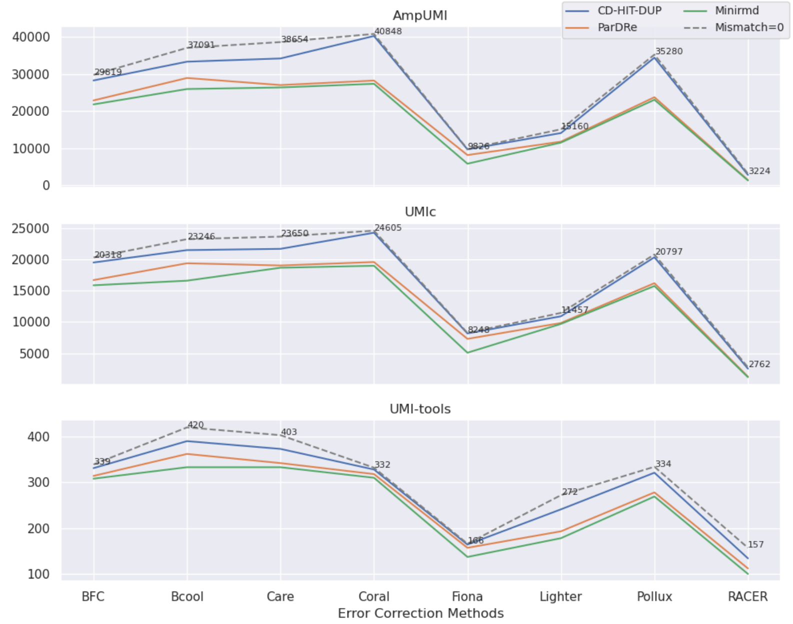


Supplementary Figure 46: Line charts comparing the overlapped read numbers in the deduplicated read set by the PCR-deduplication methods of CD-HIT-DUP, ParDRe, and Minirmd on the error-corrected dataset SRR1543970, with each UMI-based PCR-deduplication methods of UMI-tools, AmpUMI, and UMIc on dataset SRR1543970. Error correction was performed using error-correction methods of BFC, Bcool, Care, Coral, Fiona, Lighter, Pollux, and RACER, respectively. CD-HIT-DUP, ParDRe, and Minirmd employed a mismatched number set to 2. The dashed line labelled 'Mismatch=0' represents results obtained by CD-HIT-DUP with a mismatch setting of 0.

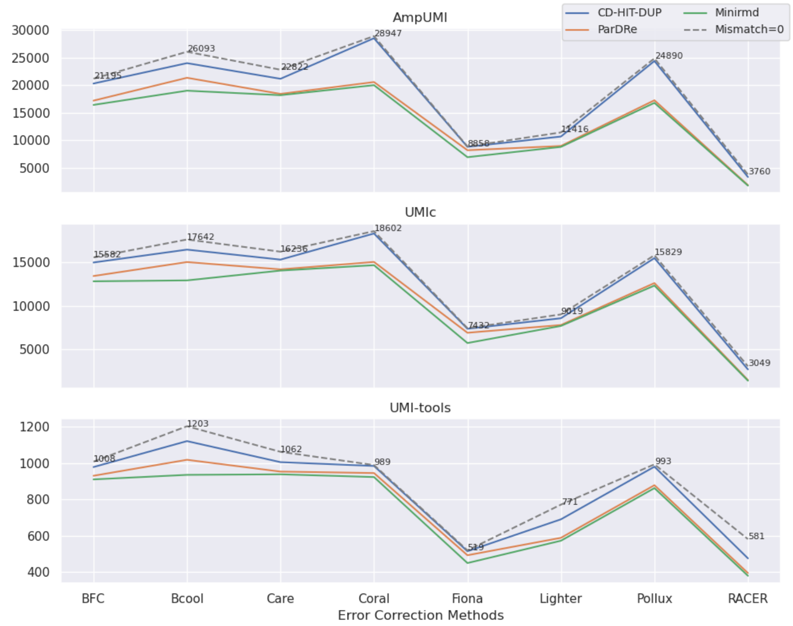


Supplementary Figure 47: Line charts comparing the overlapped read numbers in the deduplicated read set by the PCR-deduplication methods of CD-HIT-DUP, ParDRe, and Minirmd on the error-corrected dataset SRR1543971, with each UMI-based PCR-deduplication methods of UMI-tools, AmpUMI, and UMIc on dataset SRR1543971. Error correction was performed using error-correction methods of BFC, Bcool, Care, Coral, Fiona, Lighter, Pollux, and RACER, respectively. CD-HIT-DUP, ParDRe, and Minirmd employed a mismatched number set to 2. The dashed line labelled 'Mismatch=0' represents results obtained by CD-HIT-DUP with a mismatch setting of 0.

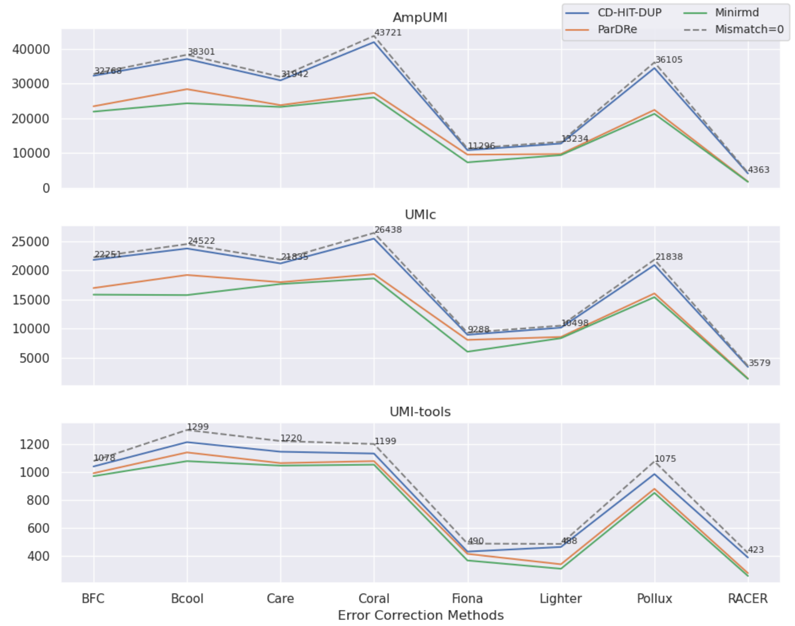


Supplementary Figure 48: Line charts comparing the overlapped read numbers in the deduplicated read set by the PCR-deduplication methods of CD-HIT-DUP, ParDRe, and Minirmd on the error-corrected dataset SRR1543965, with each UMI-based PCR-deduplication methods of UMI-tools, AmpUMI, and UMIc on dataset SRR1543965. Error correction was performed using error-correction methods of BFC, Bcool, Care, Coral, Fiona, Lighter, Pollux, and RACER, respectively. CD-HIT-DUP, ParDRe, and Minirmd employed a mismatched number set to 3. The dashed line labelled ‘Mismatch=0’ represents results obtained by CD-HIT-DUP with a mismatch setting of 0.

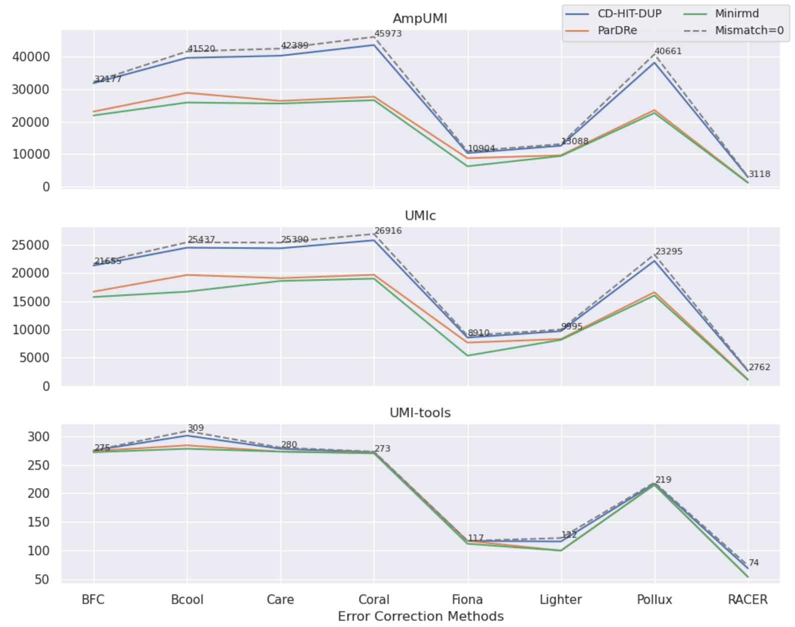


Supplementary Figure 49: Line charts comparing the overlapped read numbers in the deduplicated read set by the PCR-deduplication methods of CD-HIT-DUP, ParDRe, and Minirmd on the error-corrected dataset SRR1543966, with each UMI-based PCR-deduplication methods of UMI-tools, AmpUMI, and UMIc on dataset SRR1543966. Error correction was performed using error-correction methods of BFC, Bcool, Care, Coral, Fiona, Lighter, Pollux, and RACER, respectively. CD-HIT-DUP, ParDRe, and Minirmd employed a mismatched number set to 3. The dashed line labelled ‘Mismatch=0’ represents results obtained by CD-HIT-DUP with a mismatch setting of 0.

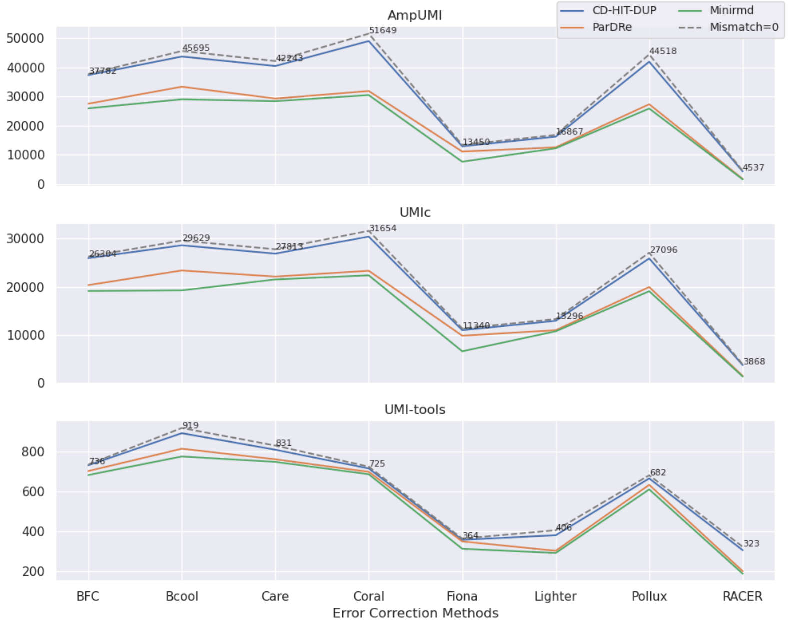


Supplementary Figure 50: Line charts comparing the overlapped read numbers in the deduplicated read set by the PCR-deduplication methods of CD-HIT-DUP, ParDRe, and Minirmd on the error-corrected dataset SRR1543967, with each UMI-based PCR-deduplication methods of UMI-tools, AmpUMI, and UMIc on dataset SRR1543967. Error correction was performed using error-correction methods of BFC, Bcool, Care, Coral, Fiona, Lighter, Pollux, and RACER, respectively. CD-HIT-DUP, ParDRe, and Minirmd employed a mismatched number set to 3. The dashed line labelled ‘Mismatch=0’ represents results obtained by CD-HIT-DUP with a mismatch setting of 0.

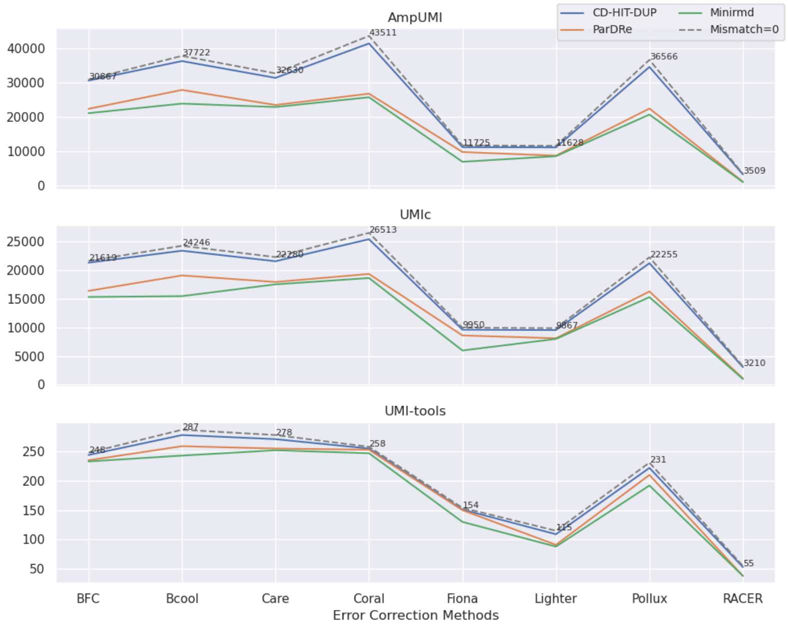


Supplementary Figure 51: Line charts comparing the overlapped read numbers in the deduplicated read set by the PCR-deduplication methods of CD-HIT-DUP, ParDRe, and Minirmd on the error-corrected dataset SRR1543968, with each UMI-based PCR-deduplication methods of UMI-tools, AmpUMI, and UMIc on dataset SRR1543968. Error correction was performed using error-correction methods of BFC, Bcool, Care, Coral, Fiona, Lighter, Pollux, and RACER, respectively. CD-HIT-DUP, ParDRe, and Minirmd employed a mismatched number set to 3. The dashed line labelled ‘Mismatch=0’ represents results obtained by CD-HIT-DUP with a mismatch setting of 0.

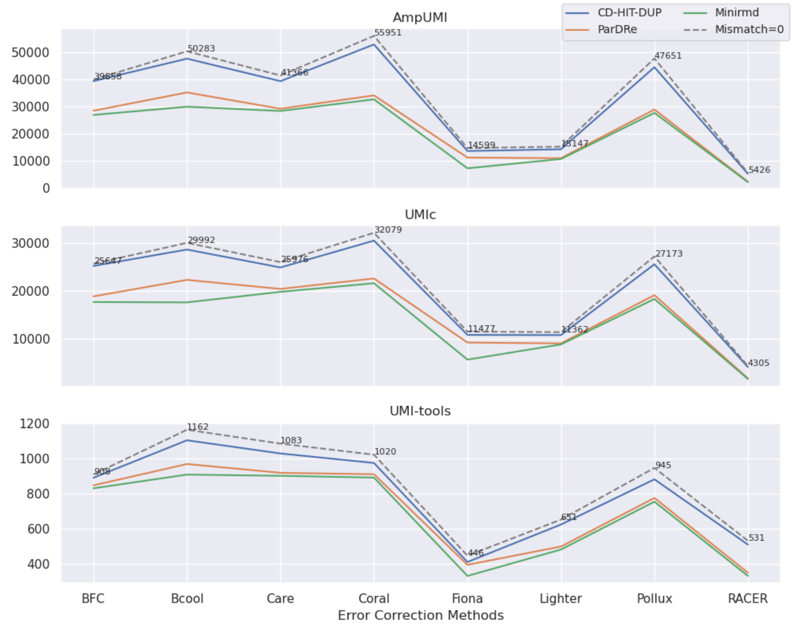


Supplementary Figure 52: Line charts comparing the overlapped read numbers in the deduplicated read set by the PCR-deduplication methods of CD-HIT-DUP, ParDRe, and Minirmd on the error-corrected dataset SRR1543969, with each UMI-based PCR-deduplication methods of UMI-tools, AmpUMI, and UMIc on dataset SRR1543969. Error correction was performed using error-correction methods of BFC, Bcool, Care, Coral, Fiona, Lighter, Pollux, and RACER, respectively. CD-HIT-DUP, ParDRe, and Minirmd employed a mismatched number set to 3. The dashed line labelled 'Mismatch=0' represents results obtained by CD-HIT-DUP with a mismatch setting of 0.

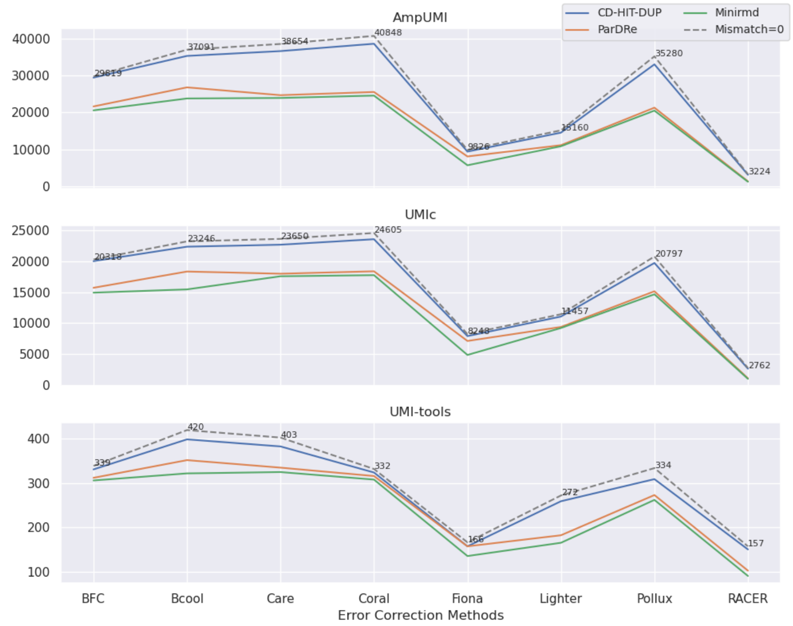


Supplementary Figure 53: Line charts comparing the overlapped read numbers in the deduplicated read set by the PCR-deduplication methods of CD-HIT-DUP, ParDRe, and Minirmd on the error-corrected dataset SRR1543970, with each UMI-based PCR-deduplication methods of UMI-tools, AmpUMI, and UMIc on dataset SRR1543970. Error correction was performed using error-correction methods of BFC, Bcool, Care, Coral, Fiona, Lighter, Pollux, and RACER, respectively. CD-HIT-DUP, ParDRe, and Minirmd employed a mismatched number set to 3. The dashed line labelled 'Mismatch=0' represents results obtained by CD-HIT-DUP with a mismatch setting of 0.

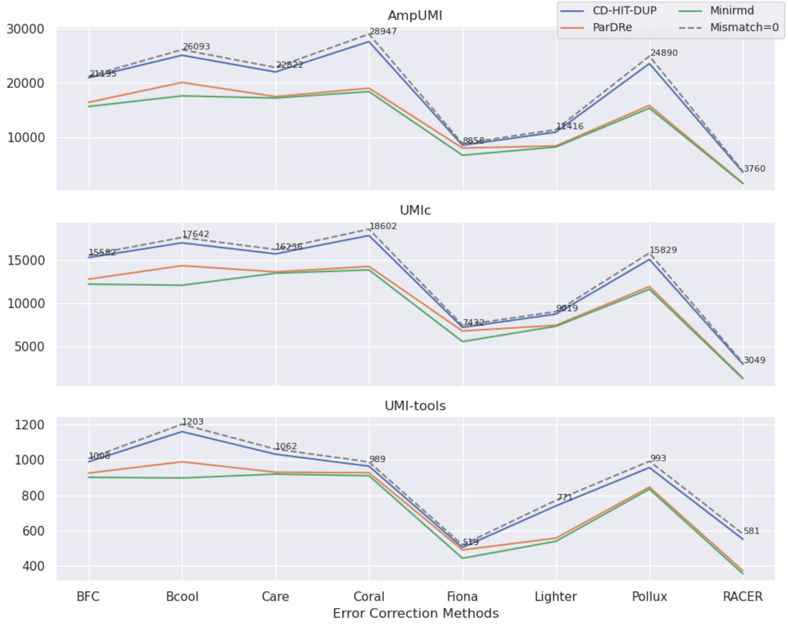


Supplementary Figure 54: Line charts comparing the overlapped read numbers in the deduplicated read set by the PCR-deduplication methods of CD-HIT-DUP, ParDRe, and Minirmd on the error-corrected dataset SRR1543971, with each UMI-based PCR-deduplication methods of UMI-tools, AmpUMI, and UMIc on dataset SRR1543971. Error correction was performed using error-correction methods of BFC, Bcool, Care, Coral, Fiona, Lighter, Pollux, and RACER, respectively. CD-HIT-DUP, ParDRe, and Minirmd employed a mismatched number set to 3. The dashed line labelled 'Mismatch=0' represents results obtained by CD-HIT-DUP with a mismatch setting of 0.

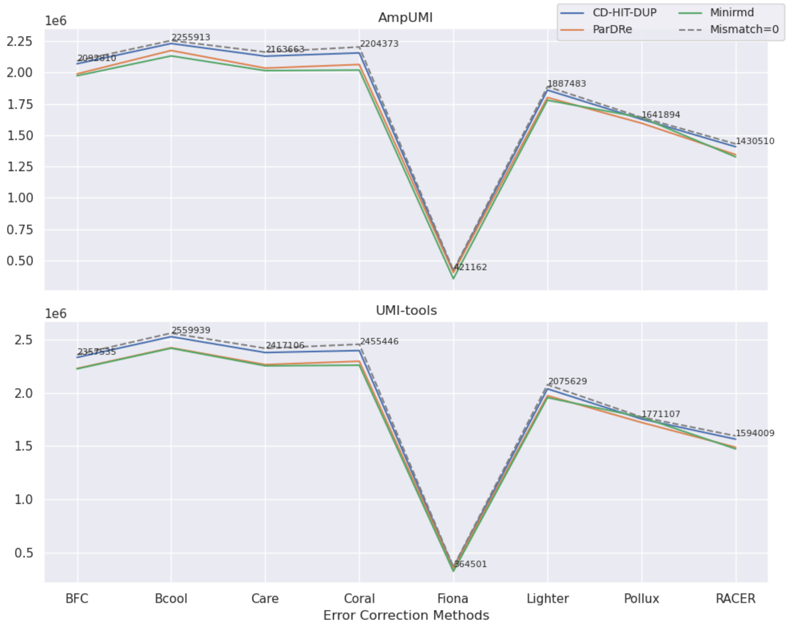


Supplementary Figure 55: Line charts comparing the overlapped read numbers in the deduplicated read set by the PCR-deduplication methods of CD-HIT-DUP, ParDRe, and Minirmd on the error-corrected dataset SRR28313972, with each UMI-based PCR-deduplication methods of UMI-tools and AmpUMI on dataset SRR28313972. Error correction was performed using error-correction methods of BFC, Bcool, Care, Coral, Fiona, Lighter, Pollux, and RACER, respectively. CD-HIT-DUP, ParDRe, and Minirmd employed a mismatched number set to 1. The dashed line labelled 'Mismatch=0' represents results obtained by CD-HIT-DUP with a mismatch setting of 0.

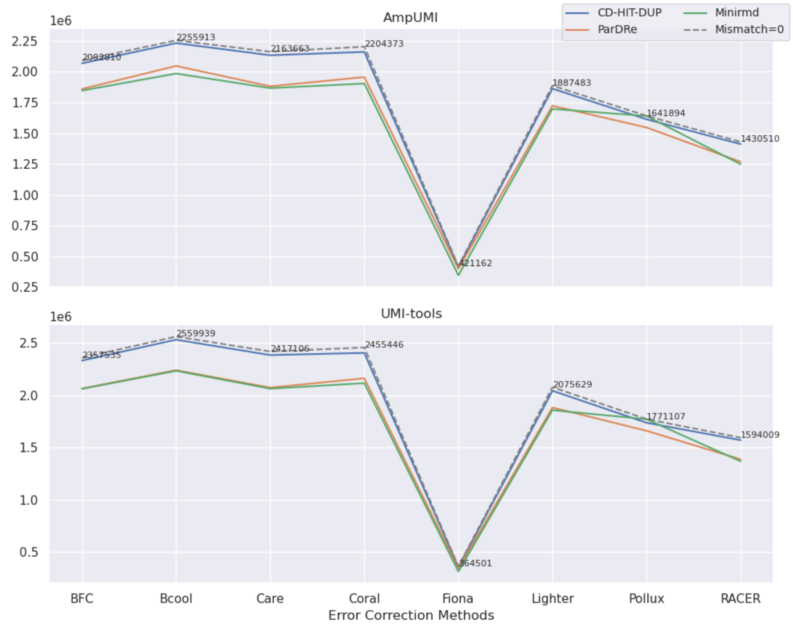


Supplementary Figure 56: Line charts comparing the overlapped read numbers in the deduplicated read set by the PCR-deduplication methods of CD-HIT-DUP, ParDRe, and Minirmd on the error-corrected dataset SRR28313972, with each UMI-based PCR-deduplication methods of UMI-tools and AmpUMI on dataset SRR28313972. Error correction was performed using error-correction methods of BFC, Bcool, Care, Coral, Fiona, Lighter, Pollux, and RACER, respectively. CD-HIT-DUP, ParDRe, and Minirmd employed a mismatched number set to 2. The dashed line labelled ‘Mismatch=0’ represents results obtained by CD-HIT-DUP with a mismatch setting of 0.

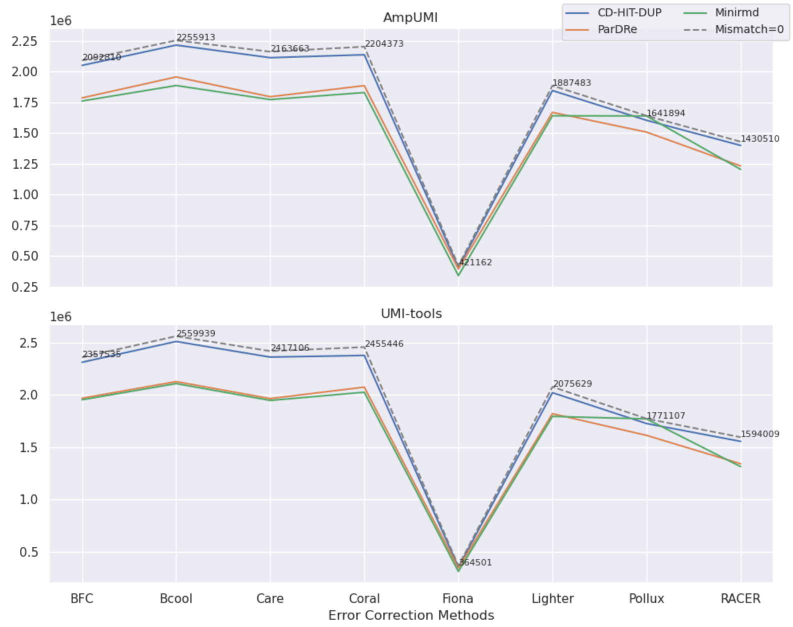


Supplementary Figure 57: Line charts comparing the overlapped read numbers in the deduplicated read set by the PCR-deduplication methods of CD-HIT-DUP, ParDRe, and Minirmd on the error-corrected dataset SRR28313972, with each UMI-based PCR-deduplication methods of UMI-tools and AmpUMI on dataset SRR28313972. Error correction was performed using error-correction methods of BFC, Bcool, Care, Coral, Fiona, Lighter, Pollux, and RACER, respectively. CD-HIT-DUP, ParDRe, and Minirmd employed a mismatched number set to 3. The dashed line labelled ‘Mismatch=0’ represents results obtained by CD-HIT-DUP with a mismatch setting of 0.

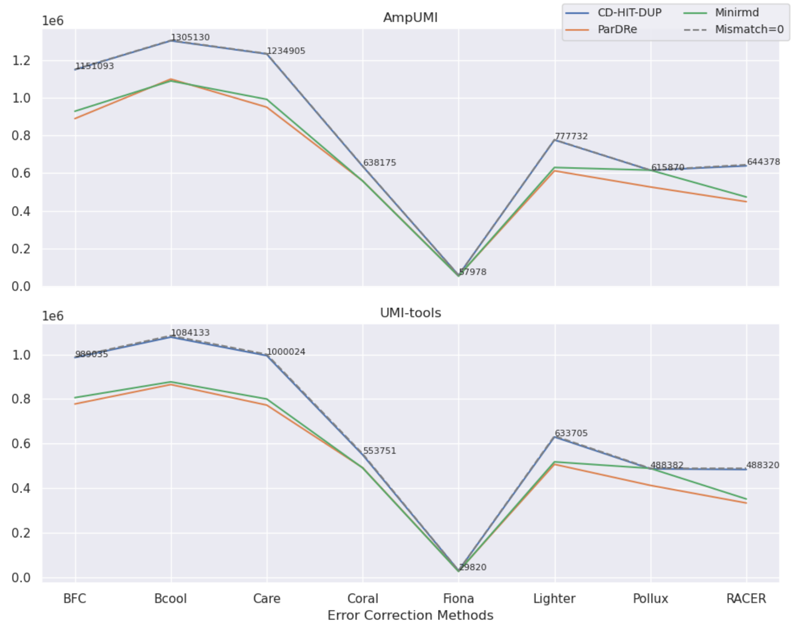


Supplementary Figure 58: Line charts comparing the overlapped read numbers in the deduplicated read set by the PCR-deduplication methods of CD-HIT-DUP, ParDRe, and Minirmd on the error-corrected dataset SRR28313990, with each UMI-based PCR-deduplication methods of UMI-tools and AmpUMI on dataset SRR28313990. Error correction was performed using error-correction methods of BFC, Bcool, Care, Coral, Fiona, Lighter, Pollux, and RACER, respectively. CD-HIT-DUP, ParDRe, and Minirmd employed a mismatched number set to 1. The dashed line labelled ‘Mismatch=0’ represents results obtained by CD-HIT-DUP with a mismatch setting of 0.

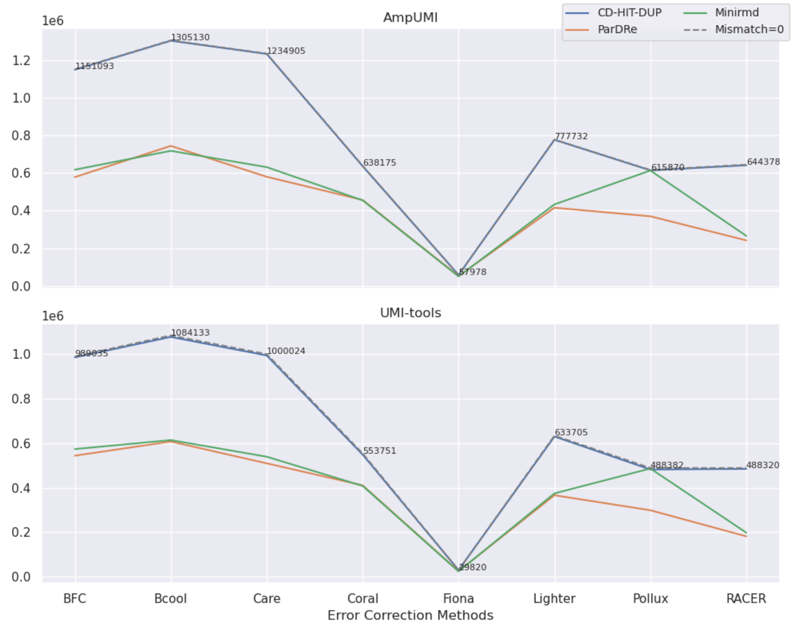


Supplementary Figure 59: Line charts comparing the overlapped read numbers in the deduplicated read set by the PCR-deduplication methods of CD-HIT-DUP, ParDRe, and Minirmd on the error-corrected dataset SRR28313990, with each UMI-based PCR-deduplication methods of UMI-tools and AmpUMI on dataset SRR28313990. Error correction was performed using error-correction methods of BFC, Bcool, Care, Coral, Fiona, Lighter, Pollux, and RACER, respectively. CD-HIT-DUP, ParDRe, and Minirmd employed a mismatched number set to 2. The dashed line labelled ‘Mismatch=0’ represents results obtained by CD-HIT-DUP with a mismatch setting of 0.

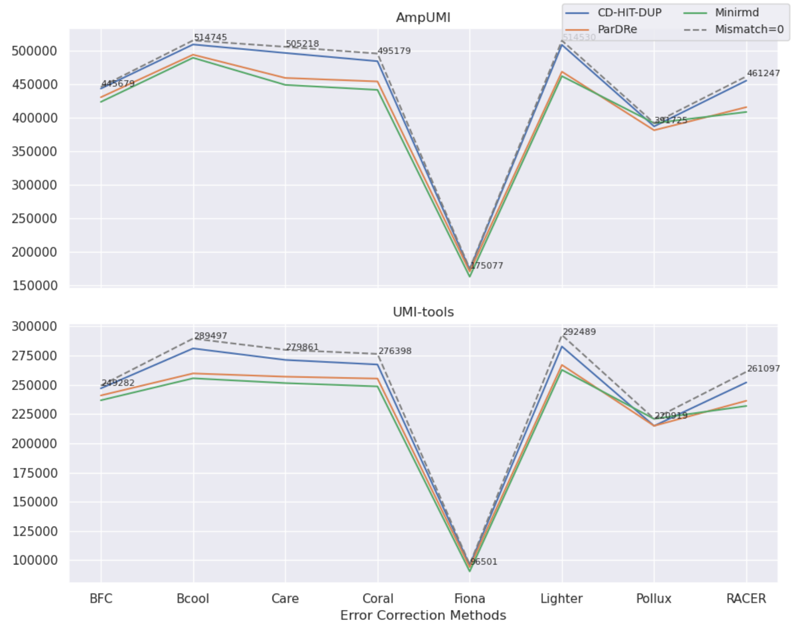


Supplementary Figure 60: Line charts comparing the overlapped read numbers in the deduplicated read set by the PCR-deduplication methods of CD-HIT-DUP, ParDRe, and Minirmd on the error-corrected dataset SRR28314008, with each UMI-based PCR-deduplication methods of UMI-tools and AmpUMI on dataset SRR28314008. Error correction was performed using error-correction methods of BFC, Bcool, Care, Coral, Fiona, Lighter, Pollux, and RACER, respectively. CD-HIT-DUP, ParDRe, and Minirmd employed a mismatched number set to 1. The dashed line labelled ‘Mismatch=0’ represents results obtained by CD-HIT-DUP with a mismatch setting of 0.

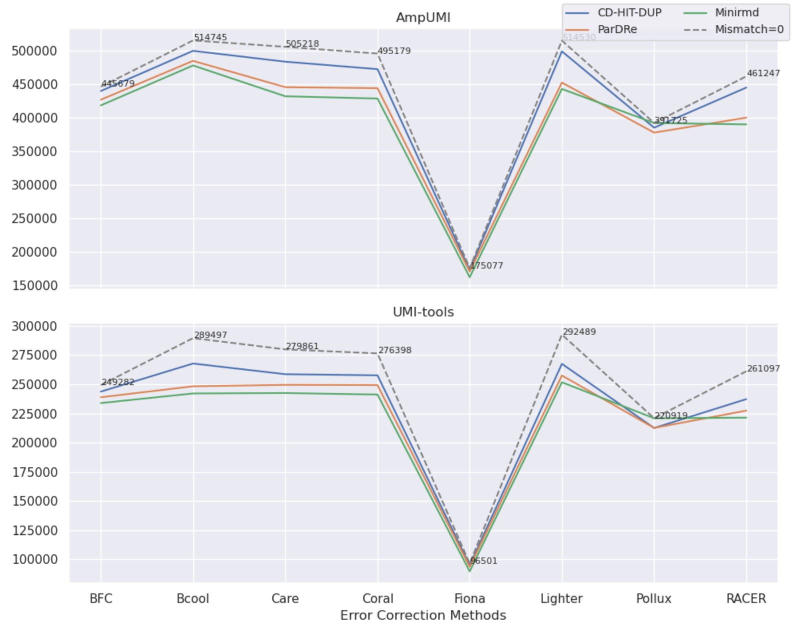


Supplementary Figure 61: Line charts comparing the overlapped read numbers in the deduplicated read set by the PCR-deduplication methods of CD-HIT-DUP, ParDRe, and Minirmd on the error-corrected dataset SRR28314008, with each UMI-based PCR-deduplication methods of UMI-tools and AmpUMI on dataset SRR28314008. Error correction was performed using error-correction methods of BFC, Bcool, Care, Coral, Fiona, Lighter, Pollux, and RACER, respectively. CD-HIT-DUP, ParDRe, and Minirmd employed a mismatched number set to 2. The dashed line labelled ‘Mismatch=0’ represents results obtained by CD-HIT-DUP with a mismatch setting of 0.

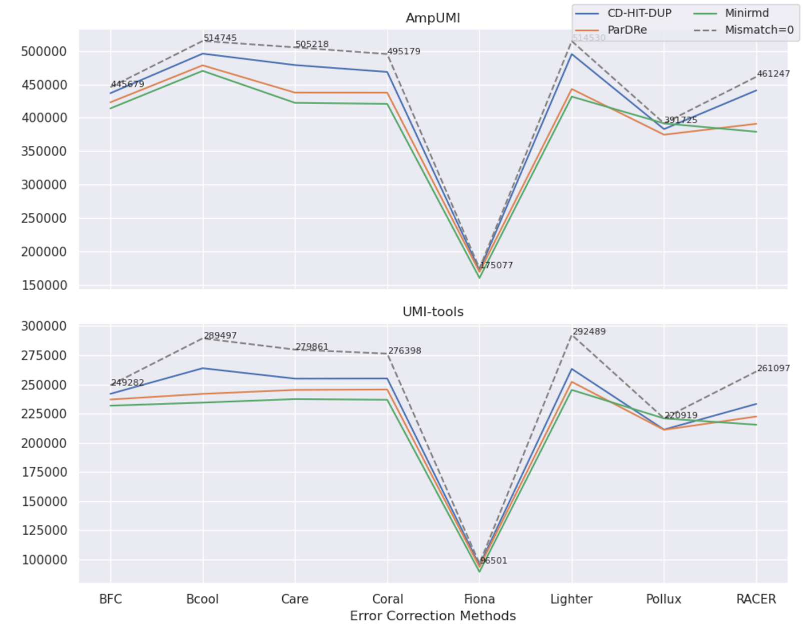


Supplementary Figure 62: Line charts comparing the overlapped read numbers in the deduplicated read set by the PCR-deduplication methods of CD-HIT-DUP, ParDRe, and Minirmd on the error-corrected dataset SRR28314008, with each UMI-based PCR-deduplication methods of UMI-tools and AmpUMI on dataset SRR28314008. Error correction was performed using error-correction methods of BFC, Bcool, Care, Coral, Fiona, Lighter, Pollux, and RACER, respectively. CD-HIT-DUP, ParDRe, and Minirmd employed a mismatched number set to 3. The dashed line labelled 'Mismatch=0' represents results obtained by CD-HIT-DUP with a mismatch setting of 0.

### 2 Supplementary Tables

Supplementary Tables 1-2 illustrate Summary of changes in unique reads, corrected reads, and erroneously introduced new reads after error correction using the error-correction methods of BFC, Bcool, Care, Coral, Fiona, Lighter, Pollux and RACER on the data sets of SRR1543965-SRR1543971, SRR28313990 and SRR28314008.

Supplementary Table 1: Summary of changes in unique reads, corrected reads, and erroneously introduced new reads after error correction using the error-correction methods of BFC, Bcool, Care, Coral, Fiona, Lighter, Pollux and RACER on the datasets SRR1543965-SRR1543969.

| Datasets | Methods | The number of unique reads | | | Corrected | Total | Corrected Percentage | New Reads Number |
|---|---|---|---|---|---|---|---|---|
| | | before correction | after correction | decreased by | reads number | | | |
| SRR1543965 | Bcool | 273707 | 225375 | 17.66% | 77398 | 952554 | 8.13% | 13430 |
| | BFC | | 272329 | 0.50% | 96237 | | 10.10% | 89155 |
| | Care | | 191284 | 30.11% | 95473 | | 10.02% | 8707 |
| | Coral | | 247341 | 9.63% | 37472 | | 3.93% | 4534 |
| | Fiona | | 169730 | 37.99% | 327099 | | 34.34% | 107473 |
| | Lighter | | 174343 | 36.30% | 433875 | | 45.55% | 118246 |
| | Pollux | | 237835 | 13.11% | 132427 | | 13.90% | 31953 |
| | RACER | | 250580 | 8.45% | 825232 | | 86.63% | 230539 |
| SRR1543966 | Bcool | 305104 | 260389 | 14.66% | 71497 | 1225804 | 5.83% | 16316 |
| | BFC | | 302654 | 0.80% | 120505 | | 9.83% | 103804 |
| | Care | | 267511 | 12.32% | 45436 | | 3.71% | 6206 |
| | Coral | | 271573 | 10.99% | 42835 | | 3.49% | 3057 |
| | Fiona | | 191718 | 37.16% | 291728 | | 23.80% | 122048 |
| | Lighter | | 193137 | 36.70% | 508163 | | 41.46% | 128164 |
| | Pollux | | 271472 | 11.02% | 149290 | | 12.18% | 28107 |
| | RACER | | 257834 | 15.49% | 1114938 | | 90.96% | 247740 |
| SRR1543967 | Bcool | 336566 | 280936 | 16.53% | 92368 | 1404275 | 6.58% | 17989 |
| | BFC | | 335031 | 0.46% | 129612 | | 9.23% | 116100 |
| | Care | | 266289 | 20.88% | 80193 | | 5.71% | 6978 |
| | Coral | | 303146 | 9.93% | 44945 | | 3.20% | 5196 |
| | Fiona | | 211267 | 37.23% | 344803 | | 24.55% | 137048 |
| | Lighter | | 217844 | 35.27% | 509769 | | 36.30% | 128122 |
| | Pollux | | 296749 | 11.83% | 173284 | | 12.34% | 29323 |
| | RACER | | 308299 | 8.40% | 1245191 | | 88.67% | 289427 |
| SRR1543968 | Bcool | 277589 | 227312 | 18.11% | 95326 | 1132736 | 8.42% | 13847 |
| | BFC | | 275656 | 0.70% | 114891 | | 10.14% | 105263 |
| | Care | | 198186 | 28.60% | 88941 | | 7.85% | 6611 |
| | Coral | | 251775 | 9.30% | 35155 | | 3.10% | 2468 |
| | Fiona | | 169764 | 38.84% | 299686 | | 26.46% | 105442 |
| | Lighter | | 162886 | 41.32% | 515265 | | 45.49% | 113154 |
| | Pollux | | 240844 | 13.24% | 151317 | | 13.36% | 24506 |
| | RACER | | 254251 | 8.41% | 1045490 | | 92.30% | 243957 |
| SRR1543969 | Bcool | 375335 | 325515 | 13.27% | 92941 | 1338638 | 6.94% | 30150 |
| | BFC | | 373174 | 0.58% | 158448 | | 11.84% | 143477 |
| | Care | | 263721 | 29.74% | 128400 | | 9.59% | 12058 |
| | Coral | | 331403 | 11.70% | 60184 | | 4.50% | 5044 |
| | Fiona | | 234396 | 37.55% | 377712 | | 28.22% | 152277 |
| | Lighter | | 224814 | 40.10% | 583947 | | 43.62% | 141116 |
| | Pollux | | 319236 | 14.95% | 195977 | | 14.64% | 31433 |
| | RACER | | 315614 | 15.91% | 1199132 | | 89.58% | 291860 |

Supplementary Table 2: Summary of changes in unique reads, corrected reads, and erroneously introduced new reads after error correction using the error-correction methods of BFC, Bcool, Care, Coral, Fiona, Lighter, Pollux and RACER on the datasets SRR1543970-SRR1543971, SRR28313990 and SRR28314008.

| Datasets | Methods | The number of unique reads | | | Corrected | Total | Corrected | New Reads |
|---|---|---|---|---|---|---|---|---|
| | | before correction | after correction | decreased by | reads number | | Percentage | Number |
| SRR1543970 | Bcool | 290058 | 238083 | 17.92% | 80100 | 1219666 | 6.57% | 14812 |
| | BFC | | 287510 | 0.88% | 118777 | | 9.74% | 105239 |
| | Care | | 252010 | 13.12% | 46465 | | 3.81% | 6768 |
| | Coral | | 250633 | 13.59% | 48603 | | 3.98% | 3098 |
| | Fiona | | 171523 | 40.87% | 292715 | | 24.00% | 118053 |
| | Lighter | | 186026 | 35.87% | 430315 | | 35.28% | 95734 |
| | Pollux | | 252119 | 13.08% | 134531 | | 11.03% | 28292 |
| | RACER | | 257492 | 11.23% | 1106449 | | 90.72% | 245119 |
| SRR1543971 | Bcool | 182485 | 146958 | 19.47% | 50130 | 746724 | 6.71% | 7379 |
| | BFC | | 181088 | 0.77% | 76835 | | 10.29% | 70518 |
| | Care | | 129353 | 29.12% | 61209 | | 8.20% | 5329 |
| | Coral | | 157737 | 13.56% | 34687 | | 4.65% | 4064 |
| | Fiona | | 111720 | 38.78% | 231785 | | 31.04% | 65538 |
| | Lighter | | 125096 | 31.45% | 265978 | | 35.62% | 76783 |
| | Pollux | | 158233 | 13.29% | 94785 | | 12.69% | 19978 |
| | RACER | | 189741 | -3.98% | 637304 | | 85.35% | 173945 |
| SRR28313990 | Bcool | 2669507 | 2587568 | 3.07% | 205105 | 5943780 | 3.45% | 4599 |
| | Care | | 2499211 | 6.38% | 320028 | | 5.38% | 141238 |
| | Coral | | 1614311 | 39.53% | 1572630 | | 26.46% | 303701 |
| | Fiona | | 1691600 | 36.63% | 4054483 | | 68.21% | 1613490 |
| | Lighter | | 2457358 | 7.95% | 1439218 | | 24.21% | 852568 |
| | Pollux | | 1668321 | 37.50% | 1811727 | | 30.48% | 407231 |
| | RACER | | 2953113 | -10.62% | 2191865 | | 36.88% | 1723226 |
| | BFC | | 2678966 | -0.35% | 699713 | | 11.77% | 603516 |
| SRR28314008 | Bcool | 1696037 | 1272751 | 24.96% | 673892 | 7563556 | 8.91% | 24329 |
| | Care | | 1249364 | 26.34% | 603932 | | 7.98% | 106565 |
| | Coral | | 1179827 | 30.44% | 743128 | | 9.83% | 132303 |
| | Fiona | | 774328 | 54.34% | 3325931 | | 43.97% | 547459 |
| | Lighter | | 1324479 | 21.91% | 653864 | | 8.64% | 46386 |
| | Pollux | | 826139 | 51.29% | 1435655 | | 18.98% | 238401 |
| | RACER | | 1417912 | 16.40% | 948832 | | 12.54% | 261460 |
| | BFC | | 1678715 | 1.02% | 1149440 | | 15.20% | 964572 |
